# Cross-covariance isolate detect: a new change-point method for estimating dynamic functional connectivity

**DOI:** 10.1101/2020.12.20.423696

**Authors:** Andreas Anastasiou, Ivor Cribben, Piotr Fryzlewicz

**Author notes:** Joint first authors (Andreas Anastasiou, Ivor Cribben).

## Abstract

Evidence of the non stationary behavior of functional connectivity (FC) networks has been observed in task based functional magnetic resonance imaging (fMRI) experiments and even prominently in resting state fMRI data. This has led to the development of several new statistical methods for estimating this time-varying connectivity, with the majority of the methods utilizing a sliding window approach. While computationally feasible, the sliding window approach has several limitations. In this paper, we circumvent the sliding window, by introducing a statistical method that finds change-points in FC networks where the number and location of change-points are unknown a priori. The new method, called *cross-covariance isolate detect* (CCID), detects multiple change-points in the second-order (cross-covariance or network) structure of multivariate, possibly high-dimensional time series. CCID allows for change-point detection in the presence of frequent changes of possibly small magnitudes, can assign change-points to one or multiple brain regions, and is computationally fast. In addition, CCID is particularly suited to task based data, where the subject alternates between task and rest, as it firstly attempts isolation of each of the change-points within subintervals, and secondly their detection therein. Furthermore, we also propose a new information criterion for CCID to identify the change-points. We apply CCID to several simulated data sets and to task based and resting state fMRI data and compare it to recent change-point methods. CCID may also be applicable to electroencephalography (EEG), magentoencephalography (MEG) and electrocorticography (ECoG) data. Similar to other biological networks, understanding the complex network organization and functional dynamics of the brain can lead to profound clinical implications. Finally, the R package **ccid** implementing the method from the paper is available from CRAN.

## 1. Introduction

Functional connectivity (FC) is the undirected association between two or more functional magnetic resonance imaging (fMRI) time series. Evidence of the non stationary behavior of FC (or FC networks) has been observed in high temporal resolution electroencephalography (EEG) data, task based fMRI experiments (Fox et al., 2005; Debener et al., 2006; Cribben et al., 2012, 2013) and even prominently in resting state fMRI data (Delamillieure et al., 2010; Doucet et al., 2012; Cribben & Yu, 2017; Ofori-Boateng et al., 2020). By estimating a FC network across the entire experimental time course, the resulting FC network is simply an average of the changing connectivity structures. While this is convenient for estimation and computation purposes, as it keeps the FC network estimation from becoming too complex, it presents a simplified version of a highly integrated and dynamic phenomenon.

In order to estimate this time varying phenomenon (or dynamic FC as it is widely known), many research papers first considered a sliding (or moving) window. These approaches begin at the first time point, then a block of a fixed number of time points (the window) are selected and all the data points within the block are used to estimate the FC. The window is then shifted a certain number of time points (overlapping or non-overlapping) and the FC is estimated on the new data. By shifting the window to the end of the experimental time course, researchers can estimate the dynamic FC. Chang & Glover (2010), Kiviniemi et al. (2011), Hutchison et al. (2013b) and Leonardi et al. (2013) investigated the dynamic functional FC in resting-state data using a moving-window approach, based on a time-frequency coherence analysis with wavelet transforms, an independent component analysis, a correlation analysis, and a principal component analysis, respectively. Allen et al. (2014), Handwerker et al. (2012), Jones et al. (2012) and Sakoğlu et al. (2010) considered a group independent component analysis (Calhoun et al., 2001) to decompose multi-subject resting-state data into functional brain regions, and a moving-window and *k*-means clustering of the windowed correlation matrices to study whole brain dynamic FC networks. While the sliding window approach is computationally feasible, it also has limitations (Hutchison et al., 2013a). For example, the choice of window size is crucial and sensitive, as different window sizes can lead to quite different FC patterns. Another disadvantage is that equal weight is given to all *k* neighbouring time points and 0 weight to all the others (Lindquist et al., 2014). Other methods have also been proposed that do away with the sliding window. For example, Zhang et al. (2014) proposed the dynamic Bayesian variable partition model that estimates and models multivariate dynamic functional interactions using a unified Bayesian framework.

Change-point methods have also been considered. There exists an extensive literature and a long history on change-points. The most widely discussed problems have been concerned with finding multiple change-points in univariate time series (Inclan & Tiao, 1994; Chen & Gupta, 1997). Recently, the multiple change-point detection problem in multivariate time series has received some attention especially in non-stationary practical problems (Fan et al., 2011). High dimensional time series change-point detection problems are the obvious but by no means straightforward extension of the univariate case. To detect changes in the covariance matrix of a multivariate time series, Aue et al. (2009) introduced a method using a nonparametric CUSUM type test, Dette & Wied (2016) proposed a test where the dimension of the data is fixed while Kao et al. (2018) considered the case where the dimension of the data increases with the sample size (they also investigated change-point analysis based on Principal Component Analysis). Sundararajan & Pourahmadi (2018) proposed a new method for detecting multiple change-points in the covariance structure of a multivariate piecewise-stationary process.

In other related work, Barnett & Onnela (2016) considered a method for detecting change-points in correlation networks that, unlike previous change-point detection methods designed for time series data, requires minimal distributional assumptions. Gibberd & Nelson (2014) studied the consistency properties of a regularised estimator for the simultaneous identification of both change-points and the graphical dependency structure in multivariate time series. The first comprehensive treatment of high dimensional time series factor models with multiple change-points in their second–order structure is put forward by Barigozzi et al. (2018). Li et al. (2019) considered multiple structural breaks in large contemporaneous covariance matrices of high dimensional time series satisfying an approximate factor model. Cho & Fryzlewicz (2015) segmented the multivariate time series into partitions based on the second-order structure. Within neuroscience, Cribben et al. (2012, 2013) put forward the Dynamic Connectivity Regression method for detecting multiple changepoints in the precision matrices (or undirected graph) from a multivariate time series. Schröder & Ombao (2019) introduced FreSpeD that employed a multivariate cumulative sum (CUSUM)-type procedure to detect change-points in autospectra and coherences for multivariate time series. Their method allows for the segmentation of the multivariate time series but also for the direct interpretation of the change in the sense that the change-point can be assigned to one or multiple time series (or Electroencephalogram channels) and frequency bands. Kirch et al. (2015) considered the At Most One Change setting and the epidemic setting (two change-points, where the process reverts back to the original regime after the second change-point) and provide some theoretical results. Cribben & Yu (2017) introduced the Network Change Point Detection method that uses an eigen-space based statistic for testing the community structures changes in stochastic block model sequences. More recently, Kundu et al. (2018) proposed a fully automated two-stage approach which pools information across multiple subjects to estimate change-points in functional connectivity, while Dai et al. (2019) developed a formal statistical test for finding change-points in time series associated with FC. Finally, Ofori-Boateng et al. (2020) introduced a new method that firstly presents each network snapshot of fMRI data as a linear object and finds its respective univariate characterization via local and global network topological summaries and then adopts a change-point detection method for (weakly) dependent time series based on efficient scores.

All of these methods have limitations. The most obvious is that they all use binary segmentation (BS) to identify multiple change-points. BS, while computationally feasible, is not optimal as it searches for a single change-point at a time. Hence, BS struggles to find multiple change-points in task based fMRI experiments, where the subject alternates between two states, rest and task, as the network structure for any two segments in the data split by a change-point is very similar. Furthermore, it is difficult for BS to find frequent changes of possibly small magnitudes. In addition, many of the previous methods (Cribben et al., 2012, 2013; Schröder & Ombao, 2019; Kirch et al., 2015; Dai et al., 2019) can only consider a relatively low number (up to *p*=30) of time series from either the channels or brain regions. Hence, their ability to detect whole brain connectivity change-points is impeded. In order to include a large number of time series, other methods (Cribben & Yu, 2017; Ofori-Boateng et al., 2020; Ondrus et al., 2021) perform a dimension reduction technique, such as singular value decomposition or summarize the network using graph summary statistics, to the data to make the problem more computationally feasible, but may lose information from the network structure, and hence the change-points therein, in this process.

In this paper, we introduce a new method, called *cross-covariance isolate detect* (CCID), to detect multiple change-points in the second-order structure of multivariate, possibly high-dimensional time series. The second-order structure can also be referred to as the dependence structure, the cross-covariance structure or the network structure, between the time series. CCID begins by first converting the multivariate time series (or regions of interest time series) into local wavelet periodograms and cross-periodograms. To detect the change-points, CCID aggregates across the multivariate time series through the *L*_2_ or the *L*_∞_ metric and then uses a scaled CUSUM statistic to decide whether a candidate is indeed a change-point or not, based on whether the relevant scaled CUSUM value is larger than an appropriately chosen threshold. Furthermore, we propose a new information criterion for CCID as an alternative approach to the threshold method for the change-point detection problem. CCID has the following unique and important attributes. First, CCID uses the Isolate-Detect principle (Anastasiou & Fryzlewicz, 2021) to find the multiple change-points. CCID works by first isolating each of the change-points within subintervals and then secondly detecting them within these subintervals. For general data sets we consider in this work, we find that CCID performs as well as, if not better than, the previous methods that use BS. However, for our simulations, where the subject alternates between two states and for our task based fMRI experiment, CCID clearly outperforms the BS methods. This is due to CCID’s ability to isolate the change-points between tasks while BS looks to partition the data into two intervals which is particularly difficult given the similarity between any two partitions (for this particular data type) and the fact that BS tends not to handle frequent change-point scenarios well. Second, CCID has the ability to find change-points that are very close to one another, which provides more insight into functional dynamics of the brain. Third, after detecting the change-points, CCID can assign them to one or multiple brain regions. Fourth, CCID is computationally fast. Fifth, CCID may also be applicable to electroencephalography (EEG), magentoencephalography (MEG) and electrocorticography (ECoG) data. Finally, the R package **ccid** implementing the method from the paper is available from CRAN.

The remainder of this paper is organized as follows. We introduce our new method, CCID, in Section 2. We describe our simulated data and our fMRI data in Section 3. The performance of the proposed method on both the simulated and real data is detailed in Section 4. A discussion of the limitations of the method and future work is described in Section 5. Finally, we conclude in Section 6 and provide details on the R package **ccid** in Section 7.

## 2. Materials and methods

Let **X** ∈ ℝ^*T* ×*p*^ be a multivariate time series with *T* time points and *p* regions of interest (ROIs: or data sequences). We assume that each univariate time series within **X** follows a Gaussian distribution. Smith et al. (2011) conclude that the Gaussian assumption is valid on good-quality fMRI data. We show how cross-covariance isolate detect (CCID) can be adapted to detect multiple change-points in the second-order (or cross-covariance) structure of multivariate, possibly high-dimensional time series. The second-order structure can also be referred to as the dependence structure, the cross-covariance structure or the network structure, between the time series. CCID first constructs wavelet-based local periodogram and cross-periodogram sequences from **X**, to which we apply the isolate-detect principle (Anastasiou & Fryzlewicz, 2021) to find multiple change-points in the second order structure of **X**.

### 2.1. Wavelets

A wavelet is a wave-like oscillation. Its amplitude begins at zero, increases (decreases), and then decreases (increases) back to zero. In wavelet theory, the wavelet correlates with another signal if it is of similar frequency at the location overlapping with the support of the wavelet. As a mathematical tool, wavelets can be used to extract information from many different types of data. We use wavelets in this work to transform the problem of detecting change-points in the second-order structure of the process to the expectations of the wavelet cross-periodograms and to “whiten” the data. For a detailed review of wavelets we refer readers to Daubechies (1992) and Vidakovic (2009).

### 2.2. Locally Stationary Wavelet model

By way of introduction, we consider Haar wavelets (the simplest example of a wavelet system) which are defined as

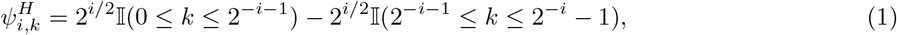

where *i* ∈ {−1, −2, …} and *k* ∈ ℤ denote scale and location parameters, respectively. Small negative values of the scale parameter *i* denote fine scales where the wavelet vectors are the most localized and oscillatory, while large negative values denote coarse scales with longer, less oscillatory wavelet vectors. With such wavelets as building blocks, we introduce the *p*-variate Locally Stationary Wavelet (LSW: Cho & Fryzlewicz, 2015) model as follows:

#### Definition 1

The *p*-variate LSW process 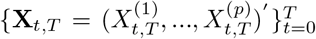 for *T* = 1, 2, …, is a triangular stochastic array with the following representation:

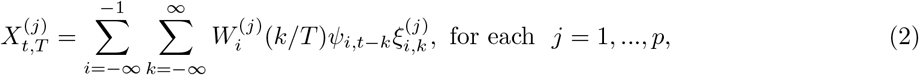

where 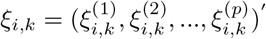 are independently generated from multivariate normal distributions *N*_*p*_{**0**, **Σ**_*i*_(*k/T*)}, with 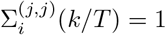 and

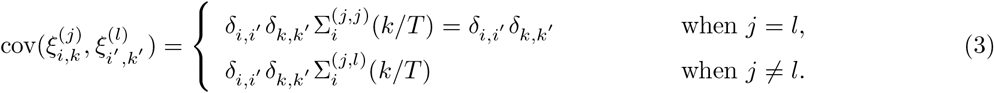

The Kronecker delta function *δ*_*i,i*′_ is 1 when *i* = *i*′ and 0 otherwise. For each *i* and *j, l* = 1, …, *p*, the functions 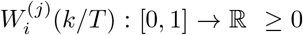 and 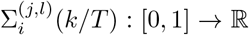 are piecewise-constant with an unknown number of change-points. The autocovariance and cross-covariance functions of **X**_*t,T*_ approximately inherit the piecewise-constant property of 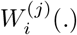 and 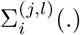 with almost identical change-point locations. See Cho & Fryzlewicz (2015) for details.

### 2.3. Wavelet periodograms and cross-periodograms

We now construct appropriate wavelet-based local periodogram sequences from the *p*-variate LSW time series **X**_*t,T*_. With the empirical wavelet coefficients at scale *i* denoted by 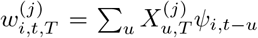 for each 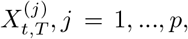 the wavelet periodogram of 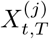 and the wavelet cross-periodogram between 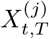 and 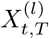 at scale *i* are defined as

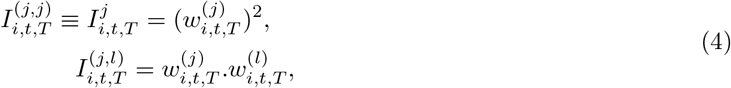

respectively. As 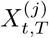 is Gaussian, 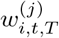 is also Gaussian, hence each 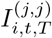 follows a scaled 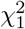 distribution.

For a multivariate LSW process **X**_*t,T*_, its theoretical (population) local autocovariance and cross-covariance functions are defined by

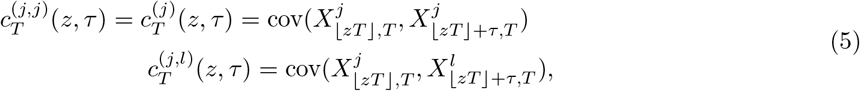

In the multivariate LSW model, Cho & Fryzlewicz (2015) showed that there is an asymptotic one-to-one correspondence between these theoretical local autocovariance and cross-covariance functions, the population wavelet periodograms and cross-periodograms, and the expectations of wavelet periodograms and cross-periodograms. Specifically, there exists an asymptotic one-to-one correspondence for any pair of 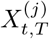 and 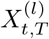 between the following quantities: the autocovariance and cross-covariance functions 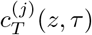 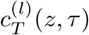 and 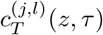 at lags *τ* = 0, 1, …, piecewise-constant functions 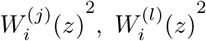 and 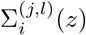 and the expectations of wavelet periodograms and cross-periodograms 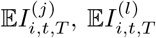 and 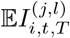 at scales *i* = −1, −2, …, Therefore, the change-points in the second order structure of the multivariate time series **X**_*t,T*_ can be detected from the wavelet periodograms and cross-periodograms at multiple scales.

We now focus on the wavelet periodogram 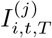 and the cross-periodogram 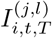, and use them as the inputs into the CCID algorithm. We firstly note that 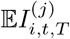 are piecewise-constant except for negligible biases around the change-points. Since 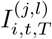 does not follow a multiplicative model, we consider instead the following alternative:

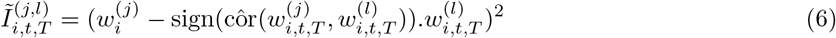

where côr is the sample correlation computed separately on each current segment (data between change-points). This is a multiplicative model but results in no loss of information about any change-points, see Cho & Fryzlewicz (2015).

There are four main advantages for using wavelets (the LSW model) in our CCID model for the purpose of change-point detection. First, there is one-to-one correspondence between the auto/cross-covariance structure of a time series, and its wavelet spectrum, or alternatively its (population) wavelet cross-periodograms. The (population) wavelet cross-periodogram representation, enabled by using the wavelet model, is the most convenient of the three for the purpose of change-point detection. The reason is that the natural estimator of the population wavelet (cross-)periodogram is the empirical wavelet (cross-)periodogram, which is a scaled *χ*^2^ distributed quantity under Gaussianity (and follows a general multiplicative model under other distributions). This makes it particularly suited for segmentation (=change-point detection) as a great deal is known in the literature (Inclan & Tiao, 1994; Cho & Fryzlewicz, 2015) about segmenting multiplicative-model data sequences of the form present in the empirical wavelet (cross-)periodograms. Second, a known difficult issue arising in time series segmentation is the presence of autocorrelation in the data, which can ‘fool’ change-point detectors as natural fluctuations in the time series due to autocorrelations can mask as change-points, and vice versa. The use of wavelets (we need to take a wavelet transform of the data in order to compute the empirical cross-periodograms) are helpful here due to their well-known whitening property (Vidakovic, 2009), whereby they tend to reduce positive autocorrelation in time series, hence making the change-point detection problem easier. Third, the whole array of the wavelet periodograms at all scales are easily and rapidly computable via the wavelet transform. Fourth, the normality assumption in LSW is reasonable for high quality fMRI data.

### 2.4. Model specification

With *p* being the dimensionality of the initial given time series, we assume the following multiplicative model for the periodograms 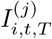 and cross periodogrames 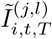 for *i* = 1, …, *p*, and *d* = *Jp*(*p*+1)/2 (assuming *J* scales), by transforming our data to periodograms and cross periodograms, which we denote as **Y**:

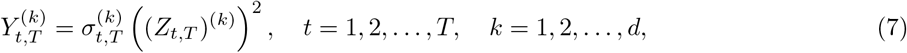

where 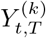 are the observed data related to the *k*^*th*^ data sequence, 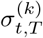 is a piecewise-constant mean function and 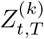 is a sequence of (possibly) auto-correlated standard normal random variables. This means that 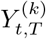 is a scaled 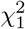-random variable, as the empirical wavelet periodogram is a squared zero-mean normal variables (Cho & Fryzlewicz, 2012), with 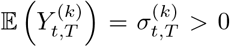. Our purpose is to detect the number and the location of changes in the mean of 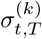, for any *k* ∈ {1, 2, …, *d*}, with each change-point being possibly shared by more than one data sequence. We only consider the finest scale of *i* = −1 used in the wavelet transformation as fMRI data typically only have lag 1 autocorrelation (Fiecas et al., 2017; Arbabshirani et al., 2014). Hence, *d* = *p*(*p* + 1)/2.

In order to detect the change-points we use the following statistic:

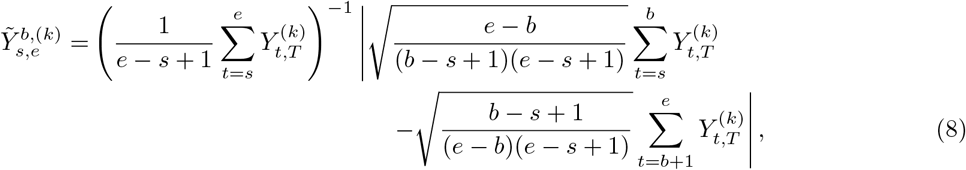

where 1 ≤ *s* ≤ *b* < *e* ≤ *T*. The statistic is a (scaled) CUSUM statistic, where we divide by the sample mean of the observations 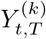 with *t* ∈ [*s, e*]. This is necessary in multiplicative settings such as (7) in order to make the results independent of the magnitude of 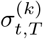. Essentially, the CUSUM statistic compares the means of the data to the left and to the right of each postulated change-point location *b*. To obtain a candidate change-point on any interval [*s, e*], we maximise the CUSUM statistic over *b*: this is where the (scaled) difference in means is the largest, so this will be the likeliest candidate for a change-point. In addition, we note that due to (7), the statistic in (8) is positive. More information on (8) can be found in Inclan & Tiao (1994) and Cho & Fryzlewicz (2015).

### 2.5. CCID

Our new method, CCID, proceeds as follows. We describe the Isolate-Detect (Anastasiou & Fryzlewicz, 2021) segmentation algorithm illustrated in Figure 1. The basic idea is that for the *d* observed data sequences of length *T* each, and with *λ*_*T*_ a suitably chosen positive integer, Isolate-Detect (ID) first creates two ordered sets of *K* = [*T/λ*_*T*_] right- and left-expanding intervals. We collect these intervals in the ordered set *S*_*RL*_ = {*R*_1_, *L*_1_, *R*_2_, *L*_2_, …, *R*_*K*_, *L*_*K*_}. Then, for each point *b* in the interval *R*_1_ and for each *k* = 1, 2 …, *d*, we calculate the values of 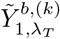 as in equation (8). As already mentioned, we aggregate the information of each of the *d* data sequences using either the *L*_2_ or the *L*_∞_ metric. For the *L*_2_ approach for CCID, we define

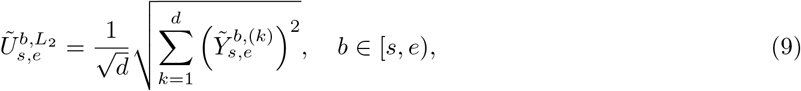

and we set 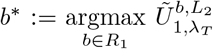. If this value exceeds a certain threshold, denoted by 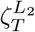, then it is taken as a change-point. If not, then the process tests the next interval in *S*_*RL*_, which in this case would be *L*_1_. Upon detection the ID methodology makes a new start from the end-point (or start-point, respectively) of the right- (or left-, respectively) expanding interval where the detection occurred. At some point we find an interval [*s, e*] that does not contain any other change-points because at each stop we expand the intervals by *λ*_*T*_ which is smaller than the minimum distance in (9) between two change-points; this is where isolation stems from as well, which in order to be ensured, the expansion parameter *λ*_*T*_ can be taken to be as small or equal to 1. If now *λ*_*T*_ > 1, then isolation is guaranteed with high probability. Theoretically for large *T*, the chosen value for *λ*_*T*_ (this typically is small; see Anastasiou & Fryzlewicz (2021) for more details) is guaranteed to be smaller than the minimum distance *δ_T_* (which is unknown but has to grow with *T*) between two consecutive change-points.

**Figure 1:**
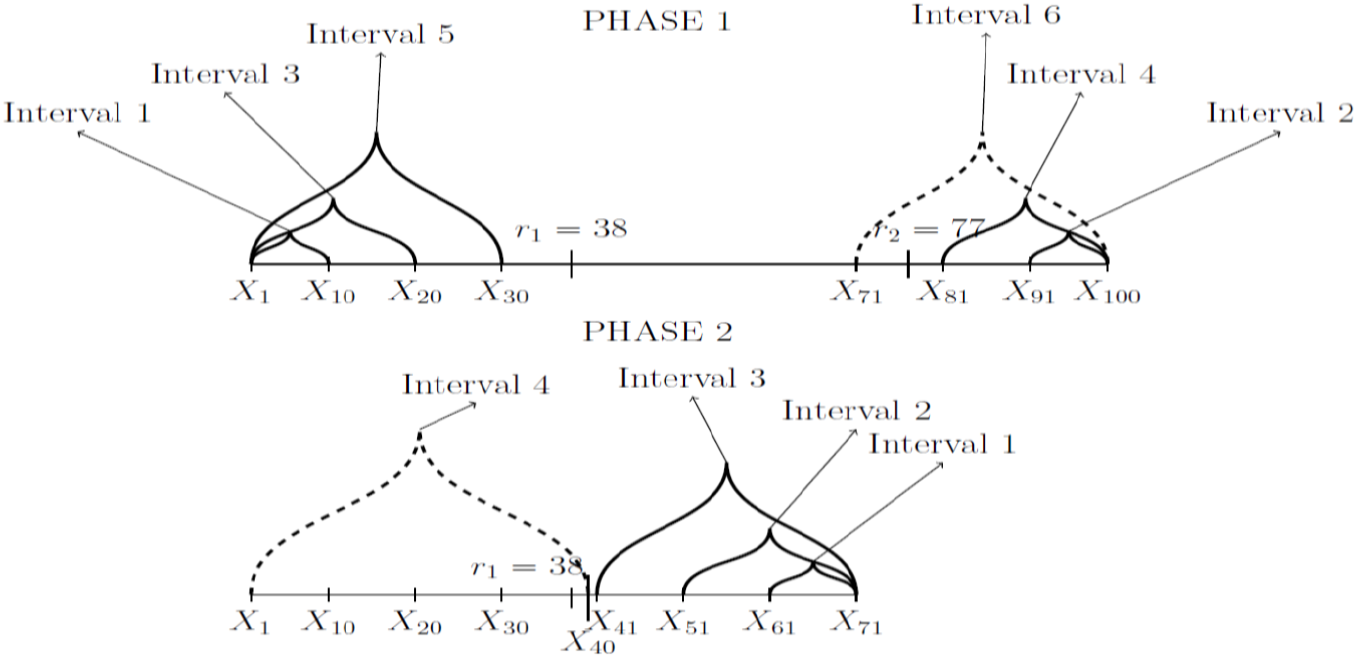
An example of how the segmentation method in CCID works. (Anastasiou & Fryzlewicz, 2021)

The *L*_∞_ approach works similarly. However, instead of using the Euclidean distance in (9), we use

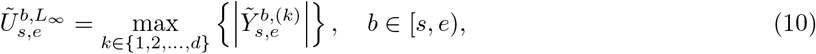

and we set 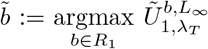. If this value exceeds a certain threshold, denoted by 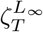, then it is taken as a change-point. We highlight that both 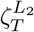 and 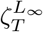are of the form 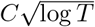, where *C* is a positive constant, with its choice discussed in Section 3.1. In the Gaussian case, 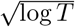 is the order of the threshold used in order to prove consistency (with respect to the estimated number and location of the change-points) results; more information can be found in Anastasiou & Fryzlewicz (2021). In this paper, our interest is in the actual numerical value of the threshold rather than in the theoretical order of its magnitude. Therefore, from now on, the 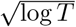 order is used, but it is calibrated through the constant, in order to optimise the performance of CCID in the specific applied framework described in this paper, which, is not of asymptotic interest.

A key ingredient of our CCID method is that, due to the aggregation of the (scaled) CUSUM statistics, the algorithm automatically identifies common change-points, rather than estimating single change-points at different locations in different components of the time series. This characteristic removes the need for post-processing across the *d*-dimensional sequence.

### 2.6. Matching the change-points to the relevant time series

We now explain how CCID can provide information on which brain region data sequence(s) each change-point arises from. The procedure to achieve this, is divided into two steps: firstly, estimate the change-points using CCID explained in Section 2.5, and secondly follow a post-processing approach, which is applied to each one of the *d* univariate data sequences in order to match the detected change-points (obtained from the first step) with the relevant data sequences. We highlight that the post-processing carried out in this section is completely different to the postprocessing carried out in Section 2.5 above. Specifically, we begin by estimating the number and the location of the change-points using either the *L*_2_ or the *L*_∞_ approach. The estimated change-points are then sorted in an increasing order in the set

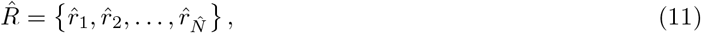

where 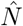 is the number of the estimated change-points. For 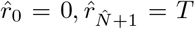, and 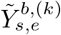 in (8), we then proceed, for each univariate data sequence 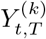as in (7), with the calculation of the quantities

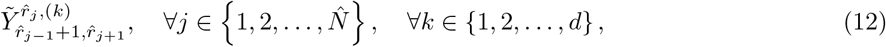

as defined in (8). Let us now denote by *S*^(*k*)^ the set of estimated change-points that appear in the *k*^*th*^ data sequence; as a note, keep in mind that, for any *k* ∈ {1, 2, …, *d*}, *S*^(*k*)^ is a subset of 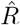 in (11). Since we now work in a univariate setting, checking all data sequences independently in order to match them with the obtained change-points, the values obtained in (12) are compared to a threshold value 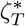. If, for example, 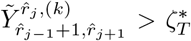, then it is deduced that the detected change-point 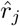 appears in the *k*^*th*^ data sequence. The threshold in this univariate setting is taken to be equal to 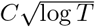. At the beginning of our approach the sets *S*^(*k*)^, *k* = 1, 2, …, *d* are empty. For each *k* ∈ {1, 2, …, *d*}, if 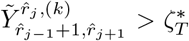, then the estimated change-point 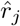 is added to the set *S*^(*k*)^. We complete this for all 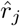 and for all the *d* data sequences, so that we operate on the same set of change-points, 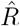 for each data sequence. Hence, the collection {*S*^(*k*)^}_*k*=1,2,…,*d*_ provides the information on which cross periodogram (and hence the pairwise relationships between brain regions at given scales of resolution) each change-point arises from. Our R package **ccid** also has a function that associates the detected change-points with the respective data sequence (or sequences) from which it was detected, but only for the threshold approach.

We now provide more details on the threshold constant *C* used in the post-processing procedure described in the previous paragraph. The threshold 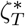 is of the same order as the threshold used for univariate change point detection in a series of papers, such as Anastasiou & Fryzlewicz (2021) and Fryzlewicz (2014). We note that 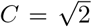 has been used in the aforementioned papers. More specifically, in Anastasiou & Fryzlewicz (2021) it was proved that for i.i.d. Gaussian noise and while *T* → ∞, the probability of a false detection when the null hypothesis of no change-point is satisfied goes to zero when 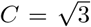. This means that, for our setting, it is sensible to take 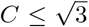. In finite sample size simulations, Fang et al. (2020) showed that for the method in Fryzlewicz (2014), under the Gaussian i.i.d. setting, when the sample size is equal to *T* = 500 and the threshold 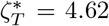 (which in our setting means *C* ≈ 1.853), then the results are very good only under the null model of no change-points, where the false positive rate is approximately 0.05. In the cases where there exist three, five, and eight change-points then the correct detection percentage with the aforementioned threshold is 82%, 64.3%, and 34.4%, respectively, which for the last two cases is especially low, meaning that a lower threshold is more suitable for data sets with multiple change-points. For CCID, we take 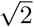, which is slightly higher than the proposed value of 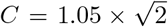 in Anastasiou & Fryzlewicz (2021) and Fryzlewicz (2014), while still lower than the aforementioned upper bound value of 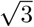. We note that this threshold is used as a post-processing tool when change-points have already been detected by CCID. Therefore, we are more interested in maintaining a good detection power when we have multiple change-points rather than having a low false positive rate in the case of no change-point; the latter is already controlled in the main algorithm.

### 2.7. The model selection approach

An alternative approach to the *L*_2_ and *L*_∞_ threshold methods is based on the optimization of a model selection criterion. In this setup, CCID begins by (hopefully) overestimating the number of change-points by choosing a suboptimal (lower) threshold value in the *L*_2_ or the *L*_∞_ methods explained in Section 2.5. The estimated change-points are sorted in an increasing order in the set 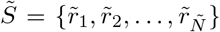, for 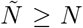 (the estimated number of change-points is greater than the true number of change-points). The next step is to run a change-point removal process using a joint approach for the *d* data sequences. For 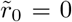 and 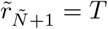, we collect the triplets 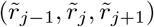 and calculate

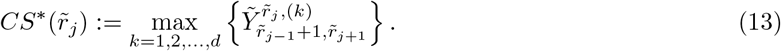

We then define 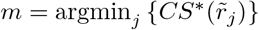, meaning that the estimation 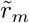 is the detected change-point which has the lowest (scaled) CUSUM statistic value in (13); 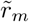 is labelled the least important detection in the set 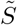. This change-point is removed from the set, reducing 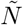 by 1. We relabel the remaining estimates (in increasing order) in 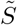, and repeat this estimate removal process until 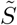 becomes the empty set. After this, the vector

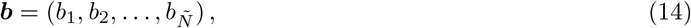

represents the collection of estimates where 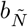 is the estimate that was first removed from 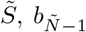 is the estimate that was removed next, and so on. The vector ***b*** is called the solution path and we define the collection of models 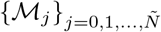 where 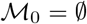 and 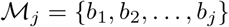.

A key element in the construction of the information criterion is the likelihood function for our data. Due to the unknown dependence structure in our data, we work instead on an approximation of the likelihood, where the data are taken to be independent (see Sections 4.1 and 5.1 for a discussion on deviations from independence); this, from now on, is called the pseudo-likelihood, denoted by 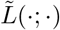 with its logarithm being 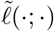. For fixed *k* ∈ {1, 2, …, *d*}, if there are no change-points in the data sequence 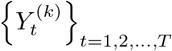 and 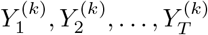 are independent random variables, then as 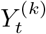 follows the scaled 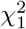 distribution with mean *σ*^(*k*)^, we have that

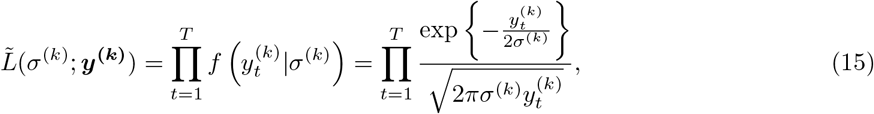

where 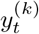 is the given *k*^*th*^ data sequence. Therefore,

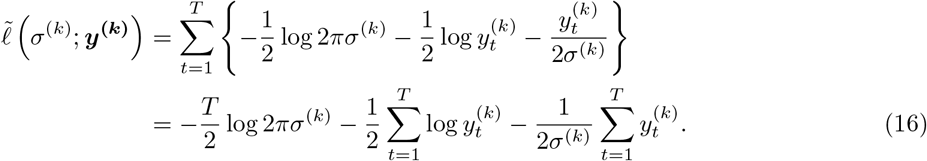

Using the maximum likelihood estimation method, based on (16), the estimator for the parameter *σ*^(*k*)^ is

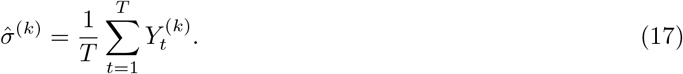

In the situation that we have *J* change-points at locations *r*_1_, *r*_2_, …, *r*_*J*_, then using (16) and (17), the pseudo-log-likelihood for the multivariate data sequence becomes

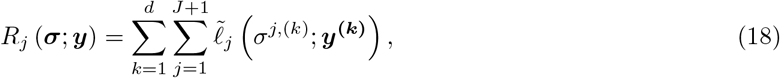

where *σ*^*j*,(*k*)^ is the mean of the *k*^*th*^ data sequence in the segment [*r*_*j*−1_ + 1, *r*_*j*_] and 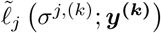 is the pseudo-log-likelihood in (16) for the aforementioned segment, where *r*_0_ = 0 and *r*_*J*+1_ = *T*. Therefore, from (16), we have that

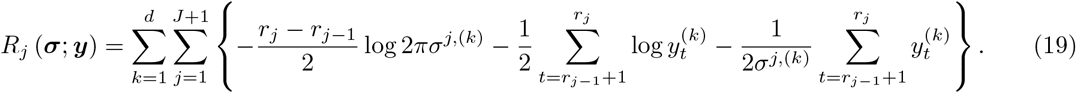

Among the collection of models 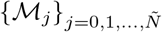, we propose to select the model that minimizes the following selection criterion

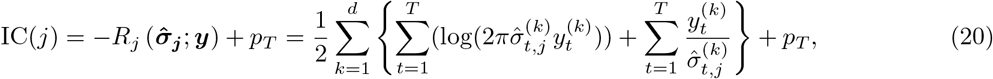

where *p*_*T*_ is an appropriately chosen penalty function which goes to infinity with *T*. For the definition of 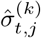, we denote for 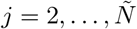,

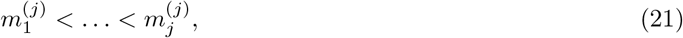

to be the sorted elements of 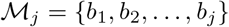. For example if *j* = 2 and *b*_2_ < *b*_1_, we have that 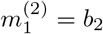 and 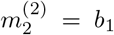; where *b*_1_ and *b*_2_ are the first two elements of the solution path as in (14). Then, for *i* = 0, 1, …, *j*,

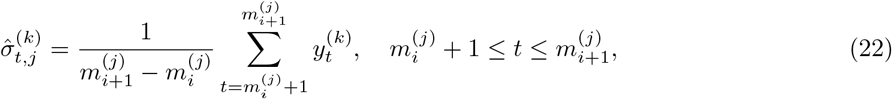

for 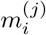, *i* ∈ {1, 2, …, *j*} as in (21) and 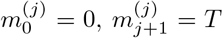, basically 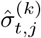, *t* = 1,2, …, *T* is the estimated piecewise-constant signal for the *k*^*th*^ data sequence when the detected change-points are *b*_1_, *b*_2_, …, *b*_*j*_. For the choice of *p*_*T*_, see Section 3.

### 2.8 Estimating brain networks

One of the main advantages of CCID over existing change-point methods is that CCID can detect change-points that are close to one another. In some cases, the number of change-points itself could be the objective of the study. However, in other cases, researchers often would like to estimate a partition-specific brain network or the undirected graph between each pair of detected change-points. This helps visualize the FC network in a more precise fashion. Estimating an undirected graph for a partition with a small number of data points and a large number of brain regions (*p* >> *T*) is difficult, especially in the case when *δ* (the minimum distance between change-points) is small (see Tables 9 and 10 in the Appendix). Nonetheless, CCID can be provided with a minimum distance between change-points as an input and it can postprocess the change-points to fit this criterion. If we let the full experimental time course be divided into subintervals separated by the detected change-points 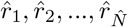 then for each subinterval 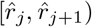, we can then estimate a sparse precision matrix to represent the FC brain network (Cribben & Fiecas, 2016) for each partition using the following log-likelihood with a SCAD penalty (Fan & Li, 2001) on the elements of the precision matrix (inverse covariance matrix)

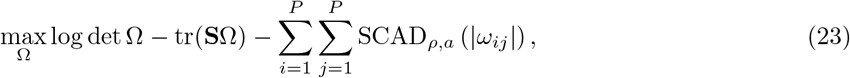

where Ω is the precision matrix, *S* is the sample covariance matrix, *ρ*_*ij*_(= *ρ* for convenience) is the penalty function on the elements of Ω, *ω*_*ij*_, and *w*_*ij*_ is the adaptive weight function. Fan & Li (2001) defined the adaptive weights to be

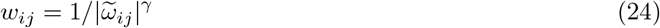

for tuning parameter *γ* > 0, where 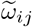 is the (*i, j*)th entry for any consistent estimate 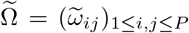. Mathematically, the SCAD penalty is symmetric and a quadratic spline on [0, ∞), whose first order derivative is

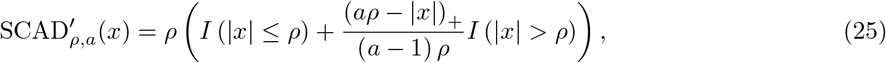

for *x* ≥ 0, where *I* is an indicator function, with *ρ* > 0 and *a* > 2 being two tuning parameters. Zhu & Cribben (2018) discuss the optimal choice for the tuning parameters in SCAD, and find that the best method for estimating sparse brain networks is SCAD in combination with the Bayesian Information Criterion.

## 3. Data

### 3.1 Simulation study setup

In this section we examine the performance of CCID through various simulations. For each simulation setting, we perform 100 repetitions, provide a diagram to illustrate how the second order structure (or FC network structure) changes over time and a quantified description of the results.

For CCID, we provide results for both the *L*_2_ and the *L*_∞_ approaches as described in Section 2.5 and the results for the information criterion discussed in Section 2.7. For the threshold, we chose the constant *c*_1_ = 0.65 for the *L*_2_ approach and the constant *c*_2_ = 2.25 for the *L*_∞_ approach as they provided a balance between specificity and sensitivity in all signals examined in a large scale simulation study; we highlight that the results presented in this paper are only a fraction of the various signals used in order to determine the value of the constants *c*_1_ and *c*_2_. It is important that CCID remains robust to alternative choices to this parameter and the practitioner has the option to obtain more change-points by decreasing this threshold value. We only present the results for the finest scale of −1 used in the wavelet transformation as fMRI data typically only have lag 1 autocorrelation (Fiecas et al., 2017). With regards to the information criterion model selection method, we define the penalty function by

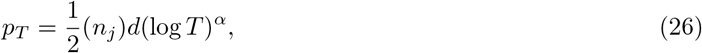

where *n_j_* is the total number of estimated parameters related to the model *M_j_*, *d* (= *p*(*p* + 1)/2) is the dimensionality of the wavelet-transformed data sequences (the periodograms/cross periodograms), *T* is the number of time points in each data sequence, and *α* > 0. We assume that *n_j_* manages to capture the complexity of the model under investigation. In (26), we propose a penalty function that is a function of the dimension of the periodograms and cross-periodograms, *d*. However, we should highlight that Zou et al. (2014) introduced a form of the Schwarz’s Information Criterion similar to (20) and they used a penalty of the form 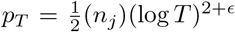, where *ϵ* > 0, and in practice *ϵ* = 0.1. Through the empirical work in this paper, where the length, *T*, of the data sequences is not larger than 1000 observations, we find that the penalty suggested by Zou et al. (2014) gives rise to too many false positive change-point detections. This is expected because the penalty used in Zou et al. (2014) does not depend on the dimensionality *d* of the time series, which can be quite large. A comparison of the two penalties is given through simulated data and it is available from Github at https://github.com/anastasiou-andreas/ccid. For both the simulated and fMRI data sets results that follow, we choose *α* = 0.1, which has been selected by considering many types of simulations involving a wide range of signals. The different structures used, consisted of general changes in the correlation/network structure, changes in the clustering (or community structure) and changes in the degree of a network, as well as no changes at all.

We compare CCID to three other competitor methods: Sparsified Binary Segmentation (SBS: Cho & Fryzlewicz, 2015) which is available through the **hdbinseg** R package, Factor (Barigozzi et al., 2018) which is available through the **factorcpt** R package, and the method developed in Barnett & Onnela (2016), which we denote by BO. There is no software available for BO, which is also the case for the majority of the methods mentioned in the introduction. However, for the BO approach, code was provided to us by the authors, which we had to adjust through a function that has already been written for the simulations carried out in Ofori-Boateng et al. (2020). For SBS, we only present the results for the finer scale of −1 used in the wavelet transformation. The Factor approach performs multiple change-point detection under factor modelling. The change-points are in the second order structure of both the common and idiosyncratic components. The main function in the R package **factorcpt** returns change-point estimates from the common and idiosyncratic components separately, and in our results we present both. The notation used for these is *Factor com* and *Factor id*, respectively.

### 3.2 Description of the settings

We now describe the simulations. The objective of the simulation study is to mimic the properties of fMRI data under various settings. While the data are simulated using various models, we display the dependence between the time series using an undirected graph (or network, which we use interchangeably) structure. A graph consists of a set of vertices *V* and corresponding edges *E* that connect pairs of vertices. Here each vertex represents a time series, or ROI, and edges encode dependencies. In the fMRI setting, a missing edge indicates a lack of functional connectivity between corresponding regions. A graph of **X** can alternatively be represented using the precision matrix (inverse covariance matrix) of **X**, with the elements of the matrix corresponding to edge weights. Here a missing edge between two vertices in the graph indicates conditional independence between the variables, giving rise to a zero element in the precision matrix.

- In Simulation 1, there are no true change-points. The data are simulated from a vector autoregression (VAR) model with *T* = 300 and *p* = 15. A VAR(1), a vector autoregression of order 1, is given by **X**_*t*_ = Π**X**_*t*−1_ + *ϵ*_*t*_, *t* = 2, …, *T*, where Π is an (*p* × *p*) coeffcient matrix and *ϵ*_*t*_ is an unobservable mean zero white noise vector process with time invariant covariance matrix. The VAR model is used to reconstruct the linear interdependency element prevalent among multivariate time series applications such as fMRI data (Cribben et al., 2012; Xu et al., 2020). For a visual display of the network structure, see Figure 2A.
- In Simulation 2, we consider a change in the network edge degree with *T* = 200 and *p* = 40. The change-point occurs in the middle of the time series with the edge degree changing from *η* = 0.95 to *η* = 0.05 before and after the change-point, respectively, where *η* represents the probability of a non-zero connection. The non-zero edges do not have the same strength. For a visual display of the change in the network structure, see Figure 2B. The simulation may appear quite easy (moving from a very dense graph to a very sparse graph), however, as the strength of the connections in both partitions are very small (≤ 0.1) the simulation in fact is more difficult than it appears.
- In Simulation 3, the data are simulated from a VAR model with *T* = 500 and *p* = 15. There are four true change-points and they are located at equal intervals across the whole time line. For a visual display of the changes in the network structure, see Figure 3A.
- In Simulation 4, the data are simulated from a VAR model with *T* = 600 and *p* = 10. There are seven true change-points and they are located at equal intervals across the whole time line. For a visual display of the changes in the network structure, see Figure 4A.
- In Simulation 5, the data are simulated from a VAR model with *T* = 500 and *p* = 15. There are four true change-points and they are located at equal intervals across the whole time line. This simulation has an ABABA structure. For a visual display of the changes in the network structure, see Figure 3B.
- In Simulation 6, there are seven true change-points and they are located at equal intervals across the whole time line. The data are simulated from a VAR model with *T* = 600 and *p* = 10. This simulation has an ABABABAB structure. For a visual display of the changes in the network structure, see Figure 4B.
- In Simulation 7, we again consider a change in the network edge degree, however, in this simulation, there are seven true change-points. The change-points are located at equal intervals across the whole time line with *T* = 600 and *p* = 40, and the edge degree changing from *η* = 0.95 to *η* = 0.05 before and after the change-point. This simulation has an ABABABAB structure. For a visual display of the change in the network structure, see Figure 5A.
- In Simulation 8, we consider a change in the network clustering (or community structure). There are again 7 change-points and they are located at equal intervals across the whole time line with *T* = 600 and *p* = 30. In the even time segments, the true number of communities *K*_*o*_ = 2, that is, there are two clusters, with the within cluster correlation equal to 0.8 and the between cluster correlation equal to 0. In the odd time segments, the true number of communities *K*_*o*_ = 6 with the within cluster correlation equal to 0.75 and the between cluster correlation equal to 0.2 (Cribben & Yu, 2017). This simulation has an ABABABAB structure. For a visual display of the change in the network structure, see Figure 5B.
- In Simulation 9, we consider a change in the network clustering (or community structure). It has exactly the same setup as Simulation 8 except the change-points are irregularly spaced with change points occurring at time points 100, 175, 275, 300, 400, 475, 575.
- In Simulation 10, we consider a change in the network clustering (or community structure). There are 3 change-points and they are located at time points 100, 175, 275, with *T* = 300 and *p* = 100. In the even time segments, the true number of communities *K*_*o*_ = 2, that is, there are two clusters, with the within cluster correlation equal to 0.8 and the between cluster correlation equal to 0. In the odd time segments, the true number of communities *K*_*o*_ = 20 with the within cluster correlation equal to 0.75 and the between cluster correlation equal to 0.2 (Cribben & Yu, 2017). This simulation has an ABAB structure and is a high dimension data example. For a visual display of the change in the network structure, see Figure 6.

**Figure 2:**
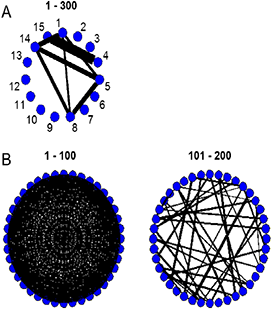
(A) The network structure for the VAR data set with no change-points (Simulation 1) and (B) The network structure for the change in the network degree with one change-point (Simulation 2). The ROI time series are represented by nodes, black edges infer positive connectivity, and the strength of connection between the ROIs is directly related to the thickness of the edges, that is, the thicker the edge the stronger the connection.

**Figure 3:**
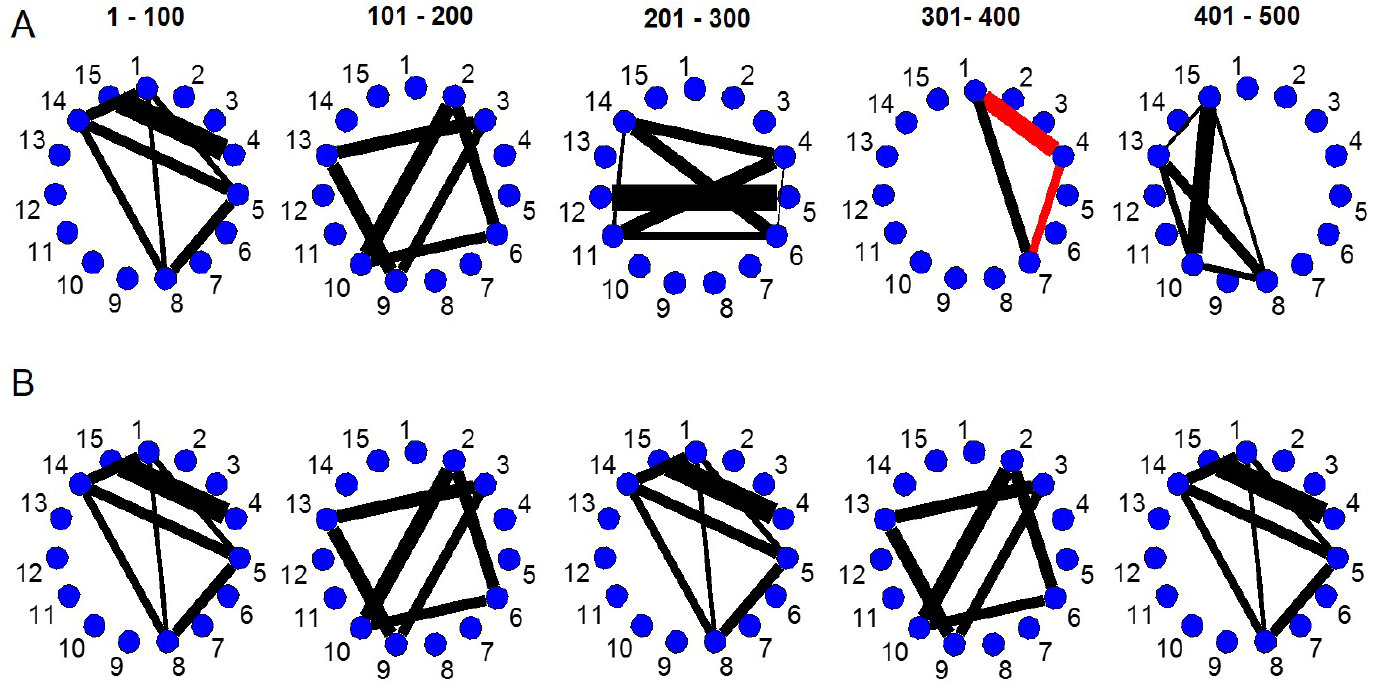
(A) The network structure for the VAR data set with four change-points (Simulation 3) and (B) The network structure for the VAR data set with four change-points (Simulation 5). The ROI time series are represented by nodes, black edges infer positive connectivity, and the strength of connection between the regions is directly related to the thickness of the edges, that is, the thicker the edge the stronger the connection.

**Figure 4:**
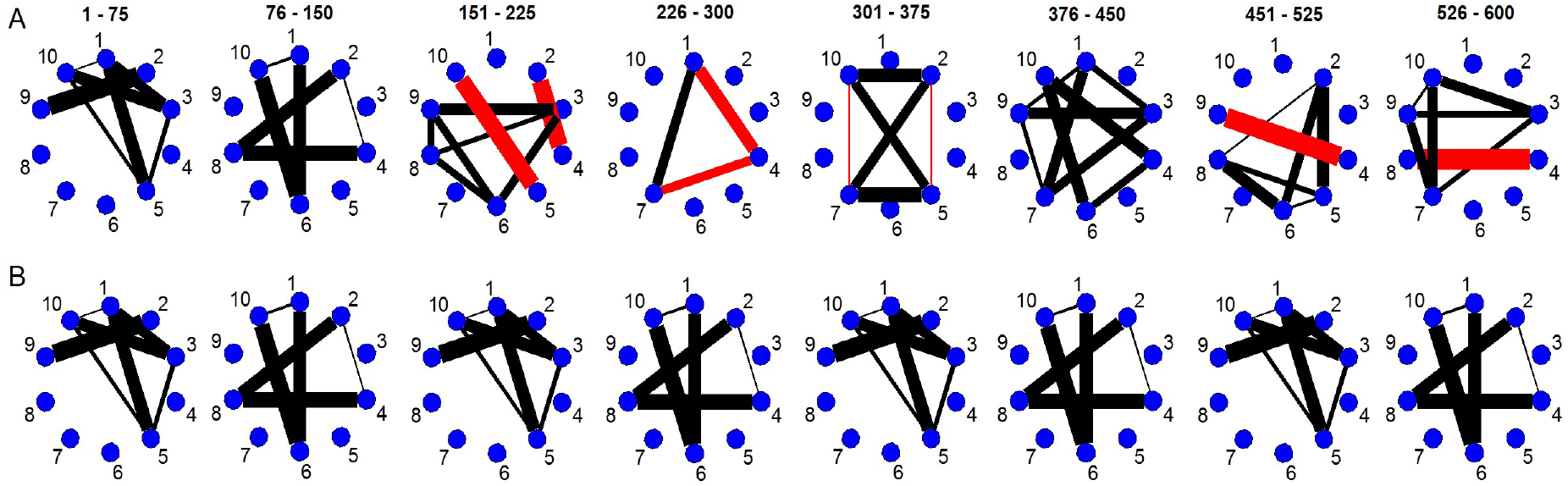
(A) The network structure for the VAR data set with seven change-points (Simulation 4) and (B) The network structure for the VAR data set with seven change-points (Simulation 6). The ROI time series are represented by nodes, black (red) edges infer positive (negative) connectivity, and the strength of connection between the regions is directly related to the thickness of the edges, that is, the thicker the edge the stronger the connection.

**Figure 5:**
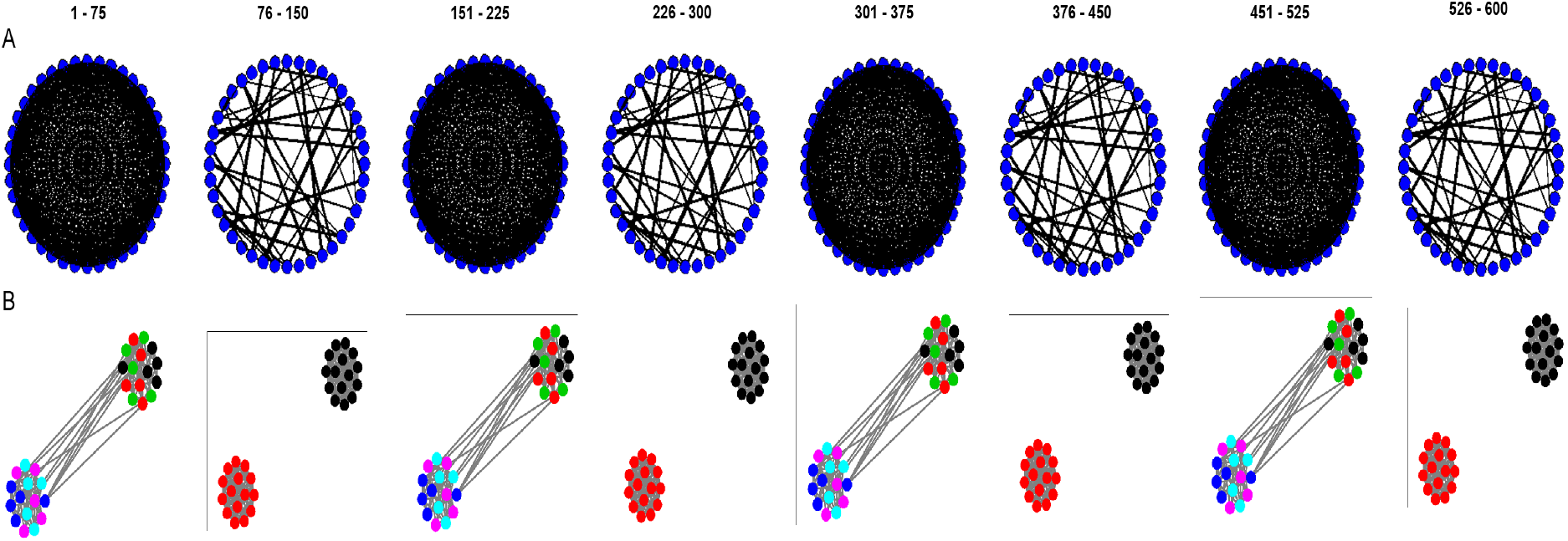
(A) The change in degree data set with seven change-points (Simulation 7). The ROI time series are represented by nodes, black edges infer positive connectivity, and the strength of connection between the regions is directly related to the thickness of the edges, that is, the thicker the edge the stronger the connection. (B) The change in clustering network structure with seven change-points (Simulation 8). In the even time segments, the true number of clusters is *K*_*o*_ = 2 (cluster membership is based on node color) within the within cluster correlation equal to 0.8 and the between cluster correlation equal to 0. In the odd time segments, the true number of clusters is *K*_*o*_ = 6 (cluster membership is based on node color) within the within cluster correlation equal to 0.75 and the between cluster correlation equal to 0.2.

**Figure 6:**
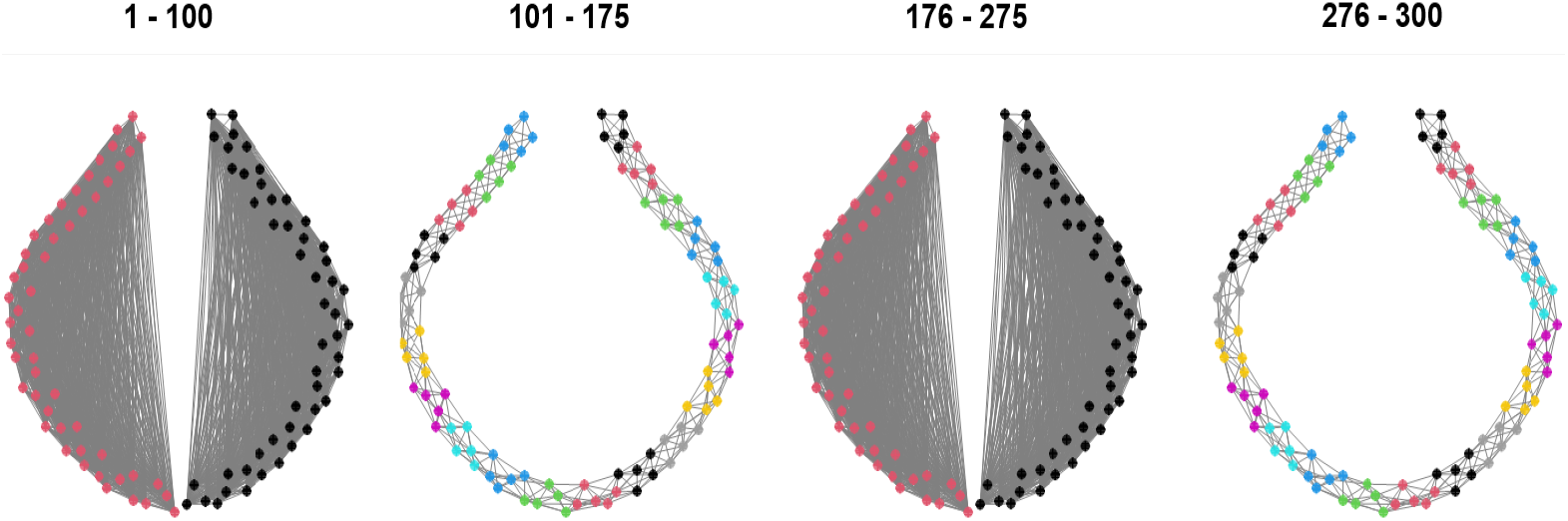
The change in clustering network structure with three change-points (Simulation 10). In the even time segments, the true number of clusters is *K*_*o*_ = 20 (cluster membership is based on node color) with the within cluster correlation equal to 0.8 and the between cluster correlation equal to 0. In the odd time segments, the true number of clusters is *K*_*o*_ = 2 (cluster membership is based on node color) with the within cluster correlation equal to 0.75 and the between cluster correlation equal to 0.2.

Simulation 1 is similar to a steady state fMRI time series, where the network structure does not change over time. Simulations 5 – 10 cover the situation where the subject alternates between 2 states (ABABA, ABABABAB and ABAB structures) and are similar to tasked based fMRI data sets. We hypothesize that our new method CCID will perform particularly well in these scenarios. This is due to CCID’s ability to isolate the change-points between tasks within subintervals and then to detect them with those subintervals. This is in contrast to the popular binary segmentation (BS) method which has been the only method applied on fMRI data. BS searches the entire time course for one change-point. Once a change-point is found, the data is split into two subsegments (hence, the term binary). A similar search is performed on each subsegment, possibly giving rise to further change-points. BS is a greedy algorithm as it is performed sequentially with each stage depending on the previous ones, which are never re-visited. Hence, BS looks to partition the experimental time course into two intervals which is difficult given the similarity between any two intervals for this particular data type. In other words, for any time points in an ABABA experimental structure, the before and after structures are very similar to the average of the A and B network structures, thus making change-points difficult to identify. However, CCID has the ability to find multiple change-points that are very close to one another.

### 3.3. Accuracy metrics

In the simulations, we compare the accuracy of the different methods using the frequency distribution of 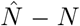, that is, the difference between the number of detected change-points and the number of true change-points. If 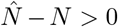, it indicates that the method finds too many false positive change-points, while if 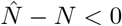, the method does not find all the true change-points. 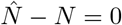 is ideal.

As a measure of the accuracy of the detected locations in time of the detected change-points compared to the location of the true change-points, we also provide the scaled Hausdorff distance,

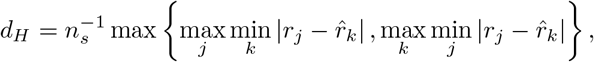

where *n*_*s*_ is the length of the largest segment, 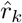 are the estimated change-points and *r*_*j*_ are the true change-points. The optimal model obtains a minimum scaled Hausdorff distance. The average computational time for all methods is also provided.

### 3.4. Task based fMRI data set

The data was taken from an anxiety-inducing experiment (Wager et al., 2009). The task was a variant of a well-studied laboratory paradigm for eliciting social threat, in which participants must give a speech under evaluative pressure. The design was an off-on-off design, with an anxiety-provoking speech preparation task occurring between lower anxiety resting periods. Participants were informed that they were to be given 2 min to prepare a 7 min speech, and that the topic would be revealed to them during scanning. They were told that after the scanning session they would deliver the speech to a panel of expert judges, though there was “small chance” they would be randomly selected not to give the speech. After the start of fMRI acquisition, participants viewed a fixation cross for 2 min (resting baseline). At the end of this period, participants viewed an instruction slide for 15 s that described the speech topic, which was “why you are a good friend”. The slide instructed participants to be sure to prepare enough for the entire 7 min period. After 2 min of silent preparation, another instruction screen appeared (a relief instruction, 15 s duration) that informed participants that they would not have to give the speech. An additional 2 min period of resting baseline completed the functional run.

Social threat is a cause of mental stress that generates physiological responses in the body (Rozanski et al., 1988). However, the connection between the human brain processes and the peripheral physiology while under stress of social threat is poorly understood. The objective of the study was to assess functional connectivity elicited by public speech preparation (SET) and its relationship with heart rate. Previous human neuroimaging studies (Critchley et al., 2000; Gianaros et al., 2004) indicated that the most likely areas for brain generators of cardiovascaular and other peripheral physiology responses to social evaluative threat are in the medial prefrontal cortex (MPFC), which projects to a set of cortical regions including the striatum. However, given MPFC’s heterogeneity, it may contain subregions with diverse relationships with regulation. In addition, animal studies implicated the ventromedial prefrontal cortex (Amat et al., 2005, 2008).

Functional blood-oxygen-level-dependent (BOLD) images were acquired with a T2*-sensitive spiral in-out sequence (TR (repetition time) = 2000 ms, TE (echo time) = 40 ms, flip angle = 90°, 24 slices in ascending sequential sequence, 4.5 × 3.4375 × 3.475 mm voxels). An LCD projector displayed stimuli on a back-projection screen placed in the scanner room. Functional images were subjected to a standard preprocessing sequence. Slice-timing acquisition correction using sync interpolation was performed using custom software, and realignment of the functional images to correct for head movement was performed using the Automated Image Registration tools (Woods et al., 1998). For details on the remaining preprocessing steps see Wager et al. (2009).

During the course of the experiment a series of 215 images were acquired (TR = 2 s). In order to create ROIs, the voxel time series were averaged across the entire regions. The data consists of 4 ROIs and heart rate for *n* = 23 subjects. The heart rate data was collected continuously during scanning using a photoplethysmograph on the left index finger (100 Hz sampling). Using customized software from the James Long Company, outliers were first identified and removed from the data (using a custom algorithm blind to task condition). Inter-beat intervals were then calculated from the remaining R-waves, and HR was averaged into 2 s bins (the scan repetition time, TR). For more details, see Wager et al. (2009). The temporal resolution of the heart rate was 1 s compared to 2 s for the fMRI data. Hence, the heart rate was down-sampled by taking every other measurement.

### 3.5. Resting state fMRI data set

The second data set is a resting-state fMRI data set, as described in Habeck et al. (2012); Cribben & Yu (2017). Participants (*n* = 45) are instructed to rest in the scanner for 9.5 minutes, with the instruction to keep their eyes open for the duration of the scan. Functional images were acquired using a 3.0 Tesla magnetic resonance scanner (Philips) using a field echo-planar imaging (FE-EPI) sequence [TE/TR = 20 ms/2000 ms; flip angle = 72°; 112×112 matrix; in-plane voxel size = 2.0 mm×2.0 mm; slice thickness = 3.0 mm (no gap); 37 transverse slices per volume]. In addition, a T1-weighted turbo field echo high resolution image was also acquired [TE/TR = 2.98 ms/6.57 ms; flip angle = 8°; 256×256 matrix; in-plane voxel size = 1.0 mm×1.0 mm; slice thickness = 1.0 mm (no gap); 165 slices]. The individual time series data were bandpass-filtered between 0.009 and 0.08 Hz, motion corrected and co-registered to the structural data, with a subsequent spatial normalization to the MNI template. The voxel time courses at white-matter and CSF locations are submitted to a Principal Component Analysis and, together with the motion parameters, we use all components with an eigenvalue strictly greater than 1 as independent variables in a subsequent nuisance regression Habeck et al. (2012). Each voxel’s time series is residualized with respect to those independent variables. The residual time series images are then smoothed with an isotropic Gaussian kernel (FWHM = 6mm). We apply the Anatomical Automatic Labeling (Tzourio-Mazoyer et al., 2002) atlas to the adjusted voxel-wise time series and produce time series for 31 Regions of Interest (ROIs) for each subject by averaging the voxel time series within the ROIs. The 31 ROIs contain 8 regions from the attentional network (frontal superior medial L, angular L, angular R, temporal middle L, temporal mid R, thalamus L, cerebellum crus1 L, cerebellum crus1 R), 2 regions from the visual network (temporal superior L, temporal superior R), 3 regions from the sensorimotor network (postcentral L, postcentral R, supplementary motor area R), 7 regions the salience network (cingulum anterior L, frontal mid L, frontal middle R, insula L, insula R, supramarginal L, supramarginal R), 9 regions from the default mode network (precentral L, precentral R, parietal superior L, occipital superior R, parietal inferior L, parietal inferior R, temporal inferior L, temporal inferior R, cingulum posterior L) and 2 regions from the auditory network (calcarine L, Calcarine R). We chose these networks because an increasing number of pathologic conditions appear to be reflected in the functional connectivity between these particular brain regions and we wanted the number of ROIs in the fMRI data to match the simulation settings. In total, each ROI time series is made up of 285 time points (9.5 minutes with TR = 2).

## 4. Results

### 4.1. Simulation study

In this section, we present the simulation and fMRI results. Table 1 shows the results over 100 iterations for Simulation 1 (no change-point: see Section 3.2). SBS (Section 3.1), Factor (Section 3.1), and our method, CCID (across both thresholds and information criteria), perform very well and correctly do not detect change-points across almost all iterations. BO has the worst performance in terms of Type I error. CCID is at least an order of magnitude computationally faster than all competing methods, a pattern which is also evident for the rest of the simulations.

**Table 1:**
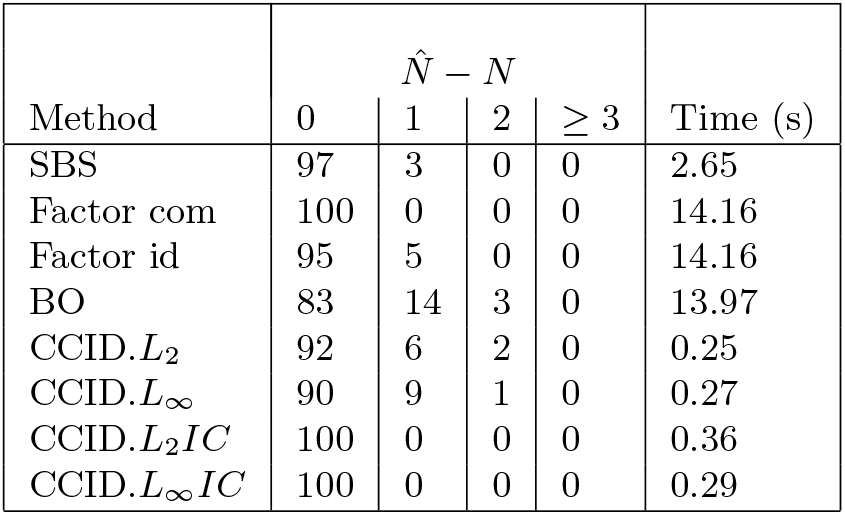
The distribution of 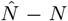 over 100 simulated data sequences from Simulation 1 (no change-point) for Sparsified Binary Segmentation (SBS), Factor with common components (Factor com), Factor with idiosyncratic components (Factor id), Barnett & Onnela (2016) (BO), and Cross-covariance isolate detect with the *L*_2_ threshold (CCID.*L*_2_), the *L*_2_ threshold (CCID.*L*_∞_), the *L*_2_ information criterion (CCID.*L*_2_*IC*) and the *L*_∞_ information criterion (CCID.*L*_∞_*IC*). The computational times for each method are also provided.

Table 2 shows the results over 100 iterations for Simulation 2 (change in network degree, see Section 3.2). As mentioned previously, this simulation is more difficult than it appears. SBS, Factor with idiosyncratic components (Factor id), BO and CCID (except for CCID.*L*_∞_) perform very well and correctly identify the change-point in the vast majority of the 100 iterations, with Factor id performing the best, finding one change point in all 100 iterations. On the other hand, Factor with common components (Factor com) performs poorly and does not detect the change-point in any of the 100 iterations. In terms of Hausdorff distance, Factor id performs the best but there is a negligible difference with SBS, BO, CCID.*L*_2_, CCID.*L*_2_*IC* and CCID.*L*_∞_*IC*.

**Table 2:** The distribution of 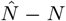 and the scaled Hausdorff distance over 100 simulated data sequences from Simulation 2 (change in network degree) for Sparsified Binary Segmentation (SBS), Factor with common components (Factor com), Factor with idiosyncratic components (Factor id), Barnett & Onnela (2016) (BO), and Cross-covariance isolate detect with the *L*_2_ threshold (CCID.*L*_2_), the *L*_∞_ threshold (CCID.*L*_∞_), the *L*_2_ information criterion (CCID.*L*_2_*IC*) and the *L*_∞_ information criterion (CCID.*L*_∞_*IC*). The computational times for each method are also provided.

| Method | $\hat{N} - N$ | | | | $d_H$ | Time (s) |
| --- | --- | --- | --- | --- | --- | --- |
| | -1 | 0 | 1 | $\geq 2$ | | |
| SBS | 0 | 99 | 1 | 0 | 0.02 | 11.20 |
| Factor com | 100 | 0 | 0 | 0 | 1 | 15.39 |
| Factor id | 0 | 100 | 0 | 0 | 0.01 | 15.39 |
| BO | 0 | 93 | 7 | 0 | 0.05 | 7.78 |
| CCID.$L_2$ | 5 | 93 | 0 | 2 | 0.06 | 1.00 |
| CCID.$L_\infty$ | 0 | 57 | 24 | 14 | 0.26 | 1.10 |
| CCID.$L_2IC$ | 3 | 93 | 4 | 0 | 0.07 | 1.62 |
| CCID.$L_\infty IC$ | 3 | 94 | 3 | 0 | 0.06 | 2.41 |

For Simulation 3 (see Section 3.2), SBS, BO and CCID (using the thresholding and information criteria) perform well with the majority of their detected change-points within ±2 (Table 3). However, CCID.*L*_2_*IC* performs the best in terms of the distribution of 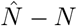, that is, the majority of its detected change-points equal the true number of change-points across all 100 iterations, but BO has a slightly lower Hausdorff distance, the detected change-points are closest to the true change-points in terms of location. Both Factor com and Factor id perform poorly in this simulation.

**Table 3:** The distribution of 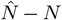 and the scaled Hausdorff distance over 100 simulated data sequences from Simulation 3 for Sparsified Binary Segmentation (SBS), Factor with common components (Factor com), Factor with idiosyncratic components (Factor id), Barnett & Onnela (2016) (BO), and Cross-covariance isolate detect with the *L*_2_ threshold (CCID.*L*_2_), the *L*_∞_ threshold (CCID.*L*_∞_), the *L*_2_ information criterion (CCID.*L*_2_*IC*) and the *L*_∞_ information criterion (CCID.*L*_∞_*IC*). The computational times for each method are also provided.

| Method | $\hat{N} - N$ | | | | | | | $d_H$ | Time (s) |
| --- | --- | --- | --- | --- | --- | --- | --- | --- | --- |
| | $\leq -3$ | -2 | -1 | 0 | 1 | 2 | $\geq 3$ | | |
| SBS | 4 | 6 | 18 | 72 | 0 | 0 | 0 | 0.51 | 5.93 |
| Factor com | 100 | 0 | 0 | 0 | 0 | 0 | 0 | 4 | 15.38 |
| Factor id | 74 | 7 | 9 | 10 | 0 | 0 | 0 | 2.73 | 15.38 |
| BO | 0 | 0 | 0 | 72 | 22 | 6 | 0 | 0.22 | 45.45 |
| $\text{CCID}.L_2$ | 0 | 2 | 37 | 57 | 4 | 0 | 0 | 0.46 | 0.42 |
| $\text{CCID}.L_\infty$ | 0 | 3 | 13 | 71 | 10 | 3 | 1 | 0.36 | 0.46 |
| $\text{CCID}.L_2IC$ | 1 | 2 | 10 | 80 | 6 | 1 | 0 | 0.27 | 0.82 |
| $\text{CCID}.L_\infty IC$ | 9 | 7 | 7 | 72 | 5 | 0 | 0 | 0.66 | 0.63 |

Simulation 4 (see Section 3.2) contains 7 change-points with CCID (across both thresholds and information criteria) outperforming SBS and Factor in terms of detection and having the smallest Hausdorff distances (Table 4). However, BO also performs well and has the third best Hausdorff distance after CCID.*L*_2_ and CCID.*L*_2_*IC*. CCID.*L*_2_*IC* performs the best while SBS is able to detect some change points, however, Factor performs poorly in this simulation with large negative values for 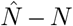 and large Hausdorff distance values. BO is particularly slow in this simulation.

**Table 4:** The distribution of 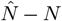 and the scaled Hausdorff distance over 100 simulated data sequences from Simulation 4 for Sparsified Binary Segmentation (SBS), Factor with common components (Factor com), Factor with idiosyncratic components (Factor id), Barnett & Onnela (2016) (BO), and Cross-covariance isolate detect with the *L*_2_ threshold (CCID.*L*_2_), the *L*_∞_ threshold (CCID.*L*_∞_), the *L*_2_ information criterion (CCID.*L*_2_*IC*) and the *L*_∞_ information criterion (CCID.*L*_∞_*IC*). The computational times for each method are also provided.

| Method | $\hat{N} - N$ | | | | | | | $d_H$ | Time (s) |
| --- | --- | --- | --- | --- | --- | --- | --- | --- | --- |
| | $\leq -3$ | -2 | -1 | 0 | 1 | 2 | $\geq 3$ | | |
| SBS | 49 | 28 | 21 | 2 | 0 | 0 | 0 | 1.66 | 4.27 |
| Factor com | 100 | 0 | 0 | 0 | 0 | 0 | 0 | 7 | 16.12 |
| Factor id | 100 | 0 | 0 | 0 | 0 | 0 | 0 | 5.03 | 16.12 |
| BO | 0 | 0 | 1 | 43 | 34 | 16 | 6 | 0.30 | 95.80 |
| $\text{CCID}.L_2$ | 0 | 0 | 1 | 52 | 39 | 7 | 1 | 0.24 | 0.21 |
| $\text{CCID}.L_\infty$ | 12 | 10 | 27 | 36 | 13 | 1 | 1 | 0.94 | 0.27 |
| $\text{CCID}.L_2IC$ | 0 | 1 | 9 | 71 | 14 | 3 | 2 | 0.26 | 0.92 |
| $\text{CCID}.L_\infty IC$ | 4 | 14 | 23 | 54 | 5 | 0 | 0 | 0.65 | 0.47 |

Simulations 1–4 show that CCID performs as well (and in most cases better than) the competing methods for the general settings. We now consider Simulations 5–8 which cover the situation where the subject changes between 2 states (ABABA and ABABABA structures) and mimic a tasked based fMRI experiment. This is a generalization of the epidemic change alternative (Kirch et al., 2015), change where change-point *r*_1_ corresponds to the time when a process **Π**_1_ is turned off (equivalently, when another process **Π**_2_ is activated) and change-point *r*_2_ corresponds to the time **Π**_2_ is turned off, thus returning to **Π**_1_. As we have stated before, we hypothesize that our new method, CCID, will perform particularly well in these scenarios. This is due to CCID’s ability to isolate the change-points between tasks within subintervals and then to detect them with those subintervals. In addition, CCID has the ability to find change-points that are very close to one another.

Table 5 shows the results from Simulation 5 (see Section 3.2), which has four change-points and an ABABA network structure. Clearly CCID with both thresholds and information criteria outperform SBS and Factor com in terms of estimating the correct number of change points and the location of the detected change points. In particular, CCID.*L*_2_ has the best performance, correctly identifying the correct number of change points in 91 out of 100 iterations and having the smallest Hausdorff distance. Furthermore, CCID.*L*_∞_ also performs well with similar but superior results to Factor id and BO.

**Table 5:**
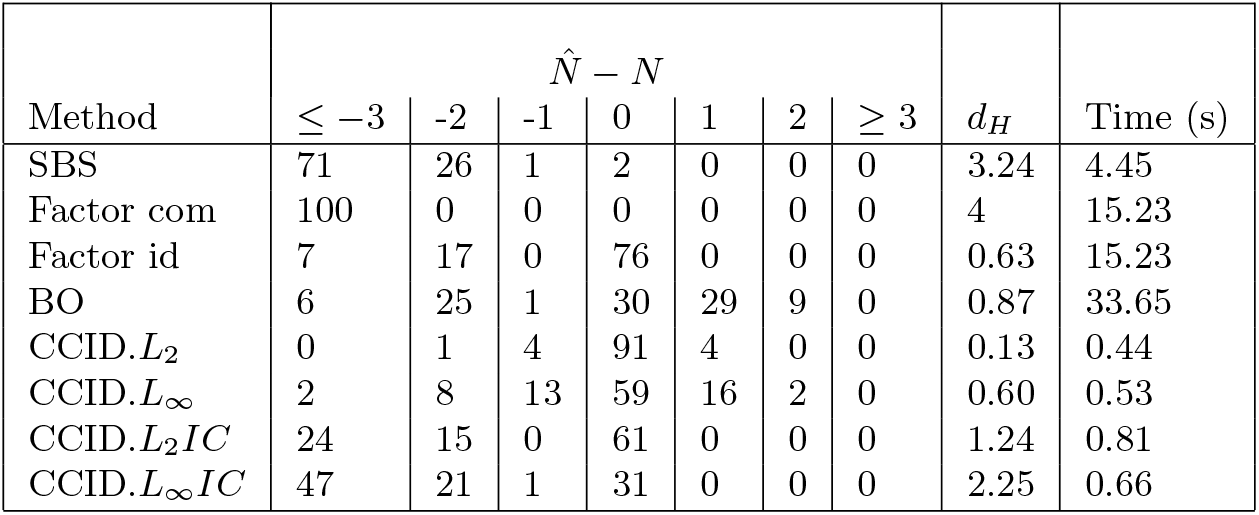
The distribution of 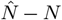 and the scaled Hausdorff distance over 100 simulated data sequences from Simulation 5 for Sparsified Binary Segmentation (SBS), Factor with common components (Factor com), Factor with idiosyncratic components (Factor id), Barnett & Onnela (2016) (BO), and Cross-covariance isolate detect with the *L*_2_ threshold (CCID.*L*_2_), the *L*_∞_ threshold (CCID.*L*_∞_), the *L*_2_ information criterion (CCID.*L*_2_*IC*) and the *L*_∞_ information criterion (CCID.*L*_∞_*IC*). The computational times for each method are also provided.

Table 6 shows the results from Simulation 6 (see Section 3.2), which has seven network change points and an ABABABAB network structure. Again, CCID with both thresholds and information criteria outperforms all of SBS, Factor and BO. In particular, CCID.*L*_2_ has the best performance in terms of estimating the correct number of change points and the location of the detected change points. SBS and Factor perform poorly in this simulation, they are unable to identify the correct number of change points in all iterations. BO is able to estimate the correct number of change points in 10 out of the 100 iterations but is very slow computationally compared to all the other methods.

**Table 6:** The distribution of 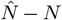 and the scaled Hausdorff distance over 100 simulated data sequences from Simulation 6 for Sparsified Binary Segmentation (SBS), Factor with common components (Factor com), Factor with idiosyncratic components (Factor id), Barnett & Onnela (2016) (BO), and Cross-covariance isolate detect with the *L*_2_ threshold (CCID.*L*_2_), the *L*_∞_ threshold (CCID.*L*_∞_), the *L*_2_ information criterion (CCID.*L*_2_*IC*) and the *L*_∞_ information criterion (CCID.*L*_∞_*IC*). The computational times for each method are also provided.

| Method | $\hat{N} - N$ | | | | | | | $d_H$ | Time (s) |
| --- | --- | --- | --- | --- | --- | --- | --- | --- | --- |
| | $\leq -3$ | -2 | -1 | 0 | 1 | 2 | $\geq 3$ | | |
| SBS | 100 | 0 | 0 | 0 | 0 | 0 | 0 | 6.73 | 2.31 |
| Factor com | 100 | 0 | 0 | 0 | 0 | 0 | 0 | 7 | 17.14 |
| Factor id | 100 | 0 | 0 | 0 | 0 | 0 | 0 | 6.72 | 17.14 |
| BO | 47 | 13 | 4 | 10 | 13 | 8 | 5 | 2.38 | 53.33 |
| CCID.$L_2$ | 0 | 0 | 1 | 68 | 23 | 7 | 1 | 0.20 | 0.11 |
| CCID.$L_\infty$ | 40 | 26 | 13 | 17 | 4 | 0 | 0 | 1.86 | 0.17 |
| CCID.$L_2IC$ | 9 | 12 | 2 | 65 | 12 | 0 | 0 | 0.75 | 0.46 |
| CCID.$L_\infty IC$ | 34 | 26 | 5 | 33 | 2 | 0 | 0 | 2.28 | 0.25 |

Table 7 shows the results from Simulation 7 (see Section 3.2), which has seven change points in the network degree ABABABAB type structure. Similar to Simulation 2, this simulation (see Figure 5A) may appear quite easy (alternating between a very dense graph to a very sparse graph), however, as the strength of the connections in both partitions is very small (≤ 0.1) the simulation in fact is more difficult than it appears. Both SBS and Factor com struggle, they are unable to identify the correct number of change points in all iterations. CCID with both thresholds and information criteria outperforms SBS and Factor. BO, while superior to both SBS and Factor, is still inferior to CCID across both thresholds and information criteria. CCID.*L*_∞_*IC* has the best performance in terms of estimating the correct number of change points and the location of the detected change points. In all 100 iterations, it identifies the correct number of true change points. However, Factor id performs adequately but in all iterations misses two change points.

**Table 7:**
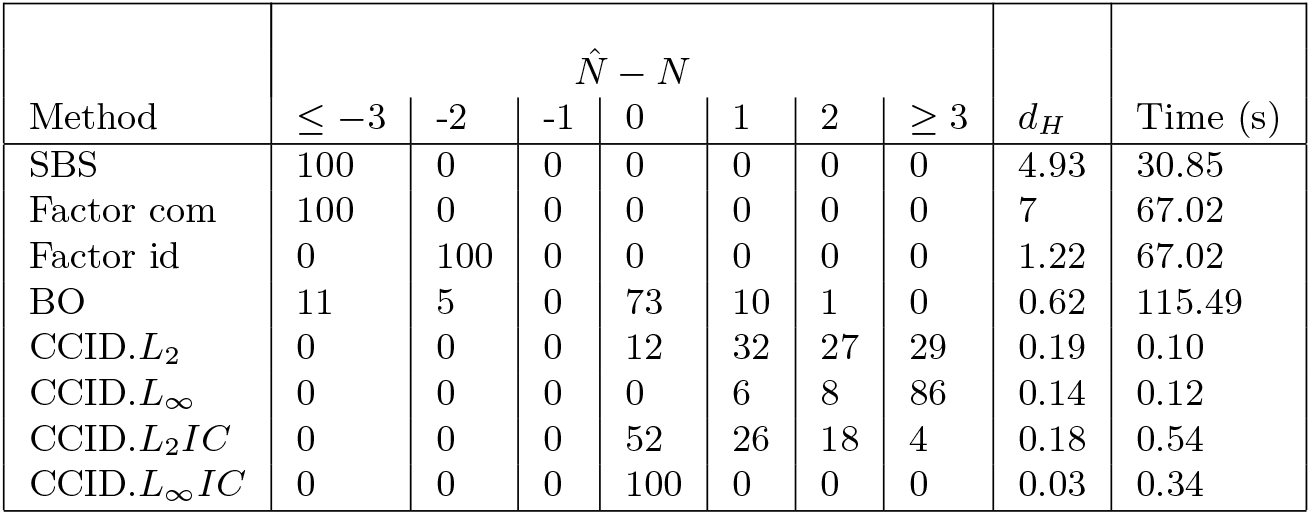
The distribution of 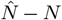 and the scaled Hausdorff distance over 100 simulated data sequences from Simulation 7 for Sparsified Binary Segmentation (SBS), Factor with common components (Factor com), Factor with idiosyncratic components (Factor id), Barnett & Onnela (2016) (BO), and Cross-covariance isolate detect with the *L*_2_ threshold (CCID.*L*_2_), the *L*_∞_ threshold (CCID.*L*_∞_), the *L*_2_ information criterion (CCID.*L*_2_*IC*) and the *L*_∞_ information criterion (CCID.*L*_∞_*IC*). The computational times for each method are also provided.

Table 8 shows the results from Simulation 8 (see Section 3.2), which has seven change points in the network clustering ABABABAB type structure. This is the most difficult simulation but, again, CCID with both thresholds and information criteria outperform SBS, Factor and BO. Similar to Simulation 7, CCID.*L*_∞_*IC* has the best performance in terms of estimating the correct number of change points and the location of the detected change points. SBS and Factor perform poorly in this simulation, they are unable to identify the correct number of change points in all iterations. BO has a decent performance but is outperformed by CCID across all thresholds and information criteria.

**Table 8:** The distribution of 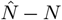 and the scaled Hausdorff distance over 100 simulated data sequences from Simulation 8 for Sparsified Binary Segmentation (SBS), Factor with common components (Factor com), Factor with idiosyncratic components (Factor id), Barnett & Onnela (2016) (BO), and Cross-covariance isolate detect with the *L*_2_ threshold (CCID.*L*_2_), the *L*_∞_ threshold (CCID.*L*_∞_), the *L*_2_ information criterion (CCID.*L*_2_*IC*) and the *L*_∞_ information criterion (CCID.*L*_∞_*IC*). The computational times for each method are also provided.

| Method | $\hat{N} - N$ | | | | | | | $d_H$ | Time (s) |
| --- | --- | --- | --- | --- | --- | --- | --- | --- | --- |
| | $\leq -3$ | -2 | -1 | 0 | 1 | 2 | $\geq 3$ | | |
| SBS | 100 | 0 | 0 | 0 | 0 | 0 | 0 | 5.82 | 16.85 |
| Factor com | 100 | 0 | 0 | 0 | 0 | 0 | 0 | 7 | 423.40 |
| Factor id | 100 | 0 | 0 | 0 | 0 | 0 | 0 | 6.63 | 423.40 |
| BO | 39 | 5 | 1 | 29 | 22 | 4 | 0 | 2.11 | 82.27 |
| $\text{CCID}.L_2$ | 0 | 0 | 0 | 42 | 28 | 20 | 10 | 0.27 | 0.82 |
| $\text{CCID}.L_\infty$ | 0 | 0 | 0 | 2 | 6 | 13 | 79 | 0.30 | 1.26 |
| $\text{CCID}.L_2 IC$ | 0 | 0 | 0 | 79 | 13 | 5 | 3 | 0.19 | 3.79 |
| $\text{CCID}.L_\infty IC$ | 0 | 0 | 1 | 94 | 4 | 1 | 0 | 0.11 | 3.12 |

Table 9 shows the results from Simulation 9 (see Section 3.2), which has seven change points (ABABABAB) in the network clustering structure. It has the same setup as Simulation 8, but the true change-points are occurring at unequally spaced time points, namely at time points *t* = 100, 175, 275, 300, 400, 475, 575. CCID with both thresholds and information criteria outperform SBS, Factor and BO, and to a larger extent than the results for Simulation 8. Again, similar to Simulation 8, the information criteria have the best performance in terms of estimating the correct number of change points and the location of the detected change points (smallest Hausdorff distance), with CCID.L_∞_IC detecting the true number of change points on 89 out of 100 iterations. SBS and Factor perform poorly in this simulation, they are unable to identify the correct number of change points across all iterations. BO has a moderate performance but is outperformed by CCID across all thresholds and information criteria.

**Table 9:** The distribution of 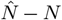 and the scaled Hausdorff distance over 100 simulated data sequences from Simulation 9 for Sparsified Binary Segmentation (SBS), Factor with common components (Factor com), Factor with idiosyncratic components (Factor id), Barnett & Onnela (2016) (BO), and Cross-covariance isolate detect with the *L*_2_ threshold (CCID.*L*_2_), the *L*_∞_ threshold (CCID.*L*_∞_), the *L*_2_ information criterion (CCID.*L*_2_*IC*) and the *L*_∞_ information criterion (CCID.*L*_∞_*IC*). The computational times for each method are also provided.

| Method | $\hat{N} - N$ | | | | | | | $d_H$ | Time (s) |
| --- | --- | --- | --- | --- | --- | --- | --- | --- | --- |
| | $\leq -3$ | -2 | -1 | 0 | 1 | 2 | $\geq 3$ | | |
| SBS | 100 | 0 | 0 | 0 | 0 | 0 | 0 | 5.21 | 14.11 |
| Factor com | 100 | 0 | 0 | 0 | 0 | 0 | 0 | 5.64 | 49.32 |
| Factor id | 100 | 0 | 0 | 0 | 0 | 0 | 0 | 5.75 | 49.32 |
| BO | 72 | 18 | 8 | 2 | 0 | 0 | 0 | 3.63 | 68.49 |
| CCID.$L_2$ | 0 | 0 | 0 | 16 | 24 | 25 | 45 | 0.36 | 2.01 |
| CCID.$L_\infty$ | 0 | 0 | 0 | 2 | 3 | 11 | 84 | 0.22 | 2.84 |
| CCID.$L_2IC$ | 1 | 12 | 0 | 63 | 13 | 9 | 2 | 0.29 | 5.48 |
| CCID.$L_\infty IC$ | 0 | 3 | 0 | 89 | 4 | 4 | 0 | 0.10 | 3.98 |

Table 10 shows the results from Simulation 10 (see Section 3.2), which has three change points in the network clustering structure (ABAB). Here, we consider a high-dimensional time series (*p* = 100). CCID with both information criteria has the best performance, BO and CCID with both thresholds perform similarly and have the next best performance, with Factor having the worst performance (it is unable to identify the correct number of change points in all iterations). BO has a moderate performance but is outperformed by CCID with the information criteria. CCID.L_∞_IC detects the true number of change points on 89 out of 100 iterations. The computation for CCID is an order of magnitude faster than SBS and two orders of magnitudes faster than BO and Factor. Finally, it is important to point out that the true dimensionality of the problem is *d*(*d* + 1)/2, where *d* is the dimensionality of the initial multivariate time series. Hence, for given data of dimensionality *p* = 100, the dimensionality of the problem is *d* = 5050 (periodograms).

**Table 10:** The distribution of 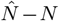 and the scaled Hausdorff distance over 100 simulated data sequences from Simulation 10 for Sparsified Binary Segmentation (SBS), Factor with common components (Factor com), Factor with idiosyncratic components (Factor id), Barnett & Onnela (2016) (BO), and Cross-covariance isolate detect with the *L*_2_ threshold (CCID.*L*_2_), the *L*_∞_ threshold (CCID.*L*_∞_), the *L*_2_ information criterion (CCID.*L*_2_*IC*) and the *L*_∞_ information criterion (CCID.*L*_∞_*IC*). The computational times for each method are also provided.

| Method | $\hat{N} - N$ | | | | | | | $d_H$ | Time (s) |
| --- | --- | --- | --- | --- | --- | --- | --- | --- | --- |
| | -3 | -2 | -1 | 0 | 1 | 2 | $\geq 3$ | | |
| SBS | 0 | 0 | 95 | 5 | 0 | 0 | 0 | 1.01 | 81.91 |
| Factor com | 100 | 0 | 0 | 0 | 0 | 0 | 0 | 2.75 | 568.72 |
| Factor id | 5 | 0 | 95 | 0 | 0 | 0 | 0 | 1.08 | 568.72 |
| BO | 19 | 0 | 3 | 50 | 25 | 2 | 1 | 0.67 | 712.49 |
| CCID.$L_2$ | 0 | 0 | 0 | 36 | 21 | 20 | 23 | 0.28 | 0.97 |
| CCID.$L_\infty$ | 0 | 0 | 0 | 0 | 1 | 0 | 99 | 0.57 | 1.38 |
| CCID.$L_2IC$ | 0 | 0 | 0 | 73 | 9 | 8 | 10 | 0.15 | 3.86 |
| CCID.$L_\infty IC$ | 0 | 0 | 0 | 89 | 4 | 4 | 3 | 0.08 | 3.25 |

As stated at the beginning of Section 2, CCID assumes that the ROI time series are Gaussian. It also assumes independence in the pseudo-likelihood as expressed in (15). However, CCID is robust under deviations from Gaussianity and/or from independence within the ROI time series. For example, Table 13 (in the Appendix) shows the results from applying CCID (and the competing methods, SBS, Factor and BO) to Simulations 1, 3 and 5, where the noise of each ROI time series now follows the *t* distribution with 5 degrees of freedom. While the results are inferior to the Gaussian data, there is not a vast drop off in performance. The performance of CCID remains superior to SBS, Factor and BO in these settings. In situations where we suspect that our data are far from independent and Gaussian, our CCID algorithm can still be applied to the data after some pre-processing takes place. More specifically, towards this purpose, the subsampling and pre-averaging techniques described in Section 5.1 are beneficial.

In summary, for general data sets (Simulations 1–4), we found that CCID performs as well as, if not better than, SBS, Factor and BO. For simulations where the subject alternates between two states (Simulations 5–10), CCID performs very well and clearly outperforms SBS, Factor and BO using both thresholds and information criteria. Our recommendation is to use CCID with thresholding for general settings and to use the information criteria for the alternating structure, although both the thresholds and the information criteria outperform the competing methods in this case. With respect to the computational cost, our method (any variant of it) is computationally faster than any other competitor and this can be seen in the last column of the tables.

### 4.2. Task based fMRI results

We now present the results of our method, CCID, on the task based fMRI data set described in Section 3.4; the results for SBS, Factor and BO are in the Appendix. For this data set, we apply CCID to each subject separately, that is, we do not align the change-points in any way across the subjects. Figure 7 shows the detected change-points for the 4 ROI and heart rate data using CCID with threshold *L*_∞_ (top) and information criterion *L*_∞_*IC* (bottom) using a minimum distance between change-points of *δ* = 1 (left) and *δ* = 40 (right). The blue vertical lines indicate the times of the showing and of the removal of the visual cues. In some experiments, the number of change-points itself could be the objective of the study. Hence, a minimum distance *δ* = 1 could be utilized. However, in other cases, researchers often would like to estimate a partition specific brain network or the undirected graph between each pair of detected change-points. This helps visualize the FC network in a more precise fashion. To this end, we used a minimum distance *δ* = 40 (however, this is at the discretion of the researcher). For more details on the precise location of the change-points see Table 11 in the Appendix. The y-axis depicts the subject number while the x-axis shows the change-point location times. As we have already stated, one of the main advantages of CCID over previous change-point methods is that CCID can detect change-points that are very close to one another and is not limited by a minimum distance between change-points input. Hence, CCID is not only able to detect change-points that are in the neighborhood of time point 60, which corresponds directly to the presentation of the first visual cue specifying the topic of the speech but also at the removal of said cue at time point 67.5 (see subjects 5, 12, 16 and 17 in the *L*_∞_ with minimum distance = 1 in Figure 7, top left). Likewise for time points 130 and 137.5, the second visual cue stating that the participants would in fact not have to give the presentation to the expert panel of judges after the conclusion of the scanning session was revealed (see subjects 2, 10, 19 in the *L*_∞_ with minimum distance = 1 in Figure 7, top left). The change-points occurring prior to the first visual cue may be related to anticipation of the speech topic, while change-points occurring during cues may be due to the different modes of anxiety as subjects silently prepare their speech. Finally, change-points occurring after the second visual cue may be due to the different modes of rest as subjects silently come to the end of the experiment. This pattern of change points is also seen for CCID with *L*_∞_ (Figure 7, bottom left). CCID is the only existing method (to the best of our knowledge) capable of finding change points this close to one another.

**Figure 7:**
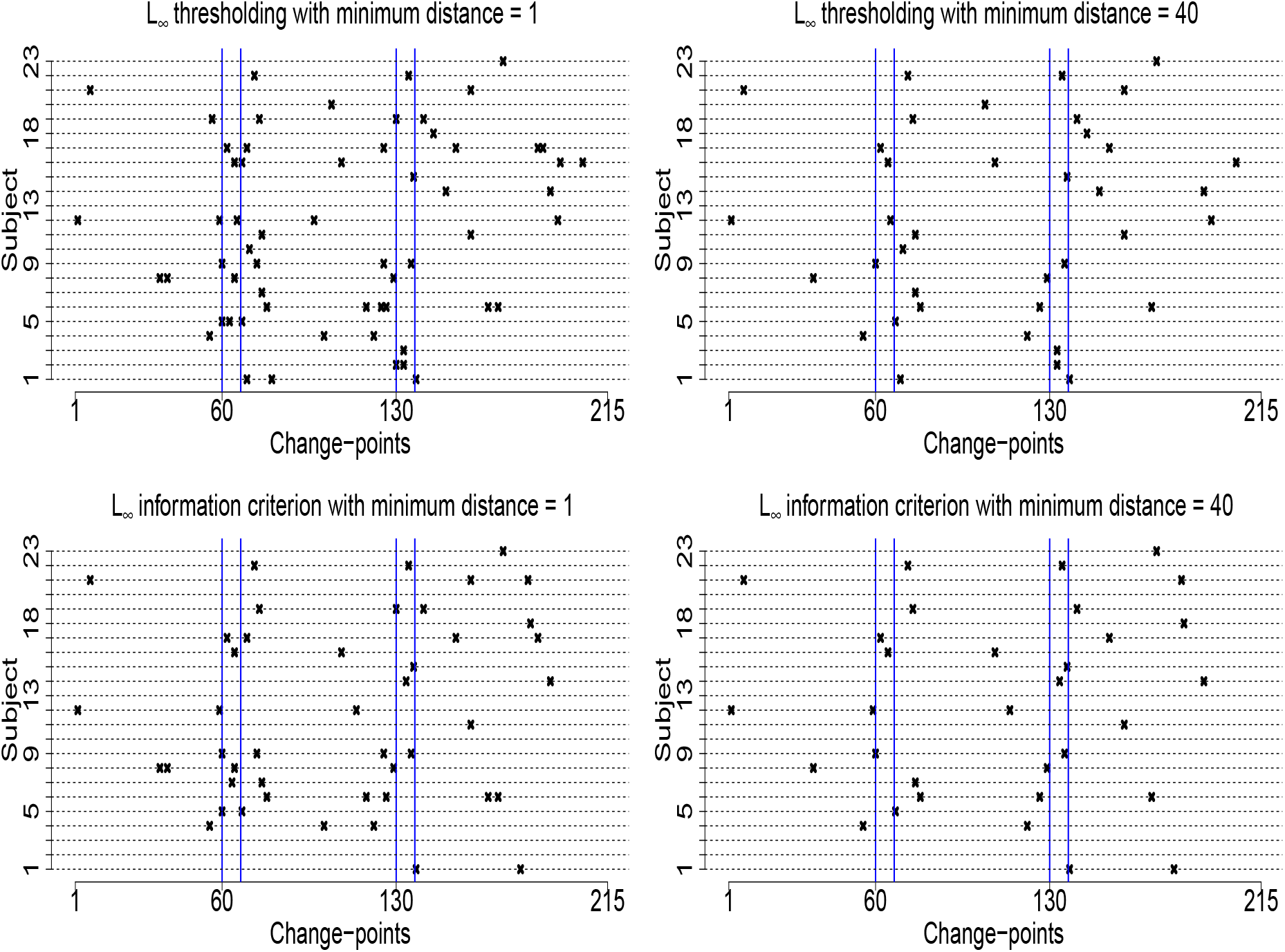
The detected change-points for the 4 ROI and heart rate data using CCID with threshold *L*_∞_ (top) and information criterion *L*_∞_*IC* (bottom) using a minimum distance between change-points of *δ* = 1 (left) and *δ* = 40 (right). The y-axis depicts the subject number while the x-axis shows the change-point location times. The blue vertical lines indicate the times of the showing and of the removal of the visual cues.

As mentioned above, often researchers would like to estimate a partition specific brain network or the undirected graph between each pair of detected change-points. This helps visualize the FC network in a more precise fashion. Hence, Figure 7 (right) shows the results where a minimum distance of *δ* = 40 was used. In this case, every subject has either two or three change-points with each subject having a change-point in the neighborhood of time point 60 (the first visual cue) and time point 130 (the second visual cue).

In addition, Figure 8 shows the detected change-points for the 4 ROI and heart rate data using CCID with threshold *L*_2_ (top) and information criterion *L*_2_*IC* (bottom) using a minimum distance between change-points of *δ* = 1 (left) and *δ* = 40 (right). All of the conclusions that we described above in Figure 7 are also true for these methods. The only difference is that CCID with threshold *L*_2_ (top) and information criterion *L*_2_*IC* detect more change points for both *δ* = 1, 40.

**Figure 8:**
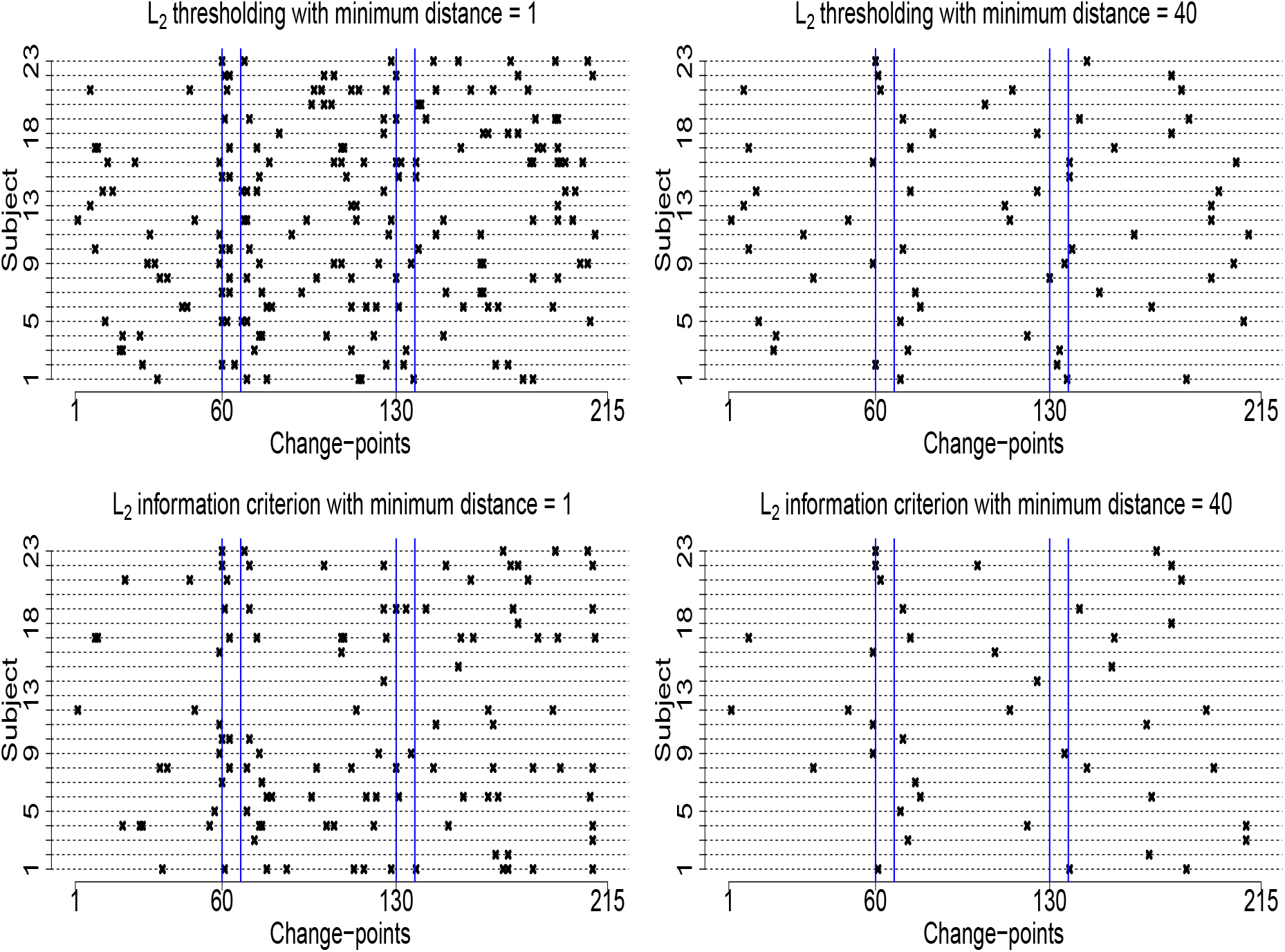
The detected change-points for the 4 ROI and heart rate data using CCID with threshold *L*_2_ (top) and information criterion *L*_2_*IC* (bottom) using a minimum distance between change-points of *δ* = 1 (left) and *δ* = 40 (right). The y-axis depicts the subject number while the x-axis shows the change-point location times. The blue vertical lines indicate the times of the showing and of the removal of the visual cues.

Figure 9 shows the density plots for the detected change-points in the 4 ROI and heart rate data using all variants of CCID across all 23 subjects using a minimum distance between change points of *δ* = 40. Again, the blue vertical lines indicate the times of the showing and of the removal of the visual cues. We can see that for all combinations of CCID there are clear peaks around time points 60-67.5 and 130-137.5, the times of the showing and the removal of the visual cues. There is consistency across the combinations although the thresholding provide the most distinct peaks.

**Figure 9:**
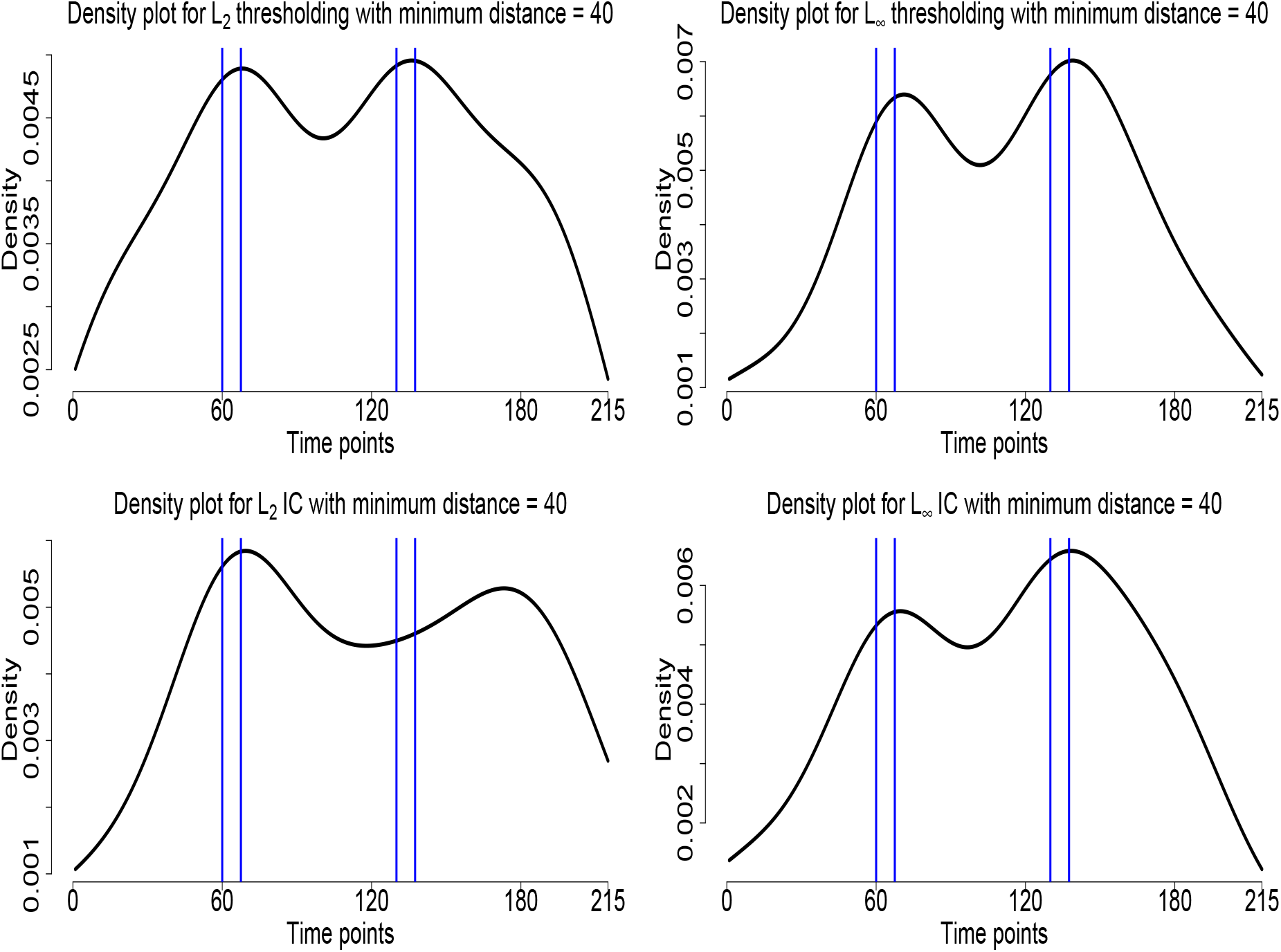
The density plots for the detected change-points in the 4 ROI and heart rate data using CCID across all 23 subjects using *δ* = 40. **Top row**: CCID with *L*_2_ thresholding (left) and CCID with *L*_∞_ thresholding (right) **Bottom row**: CCID with *L*_2_ Information-Criterion (left) and CCID with *L*_∞_ Information-Criterion (right).

For comparison, Figure 14 (in the Appendix) shows the detected change-points for the 4 ROI and heart rate data using the competing methods: SBS, Factor, and BO. Here, SBS and Factor idiosyncratic find at most one change-point for each subject and in the vast majority of cases they find no change-point at all. To be more precise, SBS does not find any change-points in 18 out of 23 subjects and Factor idiosyncratic in 16 subjects. None of the detected change-points by SBS are in the neighborhood of time point 60, which corresponds directly to the presentation of the first visual cue specifying the topic of the speech and the removal of said cue at time point 67.5 (135 seconds). Only one change-point is near time point 130 (subject 15), the second visual cue stating that the participants would in fact not have to give the presentation to the expert panel of judges after the conclusion of the scanning session was revealed. The other change-points may be due to the different modes of anxiety as subjects silently prepare their speech. The BO method detects two change-points in 4 subjects, one change-point in 5 subjects and no change-points in 14 subjects. Factor common detects three change-points in 1 subject, two change-points in also 1 subject, one change-point in 6 subjects and no change-points in 15 out of 23 subjects.

As mentioned before we reduced the number of change-points that CCID detected by specifying a minimum distance between change points. There are two other procedures available to CCID to reduce the number of detected change-points. First, the thresholds in the *L*_∞_ and *L*_2_ approaches (see Section 3.1 for a discussion) and the penalties for the *L*_∞_ and *L*_2_ information criteria approaches can be increased. Second, CCID provides the solution path for the *L*_∞_ and *L*_2_ thresholding approaches, which is an ordering of the importance of the detected change-points in descending order (see Section 2.6 for more details). Hence, a certain number of change-points could be specified using this procedure. Figure 10 shows the density plots for the detected change-points in the 4 ROI and heart rate data across all 23 subjects using a minimum distance between change points of *δ* = 40, using the solution path method for the *L*_∞_ and *L*_2_ thresholding approaches, where we assume two change-points per channel. Both of these density plots show a good behavior with a clear bimodal structure that peak close to the first and second visual cues. The results from this procedure are comparable to the original robust choices we used in Figure 9. Of course, specifying a certain number of change-points would benefit from input from the domain knowledge experts. These procedures show the flexibility of CCID and allows the researchers alternative strategies to reduce the number of detected change-points.

**Figure 10:**
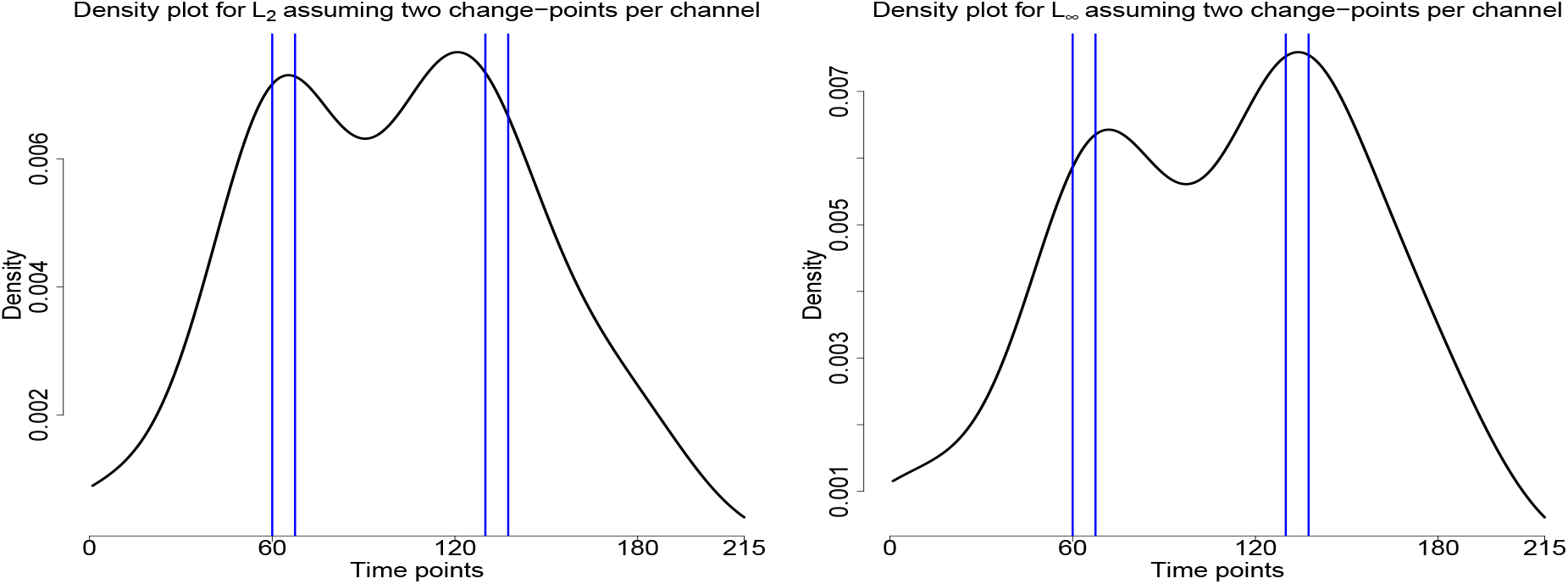
The density plots for the detected change-points in the 4 ROI and heart rate data using CCID across all 23 subjects using *δ* = 40 using the solution path method for the *L*_∞_ and *L*_2_ thresholding approaches, where we assume two change-points per channel. CCID with *L*_2_ thresholding (left) and CCID with *L*_∞_ thresholding (right)

The FC undirected graphs, estimated by SCAD and BIC (Section 2.8), for subjects 4, 6, 8, 9 and 11 between the detected change-points for CCID.*L*_∞_ are displayed in Figure 11. Black (red) edges infer positive (negative) connectivity, and the strength of connection between the regions is directly related to the thickness of the edges, that is, the thicker the edge the stronger the connection. The connectivity structure in the undirected graphs for each subject is quite different, however, the connection between the anterior mPFC and VMPFC is consistent across all subjects. From the graphs it is also evident that HR is more connected to the other ROIs during the resting periods, while during the second partition, when the speech topic is presented and the participants begin to silently prepare their speech, there is less connections between HR and the other ROIs. Perhaps this is due to the anticipation stage of the experiment being more stressful than the preparation stage.

**Figure 11:**
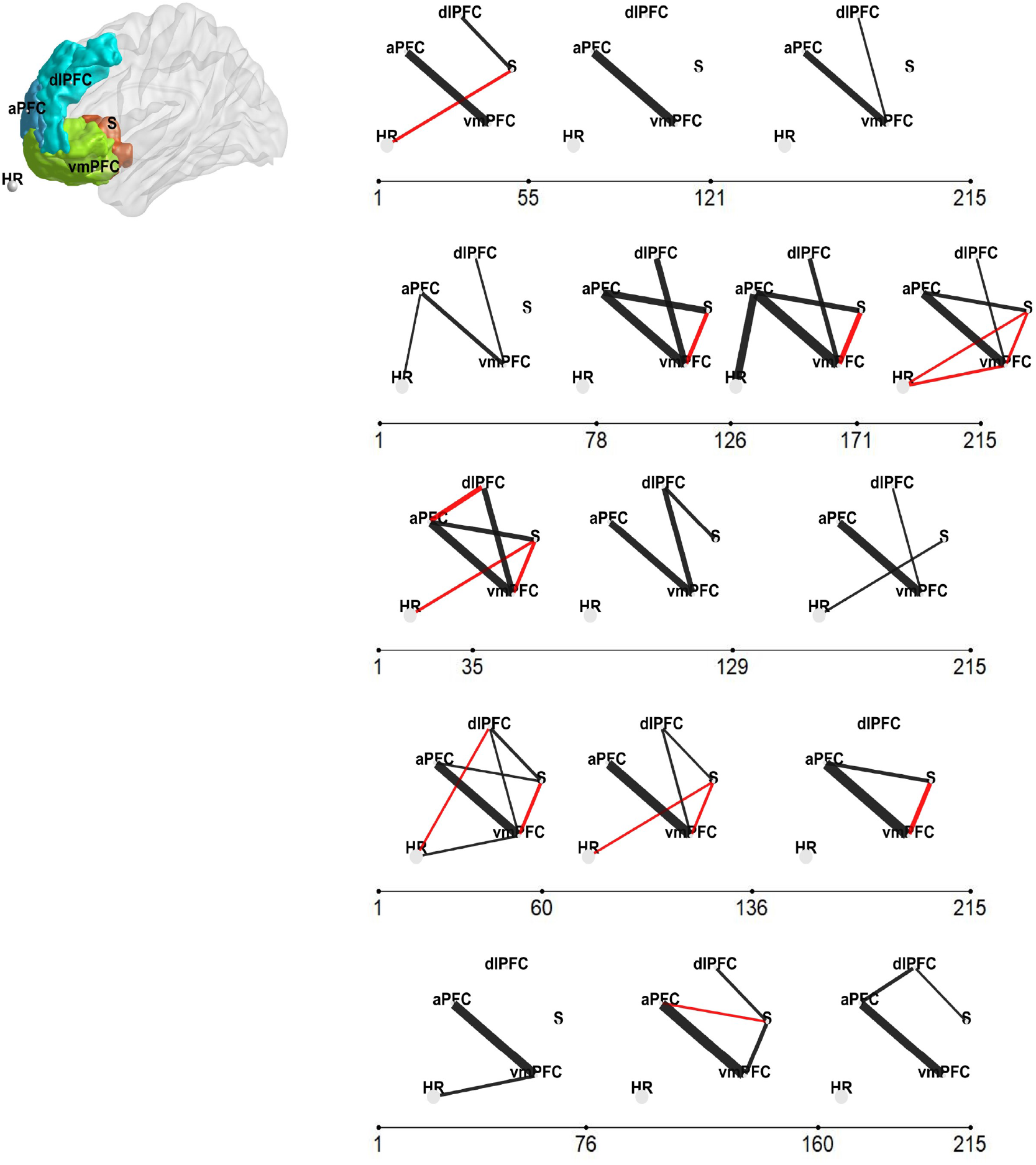
The four-ROI and heart rate data set (task based fMRI data set) the regions are : (1) VMPFC, (2) anterior mPFC, (3) Striatum/pallidum, and (4) DLPFC/IFJ and 5) heart rate (left). The FC undirected graphs, estimated by SCAD and BIC, for subjects 4, 6, 8, 9 and 11 (each row corresponds to a particular subject) between the detected change-points for CCID.*L*_∞_ (right). Black (red) edges infer positive (negative) connectivity, and the strength of connection between the regions is directly related to the thickness of the edges, that is, the thicker the edge the stronger the connection.

### 4.3. Resting state fMRI results

Evidence of the non-stationary behavior of brain activity time series has been observed not only in task based fMRI studies, but also prominently in resting-state data ((Delamillieure et al., 2010; Doucet et al., 2012; Ondrus et al., 2021; Xiong & Cribben, 2021). In these experiment mental activity is unconstrained. Figure 12 shows the detected change-points for the resting state fMRI data using CCID with threshold *L*_∞_ (top) and information criterion *L*_∞_*IC* (bottom) using a minimum distance between change-points of *δ* = 1 (left) and *δ* = 40 (right). For more details on the precise location of the change-points see Table 12 in the Appendix. The y-axis depicts the subject number while the x-axis shows the change-point location times. Where the minimum distance between change-points is not restricted, every subject has at least three change-points in their second order structure structure with the maximum number of change-points being 30 (subject 1). The results indicate that, not only does the number of second-order structure change-points differ across subjects, but the location of the change-points is also variable. In addition, some subjects remain in states for short periods whereas others transition more quickly. Hence, CCID has a major advantage over moving window-type methods as we do not have to choose the window length, which can have significant consequences on the estimated brain networks. As we have already stated for the Anxiety fMRI data, one of the main advantages of CCID over previous change-point methods is that CCID allows for detection in the presence of frequent changes of possibly small magnitudes. Hence, for resting state data, where the subjects are unconstrained, CCID is very suitable. No previous method has been able to detect as many change-points and also change-points that are very close to one another. In some studies, the number of change-points itself could be the objective of the study or an input into a model for disease classification. Hence, a small minimum distance could be utilized. However, in other cases, often researchers would like to estimate a partition specific brain network or the undirected graph between each pair of detected change-points for visualizing the FC network. Hence, Figure 12 (right) shows the results where a minimum distance of *δ* = 40 was used. In this case, every subject has up to 5 change-points with a minimum of 3 change-points.

**Figure 12:**
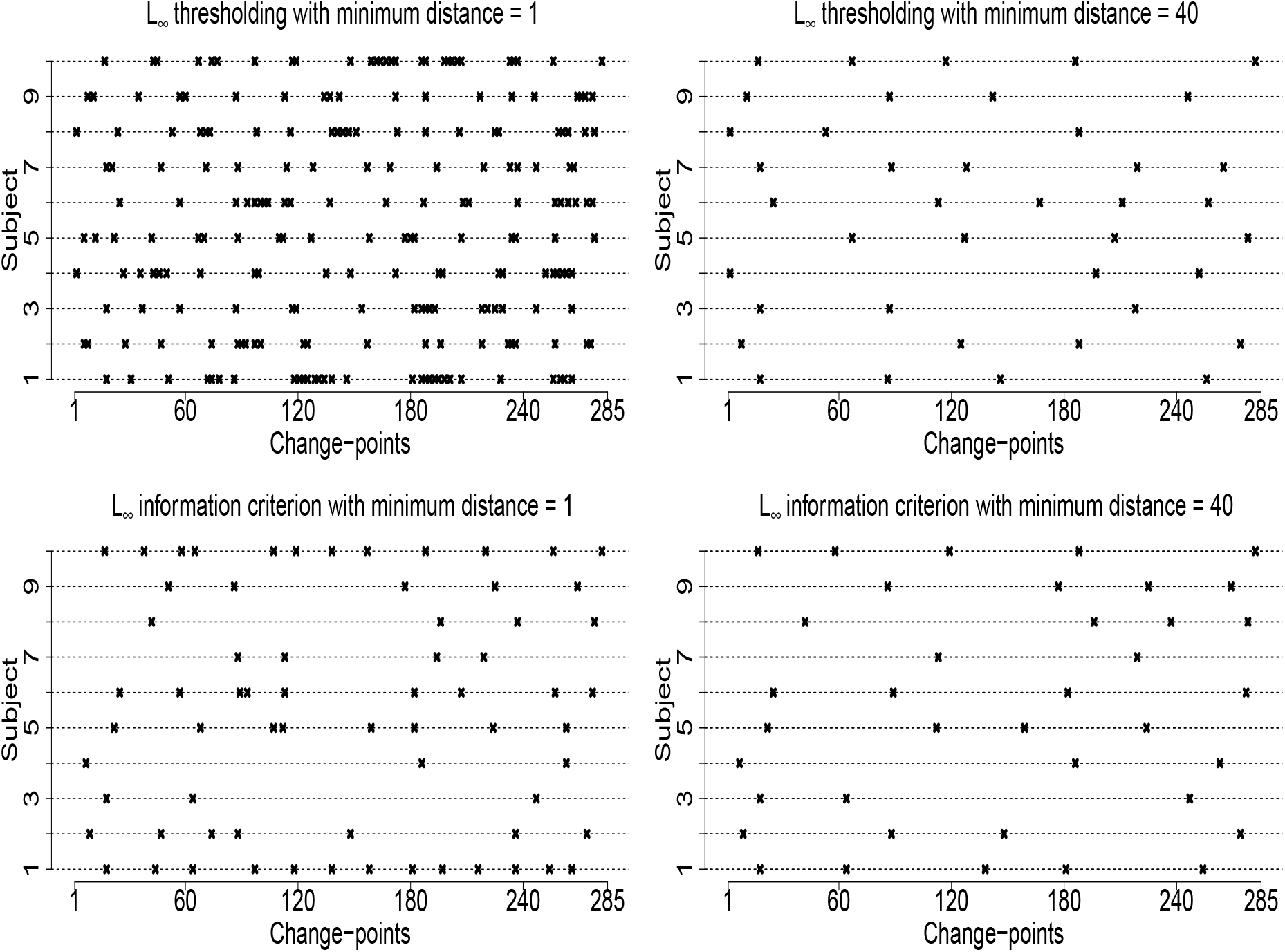
The detected change-points for the resting state fMRI data using CCID with threshold *L*_∞_ (top) and information criterion *L*_∞_*IC* (bottom) using a minimum distance between change-points of *δ* = 1 (left) and *δ* = 40 (right). The y-axis depicts the subject number while the x-axis shows the change-point location times.

Figure 13 shows the detected change-points for the resting state fMRI data using CCID with threshold *L*_2_ (top) and information criterion *L*_2_*IC* (bottom) using a minimum distance between change-points of *δ* = 1 (left) and *δ* = 40 (right). All of the conclusions that we described above in Figure 12 are similar for these methods. The only difference is that CCID with threshold *L*_2_ (top) and information criterion *L*_2_*IC* detect less change points for both *δ* = 1, 40 and are more spread out for *δ* = 1.

**Figure 13:**
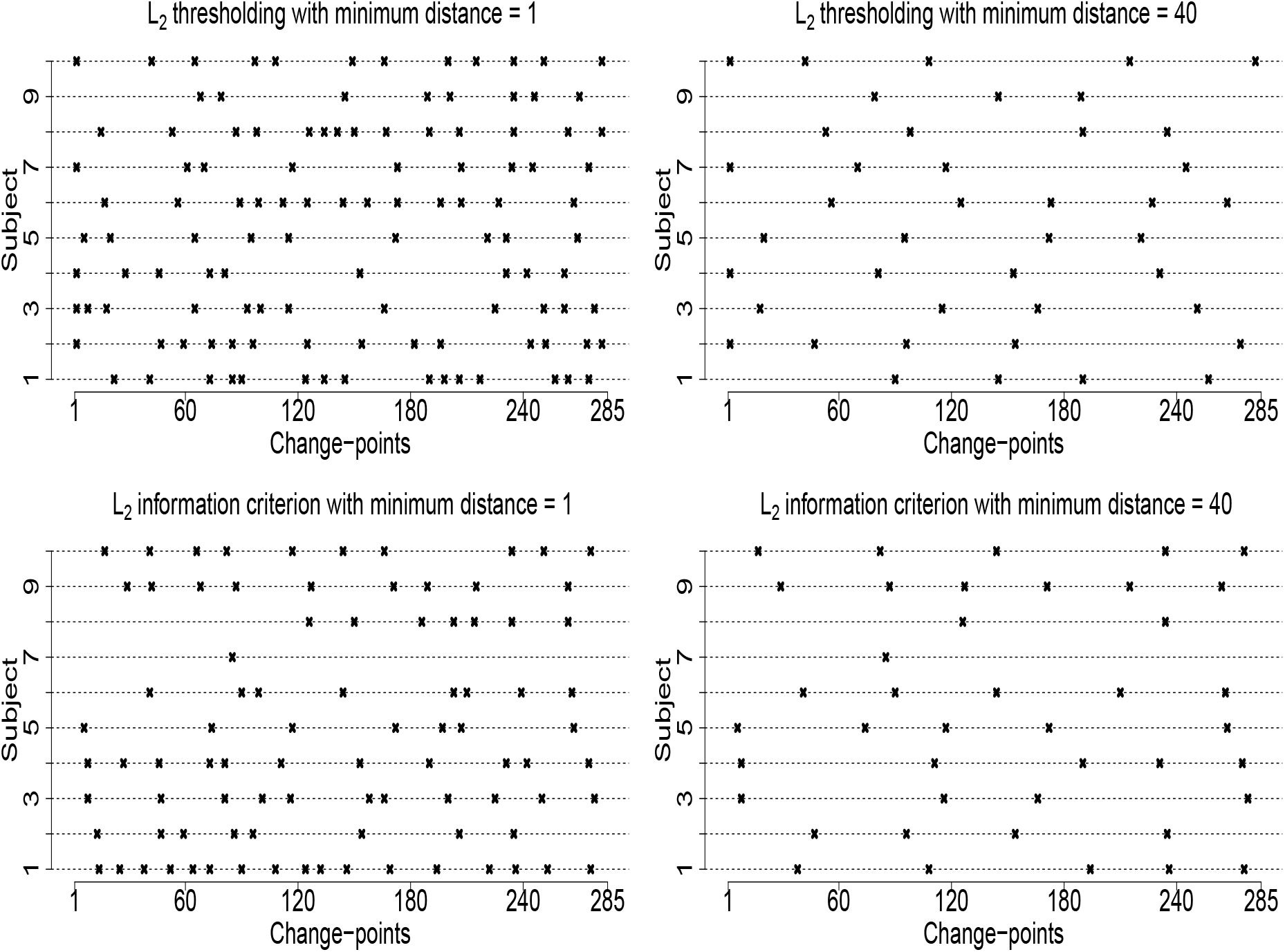
The detected change-points for the resting state fMRI data using CCID with threshold *L*_2_ (top) and information criterion *L*_2_*IC* (bottom) using a minimum distance between change-points of *δ* = 1 (left) and *δ* = 40 (right). The y-axis depicts the subject number while the x-axis shows the change-point location times.

For comparison, Figure 15 (see Appendix) shows the results for all the competing methods for the resting state fMRI data. SBS finds either one or two change-points for each subject, Factor common finds at most one change-point, while Factor idiosyncratic does not find any change-points for all subjects. The BO method finds multiple change-points for all subjects, apart from Subject 4, where this method detects one change-point. Although we are not certain of the number of change-points for resting state fMRI, as the subject is unconstrained, we expect subjects to traverse several functional states across the experiment and previous studies Ofori-Boateng et al. (2020) have located multiple change-points, something which agrees with the results of CCID and BO, but not with those of SBS and Factor. However, due to the liberal Type I errors for BO (also seen in Ofori-Boateng et al., 2020), many of BO’s change-points may be due to false positive change-points.

## 5. Discussion

### 5.1. Impact of autocorrelation

Our CCID model first constructs appropriate wavelet-based local periodogram sequences from the original multivariate time series, **X**_*t,T*_. To detect change-points, it uses a scaled CUSUM statistic on the transformed data. The optimization of model selection criteria provides an alternative approach to the thresholding methods for pruning the set of change-points, which we assume have been overestimated. A key element in the construction of the information criteria is the likelihood function for our transformed data. Due to the unknown dependence structure in our transformed data (wavelet-based local periodogram sequences), the construction of the exact likelihood is very difficult. Hence, we work instead on an approximation of the likelihood, the pseudo-likelihood, where the data are (incorrectly) taken to be independent. This is a common approach. For example, Zhang et al. (2014) assume independence among the temporal segments, which is critical to the estimation of the posterior distribution of their Dynamic Bayesian Variable Partition models. However, in our simulations and fMRI data analyses, we show that CCID with the model selection criteria (and therefore the pseudo-likelihood) performs well for both independent data and data with a moderate amount of autocorrelation. In cases where the autocorrelation in the transformed data is very high, we propose the following variants of our algorithm that can be employed to achieve better practical performance.

#### Subsampling

Suppose we have *d* periodograms after the wavelet transformation is applied on the original *p* time series, then we can subsample from the periodograms and apply CCID to the data sequences created. For conceptual simplicity, allow us to explain this subsampling variant through an example where the length of each periodogram, 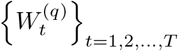 for *q* = 1, 2, *…, d*, is *T* = 1000. In this example, we subsample every *s* = 4 data points; of course different values of *s* can be taken and our specific choice is made only for the sake of explaining this subsampling variant. We note that the lower the value of *s*, the more autocorrelation in our new (subsampled) data but also the more accurate the estimated change-point locations. In our example of *T* = 1000 and *s* = 4, the following *s* data sequences for each periodogram are created:

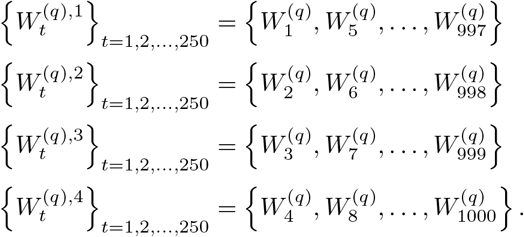

After completing this step, we apply, for each *j* = 1, 2, 3, 4 separately, CCID to the multivariate time series

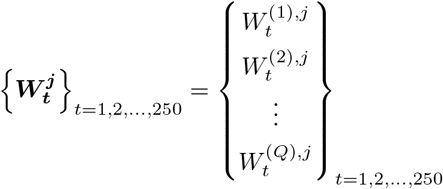

in order for each *j* = 1, 2, …, *s*, to detect the change-points for 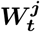; this set of change-points is denoted by 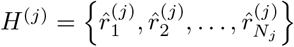. We define

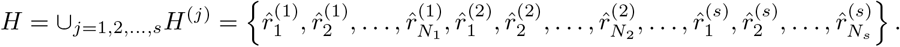

Some of the elements in the set *H* can of course be identical. The next step is to apply a majority voting rule, in the sense that we only keep those values that appear at least *η* times, where *η* ∈ {1, 2, …, *s*}. The value of *η* can be decided upon using *a priori* knowledge of the particular neuroimaging data set. Once the values that appear at least *η* times in *H* are extracted, the change-points are then transformed to represent the change-point locations with respect to the original periodograms. For example if the 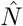 values 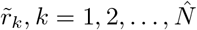, appear more than *η* times in *H*, then the estimated change-point locations with respect to the original data are 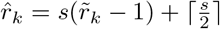.

#### Pre-averaging

Pre-averaging is a strategy where data are averaged over short time periods. Here, for a given scale number *s* and *d* periodograms 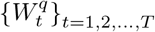 and *q* = 1, 2, …, *d*, we let *S*\* = ⌈*T/s*⌉ and we define now for each *q*,

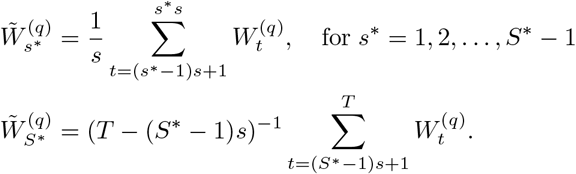

The next step is to apply CCID on

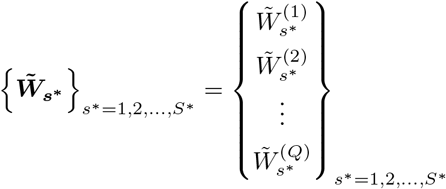

to obtain the estimated change-points, namely 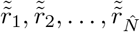, in increasing order. To estimate the original locations of the change-points we define 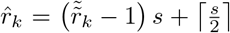, 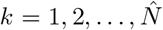. There is a trade-off in the choice of the scaling parameter *s*. If the value of *s* is large, then we assume that our new (pre-averaged) data are less autocorrelated but the amount of pre-processing obviously increases, which results in loss in power (due to less data) and also possibly in lack in the accuracy of the detected change-point locations.

### 5.2. CCID constants and recommendations

For CCID, we provided results for the *L*_2_ and the *L*_∞_ threshold approaches and the *L*_2_ and the *L*_∞_ information criteria. Our choice of constants for the approaches (*c*_1_ for the *L*_2_ threshold; *c*_2_ for the *L*_∞_ threshold and *α* for the information criteria) provided a balance between specificity and sensitivity in all signals examined in our large scale simulation study and others not included. Hence, all constants have been calibrated on higher dimensions. However, it is important that CCID remains robust to alternative choices to these parameters and the practitioner has the option to obtain more change-points by decreasing these threshold value.

In our numerical experience, it appears that the *L*_∞_ threshold performs best for low dimensional cases (*P* < 5) with the *L*_2_ threshold over detecting the change points. The reason for this is that the *L*_∞_ threshold boils down to just one cross covariance while the *L*_2_ threshold aggregates over all series. Hence, the *L*_2_ threshold works better for small changes across many series.

## 6. Conclusion

In this study, we developed a new method, called Cross-covariance isolate detect (CCID), to detect multiple change-points in the second-order (cross-covariance or network) structure of multivariate (possibly) high-dimensional time series. We assumed that the number and location of the change-points are unknown a priori. CCID first converted our multivariate time series into periodograms and cross-periodograms. To detect the change-points CCID used a scaled CUSUM statistic and aggregated across the multivariate time series by adding only those that pass a certain threshold. CCID adapted the Isolate Detect principle to find the multiple change-points. ID works by first isolating each of the true change-points within subintervals and then secondly detecting them within these subintervals. We showed using an extensive simulation study that CCID generally had higher statistical power compared to existing methodologies. Our simulation study also showed that CCID performs better than existing methods in various challenging scenarios where the subject alternates between task and rest. In comparison, binary segmentation (used by all previous change-point methods in the neuroscience literature) failed to perform as well in this scenario. In addition, using empirical task based and resting-state fMRI data, CCID provided significant superior results in terms of change-point detection to the existing methods.

We showed that CCID is also more computationally efficient than the competing methods and in many settings is at least an order of magnitude faster than them. Computational costs can be challenging at higher dimensions as the first step of our method requires the transformation of the multivariate time series to periodograms and cross-periodograms. While in theory a large number of ROIs is possible, in practice we may need to make modifications to our proposed method to accommodate a larger number of ROIs (whole brain dynamics or voxel time series). We could take advantage of parallel computing in order to speed up the method.

CCID is novel as it allows for detection in the presence of frequent changes of possibly small magnitudes, can assign change-points to one or multiple brain regions, and is computationally very fast. We also proposed new information criteria for CCID to identify the change-points. While CCID was applied to task based and resting-state fMRI data in this work, it could seamlessly be applied to Electroencephalography (EEG) or Magnetoencephalography (MEG), or electrocorticography (ECoG) data. The outputs of CCID could also be used as an input into a classification model for predicting brain disorders. Furthermore, while previous studies have attempted to find differences between controls and patients with brain disorders such as Alzheimer’s disease (Hart et al., 2018), understand FC changes in cerebral palsy before and after treatment (Bakhtiari et al., 2017), and understand the role of speech production in reading (Cummine et al., 2016) using static FC networks, it would be very interesting to see if dynamic FC estimated via CCID provides more detailed descriptions of these complicated processes. To conclude, CCID pertains to a general setting and can also be used in a variety of situations where one wishes to study the evolution of a high dimensional network over time.

## 7. Software

The authors have released an open source R package by the name of **ccid**. The package can be downloaded from CRAN.

## 8. Acknowledgements

The work of I. Cribben was supported by the Natural Sciences and Engineering Research Council (Canada) grant RGPIN-2018-06638 and the Xerox Faculty Fellowship, Alberta School of Business.

## 9. Appendix

## A The results for the real data presented in tables

**Table 11:**
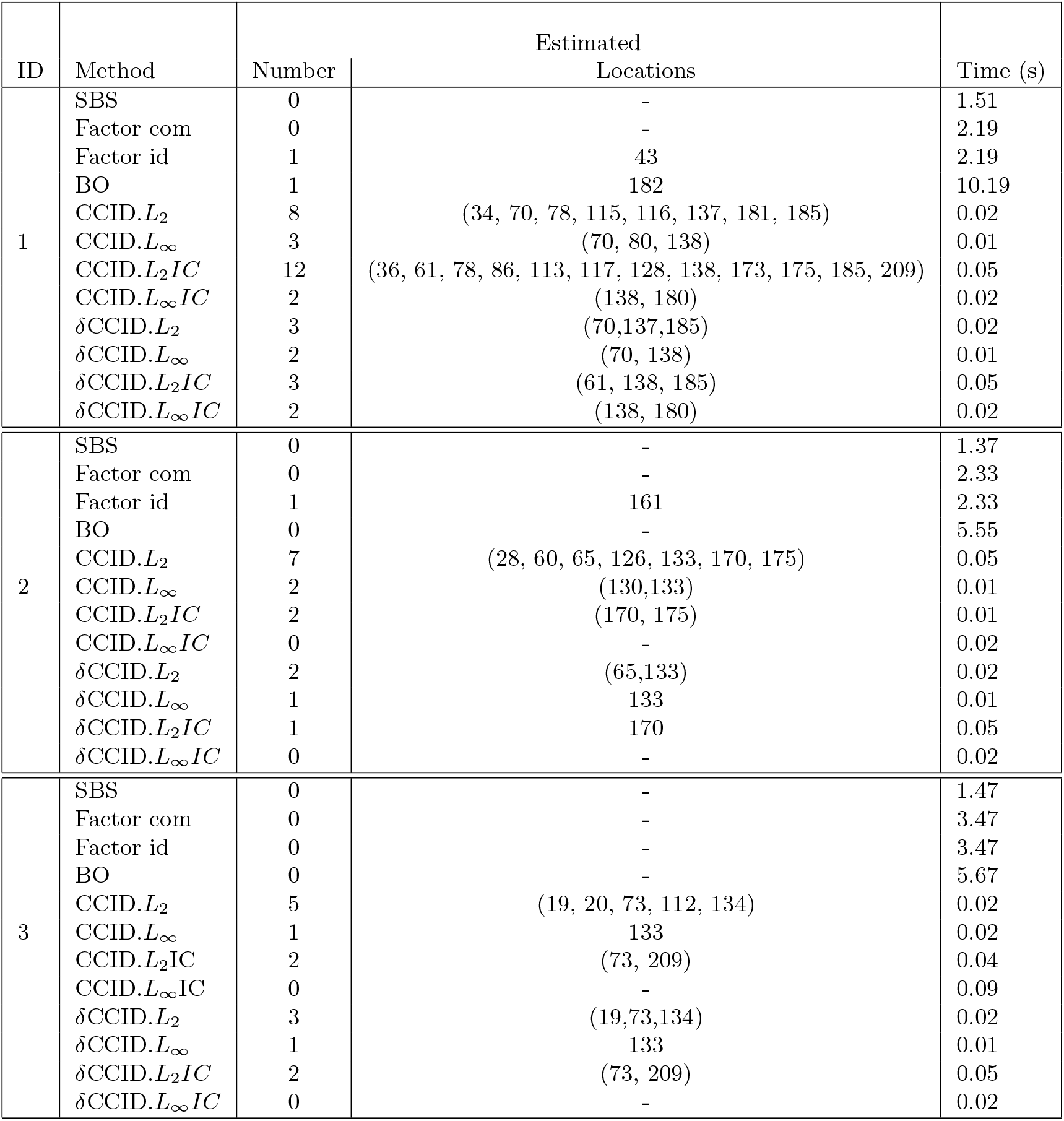
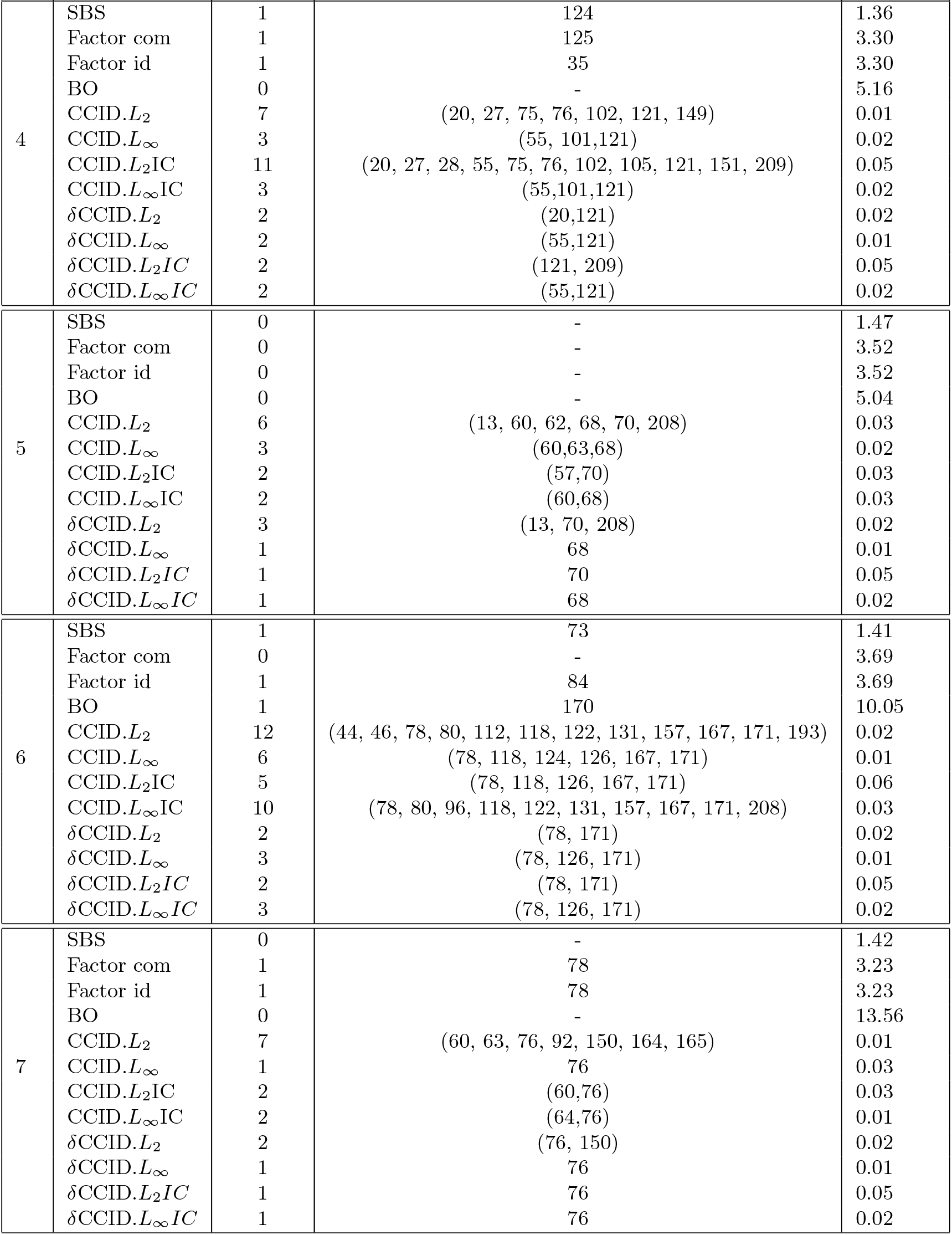
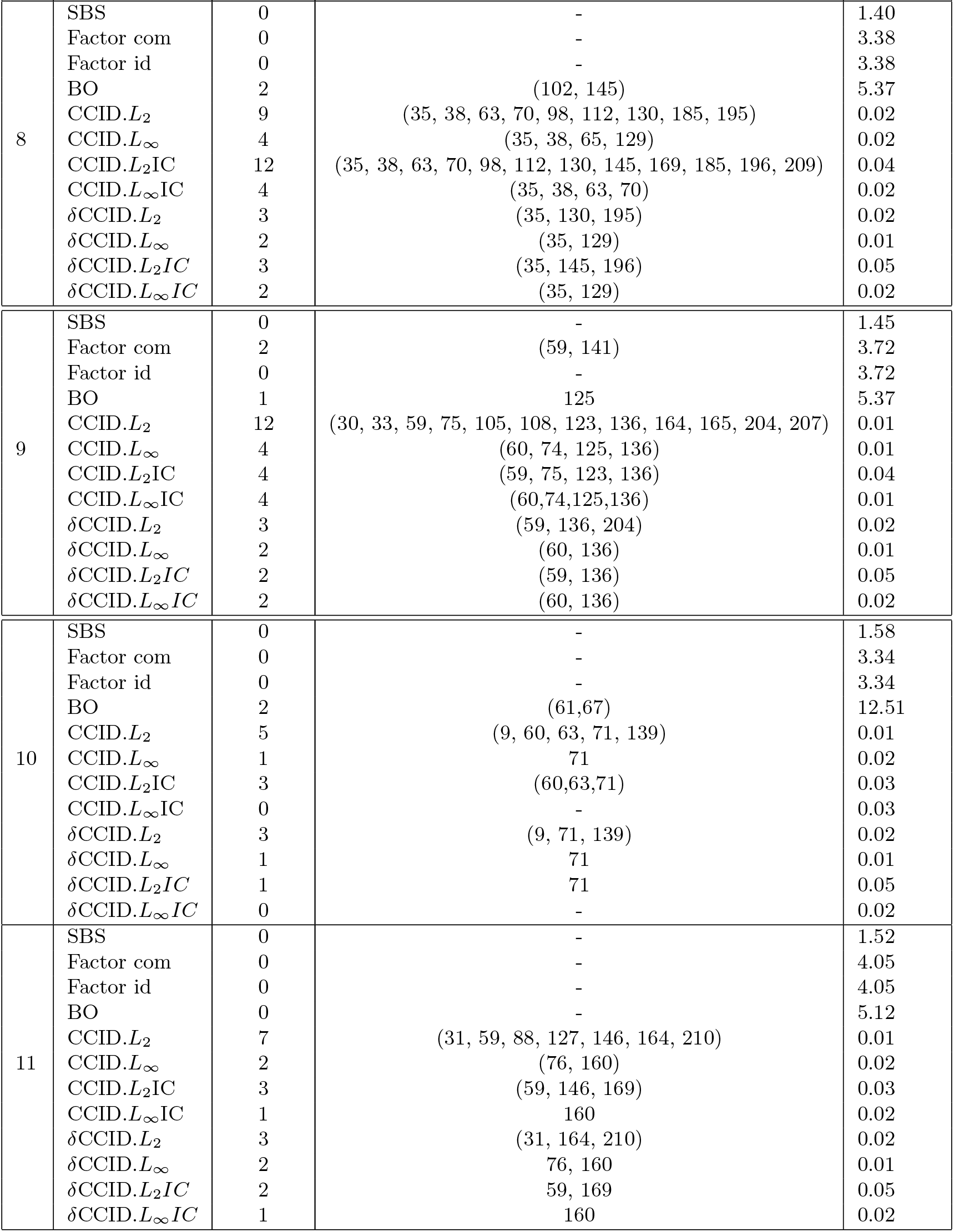
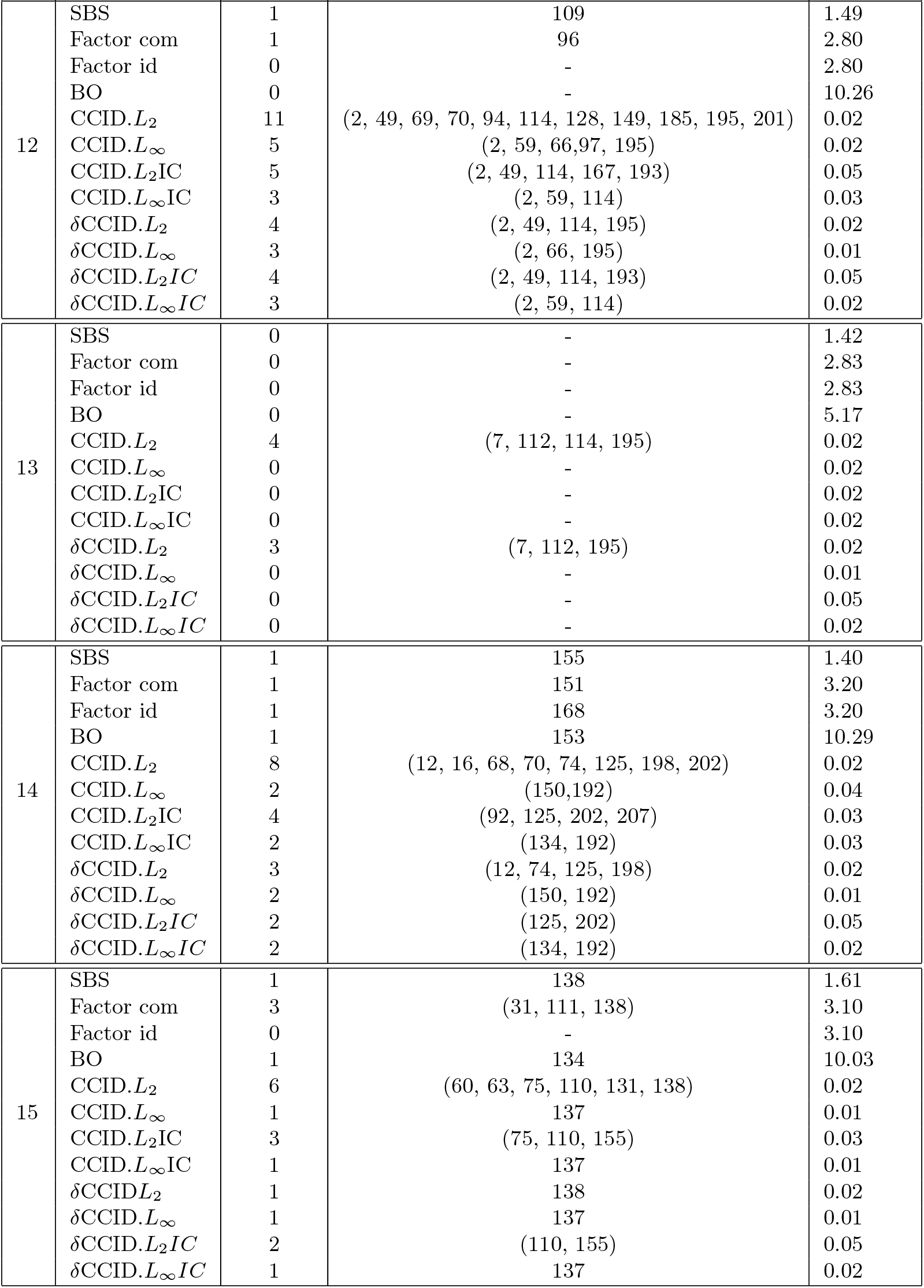
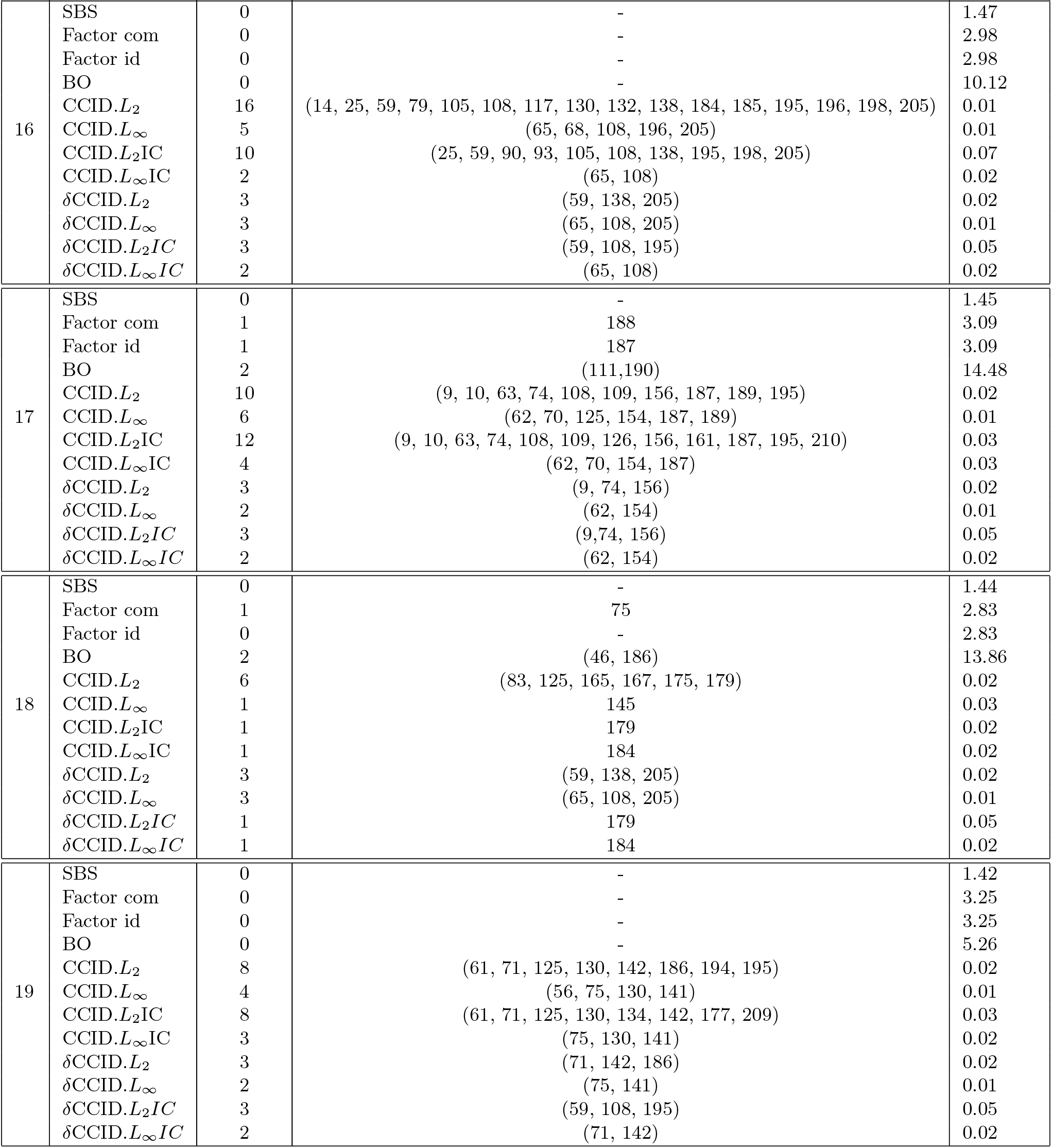
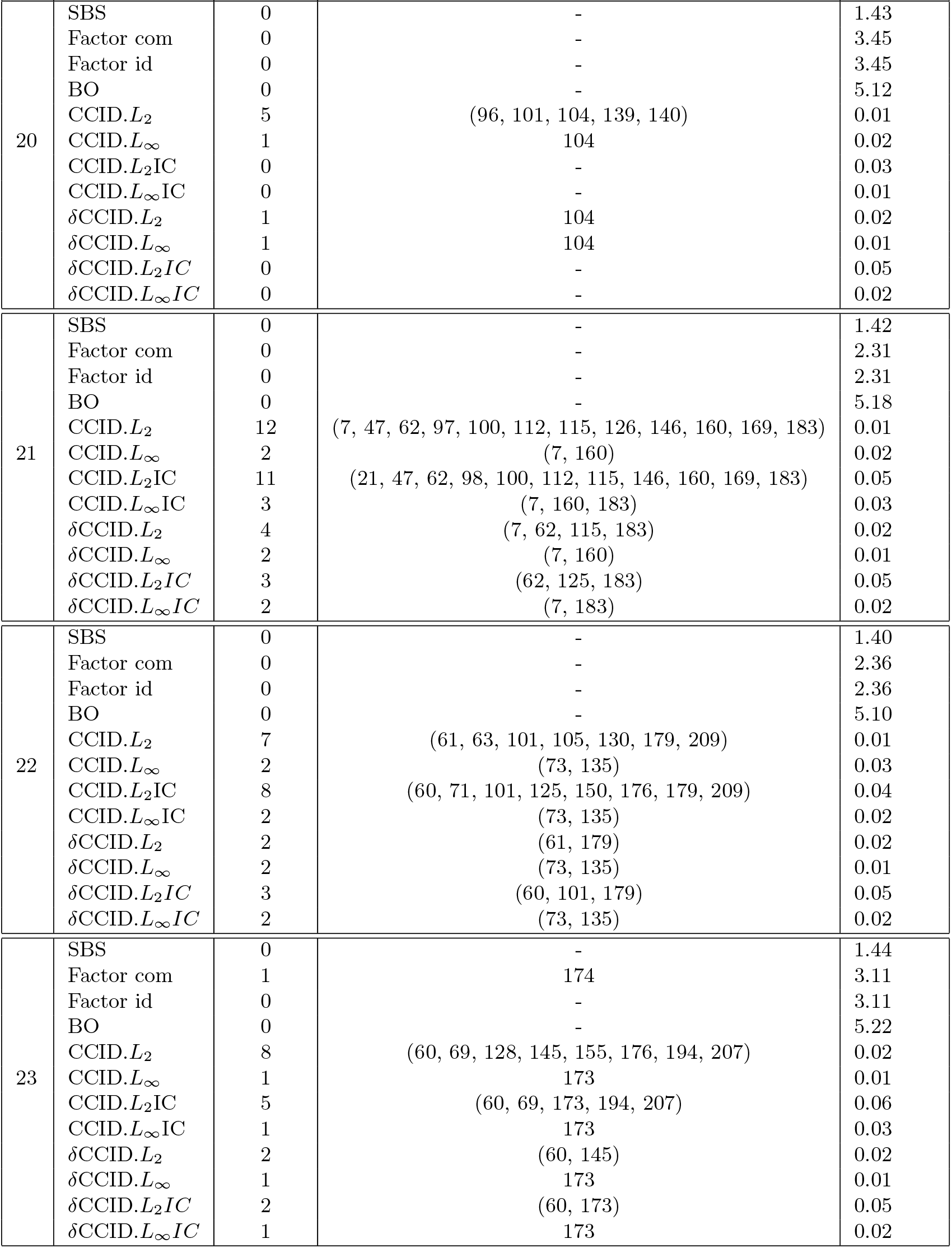
The estimated number and location of the change-points for the 4 ROIs and heart rate task based fMRI data set for Sparsified Binary Segmentation (SBS), Factor with common components (Factor com), Factor with idiosyncratic components (Factor id), Barnett & Onnela (2016) (BO), and Cross-covariance isolate detect with the *L*_2_ threshold (CCID.*L*_2_), the *L*_∞_ threshold (CCID.*L*_∞_), the *L*_2_ information criterion (CCID.*L*_2_*IC*) and the *L*_∞_ information criterion (CCID.*L*_∞_*IC*). A *δ* in the name of a method indicates a minimum distance between change-points of 40 time points was enforced. The computational times for each method are also provided.

| ID | Method | Number | Estimated Locations | Time (s) |
| --- | --- | --- | --- | --- |
| 1 | SBS | 0 | - | 1.51 |
|  | Factor com | 0 | - | 2.19 |
|  | Factor id | 1 | 43 | 2.19 |
|  | BO | 1 | 182 | 10.19 |
| | CCID.$L_2$ | 8 | (34, 70, 78, 115, 116, 137, 181, 185) | 0.02 |
| | CCID.$L_\infty$ | 3 | (70, 80, 138) | 0.01 |
| | CCID.$L_2IC$ | 12 | (36, 61, 78, 86, 113, 117, 128, 138, 173, 175, 185, 209) | 0.05 |
| | CCID.$L_\infty IC$ | 2 | (138, 180) | 0.02 |
| | δCCID.$L_2$ | 3 | (70,137,185) | 0.02 |
| | δCCID.$L_\infty$ | 2 | (70, 138) | 0.01 |
| | δCCID.$L_2IC$ | 3 | (61, 138, 185) | 0.05 |
| | δCCID.$L_\infty IC$ | 2 | (138, 180) | 0.02 |
| 2 | SBS | 0 | - | 1.37 |
|  | Factor com | 0 | - | 2.33 |
|  | Factor id | 1 | 161 | 2.33 |
|  | BO | 0 | - | 5.55 |
| | CCID.$L_2$ | 7 | (28, 60, 65, 126, 133, 170, 175) | 0.05 |
| | CCID.$L_\infty$ | 2 | (130,133) | 0.01 |
| | CCID.$L_2IC$ | 2 | (170, 175) | 0.01 |
| | CCID.$L_\infty IC$ | 0 | - | 0.02 |
| | δCCID.$L_2$ | 2 | (65,133) | 0.02 |
| | δCCID.$L_\infty$ | 1 | 133 | 0.01 |
| | δCCID.$L_2IC$ | 1 | 170 | 0.05 |
| | δCCID.$L_\infty IC$ | 0 | - | 0.02 |
| 3 | SBS | 0 | - | 1.47 |
|  | Factor com | 0 | - | 3.47 |
|  | Factor id | 0 | - | 3.47 |
|  | BO | 0 | - | 5.67 |
| | CCID.$L_2$ | 5 | (19, 20, 73, 112, 134) | 0.02 |
| | CCID.$L_\infty$ | 1 | 133 | 0.02 |
| | CCID.$L_2IC$ | 2 | (73, 209) | 0.04 |
| | CCID.$L_\infty IC$ | 0 | - | 0.09 |
| | δCCID.$L_2$ | 3 | (19,73,134) | 0.02 |
| | δCCID.$L_\infty$ | 1 | 133 | 0.01 |
| | δCCID.$L_2IC$ | 2 | (73, 209) | 0.05 |
| | δCCID.$L_\infty IC$ | 0 | - | 0.02 |
| 4 | SBS | 1 | 124 | 1.36 |
|  | Factor com | 1 | 125 | 3.30 |
|  | Factor id | 1 | 35 | 3.30 |
|  | BO | 0 | - | 5.16 |
| | CCID.$L_2$ | 7 | (20, 27, 75, 76, 102, 121, 149) | 0.01 |
| | CCID.$L_\infty$ | 3 | (55, 101, 121) | 0.02 |
| | CCID.$L_2$IC | 11 | (20, 27, 28, 55, 75, 76, 102, 105, 121, 151, 209) | 0.05 |
| | CCID.$L_\infty$IC | 3 | (55, 101, 121) | 0.02 |
| | δCCID.$L_2$ | 2 | (20, 121) | 0.02 |
| | δCCID.$L_\infty$ | 2 | (55, 121) | 0.01 |
| | δCCID.$L_2$IC | 2 | (121, 209) | 0.05 |
| | δCCID.$L_\infty$IC | 2 | (55, 121) | 0.02 |
| 5 | SBS | 0 | - | 1.47 |
|  | Factor com | 0 | - | 3.52 |
|  | Factor id | 0 | - | 3.52 |
|  | BO | 0 | - | 5.04 |
| | CCID.$L_2$ | 6 | (13, 60, 62, 68, 70, 208) | 0.03 |
| | CCID.$L_\infty$ | 3 | (60, 63, 68) | 0.02 |
| | CCID.$L_2$IC | 2 | (57, 70) | 0.03 |
| | CCID.$L_\infty$IC | 2 | (60, 68) | 0.03 |
| | δCCID.$L_2$ | 3 | (13, 70, 208) | 0.02 |
| | δCCID.$L_\infty$ | 1 | 68 | 0.01 |
| | δCCID.$L_2$IC | 1 | 70 | 0.05 |
| | δCCID.$L_\infty$IC | 1 | 68 | 0.02 |
| 6 | SBS | 1 | 73 | 1.41 |
|  | Factor com | 0 | - | 3.69 |
|  | Factor id | 1 | 84 | 3.69 |
|  | BO | 1 | 170 | 10.05 |
| | CCID.$L_2$ | 12 | (44, 46, 78, 80, 112, 118, 122, 131, 157, 167, 171, 193) | 0.02 |
| | CCID.$L_\infty$ | 6 | (78, 118, 124, 126, 167, 171) | 0.01 |
| | CCID.$L_2$IC | 5 | (78, 118, 126, 167, 171) | 0.06 |
| | CCID.$L_\infty$IC | 10 | (78, 80, 96, 118, 122, 131, 157, 167, 171, 208) | 0.03 |
| | δCCID.$L_2$ | 2 | (78, 171) | 0.02 |
| | δCCID.$L_\infty$ | 3 | (78, 126, 171) | 0.01 |
| | δCCID.$L_2$IC | 2 | (78, 171) | 0.05 |
| | δCCID.$L_\infty$IC | 3 | (78, 126, 171) | 0.02 |
| 7 | SBS | 0 | - | 1.42 |
|  | Factor com | 1 | 78 | 3.23 |
|  | Factor id | 1 | 78 | 3.23 |
|  | BO | 0 | - | 13.56 |
| | CCID.$L_2$ | 7 | (60, 63, 76, 92, 150, 164, 165) | 0.01 |
| | CCID.$L_\infty$ | 1 | 76 | 0.03 |
| | CCID.$L_2$IC | 2 | (60, 76) | 0.03 |
| | CCID.$L_\infty$IC | 2 | (64, 76) | 0.01 |
| | δCCID.$L_2$ | 2 | (76, 150) | 0.02 |
| | δCCID.$L_\infty$ | 1 | 76 | 0.01 |
| | δCCID.$L_2$IC | 1 | 76 | 0.05 |
| | δCCID.$L_\infty$IC | 1 | 76 | 0.02 |

| ID | Method | Number | Estimated<br>Locations | Time (s) |
| --- | --- | --- | --- | --- |
| 8 | SBS | 0 | - | 1.40 |
|  | Factor com | 0 | - | 3.38 |
|  | Factor id | 0 | - | 3.38 |
|  | BO | 2 | (102, 145) | 5.37 |
| | CCID.$L_2$ | 9 | (35, 38, 63, 70, 98, 112, 130, 185, 195) | 0.02 |
| | CCID.$L_\infty$ | 4 | (35, 38, 65, 129) | 0.02 |
| | CCID.$L_2$IC | 12 | (35, 38, 63, 70, 98, 112, 130, 145, 169, 185, 196, 209) | 0.04 |
| | CCID.$L_\infty$IC | 4 | (35, 38, 63, 70) | 0.02 |
| | δCCID.$L_2$ | 3 | (35, 130, 195) | 0.02 |
| | δCCID.$L_\infty$ | 2 | (35, 129) | 0.01 |
| | δCCID.$L_2$IC | 3 | (35, 145, 196) | 0.05 |
| | δCCID.$L_\infty$IC | 2 | (35, 129) | 0.02 |
| 9 | SBS | 0 | - | 1.45 |
|  | Factor com | 2 | (59, 141) | 3.72 |
|  | Factor id | 0 | - | 3.72 |
|  | BO | 1 | 125 | 5.37 |
| | CCID.$L_2$ | 12 | (30, 33, 59, 75, 105, 108, 123, 136, 164, 165, 204, 207) | 0.01 |
| | CCID.$L_\infty$ | 4 | (60, 74, 125, 136) | 0.01 |
| | CCID.$L_2$IC | 4 | (59, 75, 123, 136) | 0.04 |
| | CCID.$L_\infty$IC | 4 | (60,74,125,136) | 0.01 |
| | δCCID.$L_2$ | 3 | (59, 136, 204) | 0.02 |
| | δCCID.$L_\infty$ | 2 | (60, 136) | 0.01 |
| | δCCID.$L_2$IC | 2 | (59, 136) | 0.05 |
| | δCCID.$L_\infty$IC | 2 | (60, 136) | 0.02 |
| 10 | SBS | 0 | - | 1.58 |
|  | Factor com | 0 | - | 3.34 |
|  | Factor id | 0 | - | 3.34 |
|  | BO | 2 | (61,67) | 12.51 |
| | CCID.$L_2$ | 5 | (9, 60, 63, 71, 139) | 0.01 |
| | CCID.$L_\infty$ | 1 | 71 | 0.02 |
| | CCID.$L_2$IC | 3 | (60,63,71) | 0.03 |
| | CCID.$L_\infty$IC | 0 | - | 0.03 |
| | δCCID.$L_2$ | 3 | (9, 71, 139) | 0.02 |
| | δCCID.$L_\infty$ | 1 | 71 | 0.01 |
| | δCCID.$L_2$IC | 1 | 71 | 0.05 |
| | δCCID.$L_\infty$IC | 0 | - | 0.02 |
| 11 | SBS | 0 | - | 1.52 |
|  | Factor com | 0 | - | 4.05 |
|  | Factor id | 0 | - | 4.05 |
|  | BO | 0 | - | 5.12 |
| | CCID.$L_2$ | 7 | (31, 59, 88, 127, 146, 164, 210) | 0.01 |
| | CCID.$L_\infty$ | 2 | (76, 160) | 0.02 |
| | CCID.$L_2$IC | 3 | (59, 146, 169) | 0.03 |
| | CCID.$L_\infty$IC | 1 | 160 | 0.02 |
| | δCCID.$L_2$ | 3 | (31, 164, 210) | 0.02 |
| | δCCID.$L_\infty$ | 2 | 76, 160 | 0.01 |
| | δCCID.$L_2$IC | 2 | 59, 169 | 0.05 |
| | δCCID.$L_\infty$IC | 1 | 160 | 0.02 |
| 12 | SBS | 1 | 109 | 1.49 |
|  | Factor com | 1 | 96 | 2.80 |
|  | Factor id | 0 | - | 2.80 |
|  | BO | 0 | - | 10.26 |
| | CCID.$L_2$ | 11 | (2, 49, 69, 70, 94, 114, 128, 149, 185, 195, 201) | 0.02 |
| | CCID.$L_\infty$ | 5 | (2, 59, 66, 97, 195) | 0.02 |
| | CCID.$L_2$IC | 5 | (2, 49, 114, 167, 193) | 0.05 |
| | CCID.$L_\infty$IC | 3 | (2, 59, 114) | 0.03 |
| | δCCID.$L_2$ | 4 | (2, 49, 114, 195) | 0.02 |
| | δCCID.$L_\infty$ | 3 | (2, 66, 195) | 0.01 |
| | δCCID.$L_2$IC | 4 | (2, 49, 114, 193) | 0.05 |
| | δCCID.$L_\infty$IC | 3 | (2, 59, 114) | 0.02 |
| 13 | SBS | 0 | - | 1.42 |
|  | Factor com | 0 | - | 2.83 |
|  | Factor id | 0 | - | 2.83 |
|  | BO | 0 | - | 5.17 |
| | CCID.$L_2$ | 4 | (7, 112, 114, 195) | 0.02 |
| | CCID.$L_\infty$ | 0 | - | 0.02 |
| | CCID.$L_2$IC | 0 | - | 0.02 |
| | CCID.$L_\infty$IC | 0 | - | 0.02 |
| | δCCID.$L_2$ | 3 | (7, 112, 195) | 0.02 |
| | δCCID.$L_\infty$ | 0 | - | 0.01 |
| | δCCID.$L_2$IC | 0 | - | 0.05 |
| | δCCID.$L_\infty$IC | 0 | - | 0.02 |
| 14 | SBS | 1 | 155 | 1.40 |
|  | Factor com | 1 | 151 | 3.20 |
|  | Factor id | 1 | 168 | 3.20 |
|  | BO | 1 | 153 | 10.29 |
| | CCID.$L_2$ | 8 | (12, 16, 68, 70, 74, 125, 198, 202) | 0.02 |
| | CCID.$L_\infty$ | 2 | (150, 192) | 0.04 |
| | CCID.$L_2$IC | 4 | (92, 125, 202, 207) | 0.03 |
| | CCID.$L_\infty$IC | 2 | (134, 192) | 0.03 |
| | δCCID.$L_2$ | 3 | (12, 74, 125, 198) | 0.02 |
| | δCCID.$L_\infty$ | 2 | (150, 192) | 0.01 |
| | δCCID.$L_2$IC | 2 | (125, 202) | 0.05 |
| | δCCID.$L_\infty$IC | 2 | (134, 192) | 0.02 |
| 15 | SBS | 1 | 138 | 1.61 |
|  | Factor com | 3 | (31, 111, 138) | 3.10 |
|  | Factor id | 0 | - | 3.10 |
|  | BO | 1 | 134 | 10.03 |
| | CCID.$L_2$ | 6 | (60, 63, 75, 110, 131, 138) | 0.02 |
| | CCID.$L_\infty$ | 1 | 137 | 0.01 |
| | CCID.$L_2$IC | 3 | (75, 110, 155) | 0.03 |
| | CCID.$L_\infty$IC | 1 | 137 | 0.01 |
| | δCCID.$L_2$ | 1 | 138 | 0.02 |
| | δCCID.$L_\infty$ | 1 | 137 | 0.01 |
| | δCCID.$L_2$IC | 2 | (110, 155) | 0.05 |
| | δCCID.$L_\infty$IC | 1 | 137 | 0.02 |
| 16 | SBS | 0 | - | 1.47 |
|  | Factor com | 0 | - | 2.98 |
|  | Factor id | 0 | - | 2.98 |
|  | BO | 0 | - | 10.12 |
| | CCID.$L_2$ | 16 | (14, 25, 59, 79, 105, 108, 117, 130, 132, 138, 184, 185, 195, 196, 198, 205) | 0.01 |
| | CCID.$L_\infty$ | 5 | (65, 68, 108, 196, 205) | 0.01 |
| | CCID.$L_2$IC | 10 | (25, 59, 90, 93, 105, 108, 138, 195, 198, 205) | 0.07 |
| | CCID.$L_\infty$IC | 2 | (65, 108) | 0.02 |
| | δCCID.$L_2$ | 3 | (59, 138, 205) | 0.02 |
| | δCCID.$L_\infty$ | 3 | (65, 108, 205) | 0.01 |
| | δCCID.$L_2$IC | 3 | (59, 108, 195) | 0.05 |
| | δCCID.$L_\infty$IC | 2 | (65, 108) | 0.02 |
| 17 | SBS | 0 | - | 1.45 |
|  | Factor com | 1 | 188 | 3.09 |
|  | Factor id | 1 | 187 | 3.09 |
|  | BO | 2 | (111,190) | 14.48 |
| | CCID.$L_2$ | 10 | (9, 10, 63, 74, 108, 109, 156, 187, 189, 195) | 0.02 |
| | CCID.$L_\infty$ | 6 | (62, 70, 125, 154, 187, 189) | 0.01 |
| | CCID.$L_2$IC | 12 | (9, 10, 63, 74, 108, 109, 126, 156, 161, 187, 195, 210) | 0.03 |
| | CCID.$L_\infty$IC | 4 | (62, 70, 154, 187) | 0.03 |
| | δCCID.$L_2$ | 3 | (9, 74, 156) | 0.02 |
| | δCCID.$L_\infty$ | 2 | (62, 154) | 0.01 |
| | δCCID.$L_2$IC | 3 | (9,74, 156) | 0.05 |
| | δCCID.$L_\infty$IC | 2 | (62, 154) | 0.02 |
| 18 | SBS | 0 | - | 1.44 |
|  | Factor com | 1 | 75 | 2.83 |
|  | Factor id | 0 | - | 2.83 |
|  | BO | 2 | (46, 186) | 13.86 |
| | CCID.$L_2$ | 6 | (83, 125, 165, 167, 175, 179) | 0.02 |
| | CCID.$L_\infty$ | 1 | 145 | 0.03 |
| | CCID.$L_2$IC | 1 | 179 | 0.02 |
| | CCID.$L_\infty$IC | 1 | 184 | 0.02 |
| | δCCID.$L_2$ | 3 | (59, 138, 205) | 0.02 |
| | δCCID.$L_\infty$ | 3 | (65, 108, 205) | 0.01 |
| | δCCID.$L_2$IC | 1 | 179 | 0.05 |
| | δCCID.$L_\infty$IC | 1 | 184 | 0.02 |
| 19 | SBS | 0 | - | 1.42 |
|  | Factor com | 0 | - | 3.25 |
|  | Factor id | 0 | - | 3.25 |
|  | BO | 0 | - | 5.26 |
| | CCID.$L_2$ | 8 | (61, 71, 125, 130, 142, 186, 194, 195) | 0.02 |
| | CCID.$L_\infty$ | 4 | (56, 75, 130, 141) | 0.01 |
| | CCID.$L_2$IC | 8 | (61, 71, 125, 130, 134, 142, 177, 209) | 0.03 |
| | CCID.$L_\infty$IC | 3 | (75, 130, 141) | 0.02 |
| | δCCID.$L_2$ | 3 | (71, 142, 186) | 0.02 |
| | δCCID.$L_\infty$ | 2 | (75, 141) | 0.01 |
| | δCCID.$L_2$IC | 3 | (59, 108, 195) | 0.05 |
| | δCCID.$L_\infty$IC | 2 | (71, 142) | 0.02 |
| 20 | SBS | 0 | - | 1.43 |
|  | Factor com | 0 | - | 3.45 |
|  | Factor id | 0 | - | 3.45 |
|  | BO | 0 | - | 5.12 |
| | CCID.$L_2$ | 5 | (96, 101, 104, 139, 140) | 0.01 |
| | CCID.$L_\infty$ | 1 | 104 | 0.02 |
| | CCID.$L_2$IC | 0 | - | 0.03 |
| | CCID.$L_\infty$IC | 0 | - | 0.01 |
| | δCCID.$L_2$ | 1 | 104 | 0.02 |
| | δCCID.$L_\infty$ | 1 | 104 | 0.01 |
| | δCCID.$L_2$IC | 0 | - | 0.05 |
| | δCCID.$L_\infty$IC | 0 | - | 0.02 |
| 21 | SBS | 0 | - | 1.42 |
|  | Factor com | 0 | - | 2.31 |
|  | Factor id | 0 | - | 2.31 |
|  | BO | 0 | - | 5.18 |
| | CCID.$L_2$ | 12 | (7, 47, 62, 97, 100, 112, 115, 126, 146, 160, 169, 183) | 0.01 |
| | CCID.$L_\infty$ | 2 | (7, 160) | 0.02 |
| | CCID.$L_2$IC | 11 | (21, 47, 62, 98, 100, 112, 115, 146, 160, 169, 183) | 0.05 |
| | CCID.$L_\infty$IC | 3 | (7, 160, 183) | 0.03 |
| | δCCID.$L_2$ | 4 | (7, 62, 115, 183) | 0.02 |
| | δCCID.$L_\infty$ | 2 | (7, 160) | 0.01 |
| | δCCID.$L_2$IC | 3 | (62, 125, 183) | 0.05 |
| | δCCID.$L_\infty$IC | 2 | (7, 183) | 0.02 |
| 22 | SBS | 0 | - | 1.40 |
|  | Factor com | 0 | - | 2.36 |
|  | Factor id | 0 | - | 2.36 |
|  | BO | 0 | - | 5.10 |
| | CCID.$L_2$ | 7 | (61, 63, 101, 105, 130, 179, 209) | 0.01 |
| | CCID.$L_\infty$ | 2 | (73, 135) | 0.03 |
| | CCID.$L_2$IC | 8 | (60, 71, 101, 125, 150, 176, 179, 209) | 0.04 |
| | CCID.$L_\infty$IC | 2 | (73, 135) | 0.02 |
| | δCCID.$L_2$ | 2 | (61, 179) | 0.02 |
| | δCCID.$L_\infty$ | 2 | (73, 135) | 0.01 |
| | δCCID.$L_2$IC | 3 | (60, 101, 179) | 0.05 |
| | δCCID.$L_\infty$IC | 2 | (73, 135) | 0.02 |
| 23 | SBS | 0 | - | 1.44 |
|  | Factor com | 1 | 174 | 3.11 |
|  | Factor id | 0 | - | 3.11 |
|  | BO | 0 | - | 5.22 |
| | CCID.$L_2$ | 8 | (60, 69, 128, 145, 155, 176, 194, 207) | 0.02 |
| | CCID.$L_\infty$ | 1 | 173 | 0.01 |
| | CCID.$L_2$IC | 5 | (60, 69, 173, 194, 207) | 0.06 |
| | CCID.$L_\infty$IC | 1 | 173 | 0.03 |
| | δCCID.$L_2$ | 2 | (60, 145) | 0.02 |
| | δCCID.$L_\infty$ | 1 | 173 | 0.01 |
| | δCCID.$L_2$IC | 2 | (60, 173) | 0.05 |
| | δCCID.$L_\infty$IC | 1 | 173 | 0.02 |

**Table 12:** The estimated number and location of the change-points for the resting state fMRI data set for Sparsified Binary Segmentation (SBS), Factor with common components (Factor com), Factor with idiosyncratic components (Factor id), Barnett & Onnela (2016) (BO), and Cross-covariance isolate detect with the *L*_2_ threshold (CCID.*L*_2_), the *L*_∞_ threshold (CCID.*L*_∞_), the *L*_2_ information criterion (CCID.*L*_2_*IC*) and the *L*_∞_ information criterion (CCID.*L*_∞_*IC*). A *δ* in the name of a method indicates a minimum distance between change-points of 40 time points was enforced. The computational times for each method are also provided.

| ID | Method | Number | Estimated Locations | Time (s) |
| --- | --- | --- | --- | --- |
| 1 | SBS | 1 | 72 | 142.90 |
|  | Factor com | 0 | - | 91.03 |
|  | Factor id | 0 | - | 91.03 |
|  | BO | 9 | (11, 35, 66, 83, 116, 139, 178, 193, 211) | 352.57 |
| | CCID.$L_2$ | 15 | (22, 41, 73, 85, 90, 124, 134, 145, 190, 198, 206, 217, 257, 264, 275) | 9.23 |
| | CCID.$L_\infty$ | 30 | (18, 31, 51, 72, 74, 78, 86, 118, 121, 123, 125, 129, 131, 134, 138, 146, 181, 186, 188, 190, 193, 195, 198, 201, 207, 228, 256, 260, 262, 266) | 17.42 |
| | CCID.$L_2IC$ | 17 | (14, 25, 38, 52, 64, 73, 90, 108, 124, 132, 146, 169, 194, 222, 236, 253, 276) | 56.42 |
| | CCID.$L_\infty IC$ | 13 | (18, 44, 64, 97, 118, 138, 158, 181, 197, 216, 236, 254, 266) | 54.69 |
| | δCCID.$L_2$ | 4 | (90, 145, 190, 257) | 0.02 |
| | δCCID.$L_\infty$ | 4 | (18, 86, 146, 256) | 0.01 |
| | δCCID.$L_2IC$ | 5 | (38, 108, 194, 236, 276) | 0.05 |
| | δCCID.$L_\infty IC$ | 5 | (18, 64, 138, 181, 254) | 0.02 |
| 2 | SBS | 1 | 88 | 119.46 |
|  | Factor com | 0 | - | 92.24 |
|  | Factor id | 0 | - | 92.24 |
|  | BO | 8 | (47, 57, 115, 176, 193, 220, 232, 255) | 373.02 |
| | CCID.$L_2$ | 14 | (2, 47, 59, 74, 85, 96, 125, 154, 182, 196, 244, 252, 274, 282) | 17.69 |
| | CCID.$L_\infty$ | 22 | (6, 8, 28, 47, 74, 88, 90, 92, 97, 100, 123, 125, 157, 188, 196, 218, 232, 234, 236, 257, 274, 276) | 17.30 |
| | CCID.$L_2IC$ | 8 | (13, 47, 59, 86, 96, 154, 206, 235) | 83.03 |
| | CCID.$L_\infty IC$ | 7 | (9, 47, 74, 88, 148, 236, 274) | 83.69 |
| | δCCID.$L_2$ | 5 | (2, 47, 96, 154, 274) | 0.02 |
| | δCCID.$L_\infty$ | 4 | (8, 125, 188, 274) | 0.01 |
| | δCCID.$L_2IC$ | 4 | (47, 96, 154, 235) | 0.05 |
| | δCCID.$L_\infty IC$ | 4 | (9, 88, 148, 274) | 0.02 |
| 3 | SBS | 2 | (117, 145) | 121.60 |
|  | Factor com | 1 | 40 | 90.54 |
|  | Factor id | 0 | - | 90.54 |
|  | BO | 9 | (89, 122, 153, 164, 179, 207, 220, 249, 261) | 369.08 |
| | CCID.$L_2$ | 12 | (2, 8, 18, 65, 93, 100, 115, 166, 225, 251, 262, 278) | 13.96 |
| | CCID.$L_\infty$ | 18 | (18, 37, 57, 87, 117, 119, 154, 182, 186, 188, 190, 193, 218, 221, 225, 229, 247, 266) | 16.83 |
| | CCID.$L_2IC$ | 11 | (8, 47, 81, 101, 116, 158, 166, 200, 225, 250, 278) | 49.96 |
| | CCID.$L_\infty IC$ | 3 | (18, 64, 247) | 49.69 |
| | δCCID.$L_2$ | 4 | (18, 115, 166, 251) | 0.02 |
| | δCCID.$L_\infty$ | 3 | (18, 87, 218) | 0.01 |
| | δCCID.$L_2IC$ | 4 | (8, 116, 166, 278) | 0.05 |
| | δCCID.$L_\infty IC$ | 3 | (18, 64, 247) | 0.02 |

| ID | Method | Number | Estimated<br>Locations | Time (s) |
| --- | --- | --- | --- | --- |
| 4 | SBS | 1 | 173 | 114 |
|  | Factor com | 0 | - | 131.67 |
|  | Factor id | 0 | - | 131.67 |
|  | BO | 1 | 229 | 210.41 |
| | CCID.$L_2$ | 9 | (2, 28, 46, 73, 81, 153, 231, 242, 262) | 14.57 |
| | CCID.$L_\infty$ | 22 | (2, 27, 36, 43, 46, 50, 68, 97, 99, 135, 148, 172, 195, 197, 227, 229, 252, 256, 258, 261, 263, 266) | 18.58 |
| | CCID.$L_2IC$ | 11 | (8, 27, 46, 73, 81, 111, 153, 190, 231, 242, 275) | 44.10 |
| | CCID.$L_\infty IC$ | 3 | (7, 186, 263) | 115.88 |
| | δCCID.$L_2$ | 4 | (2, 81, 153, 231) | 0.02 |
| | δCCID.$L_\infty$ | 3 | (2, 197, 252) | 0.01 |
| | δCCID.$L_2IC$ | 5 | (8, 111, 190, 231, 275) | 0.05 |
| | δCCID.$L_\infty IC$ | 3 | (7, 186, 263) | 0.02 |
| 5 | SBS | 1 | 189 | 115.89 |
|  | Factor com | 1 | 178 | 159.23 |
|  | Factor id | 0 | - | 159.23 |
|  | BO | 10 | (30, 63, 84, 107, 141, 184, 204, 235, 248, 263) | 440.72 |
| | CCID.$L_2$ | 9 | (6, 20, 65, 95, 115, 172, 221, 231, 269) | 10.61 |
| | CCID.$L_\infty$ | 19 | (6, 12, 22, 42, 67, 70, 88, 110, 112, 127, 158, 177, 180, 182, 207, 234, 236, 257, 278) | 13.07 |
| | CCID.$L_2IC$ | 7 | (6, 74, 117, 172, 197, 207, 267) | 29.89 |
| | CCID.$L_\infty IC$ | 8 | (22, 68, 107, 112, 159, 182, 224, 263) | 49.26 |
| | δCCID.$L_2$ | 4 | (20, 95, 172, 221) | 0.02 |
| | δCCID.$L_\infty$ | 4 | (67, 127, 207, 278) | 0.01 |
| | δCCID.$L_2IC$ | 5 | (6, 74, 117, 172, 267) | 0.05 |
| | δCCID.$L_\infty IC$ | 4 | (22, 112, 159, 224) | 0.02 |
| 6 | SBS | 2 | (116, 208) | 111.42 |
|  | Factor com | 1 | 94 | 139.20 |
|  | Factor id | 0 | - | 139.20 |
|  | BO | 10 | (38, 51, 62, 90, 128, 155, 182, 200, 226, 237) | 372.46 |
| | CCID.$L_2$ | 13 | (17, 56, 89, 99, 112, 125, 144, 157, 173, 196, 207, 227, 267) | 12.01 |
| | CCID.$L_\infty$ | 22 | (25, 57, 87, 93, 97, 100, 102, 104, 113, 116, 137, 167, 187, 208, 211, 237, 257, 260, 264, 268, 274, 277) | 23.86 |
| | CCID.$L_2IC$ | 8 | (41, 90, 99, 144, 203, 210, 239, 266) | 67.78 |
| | CCID.$L_\infty IC$ | 9 | (25, 57, 89, 93, 113, 182, 207, 257, 277) | 97.33 |
| | δCCID.$L_2$ | 5 | (56, 125, 173, 227, 267) | 0.02 |
| | δCCID.$L_\infty$ | 5 | (25, 113, 167, 211, 257) | 0.01 |
| | δCCID.$L_2IC$ | 5 | (41, 90, 144, 210, 266) | 0.05 |
| | δCCID.$L_\infty IC$ | 4 | (25, 89, 182, 277) | 0.02 |
| 7 | SBS | 2 | (102, 151) | 121.45 |
|  | Factor com | 0 | - | 142.64 |
|  | Factor id | 0 | - | 142.64 |
|  | BO | 8 | (50, 66, 103, 115, 139, 169, 239, 264) | 396.44 |
| | CCID.$L_2$ | 9 | (2, 61, 70, 117, 173, 207, 234, 245, 275) | 13.94 |
| | CCID.$L_\infty$ | 16 | (18, 21, 47, 71, 88, 114, 128, 157, 169, 194, 219, 237, 257) | 16.82 |
| | CCID.$L_2IC$ | 1 | 85 | 37.50 |
| | CCID.$L_\infty IC$ | 4 | (88, 113, 194, 219) | 37.47 |
| | δCCID.$L_2$ | 4 | (2, 70, 117, 245) | 0.02 |
| | δCCID.$L_\infty$ | 5 | (18, 88, 128, 219, 265) | 0.01 |
| | δCCID.$L_2IC$ | 1 | 85 | 0.05 |
| | δCCID.$L_\infty IC$ | 2 | (113, 219) | 0.02 |
| 8 | SBS | 1 | 130 | 121.22 |
|  | Factor com | 0 | - | 171.89 |
|  | Factor id | 0 | - | 171.89 |
|  | BO | 12 | (12, 36, 61, 84, 96, 121, 131, 192, 221, 238, 251, 265) | 409.98 |
| | CCID.$L_2$ | 14 | (15, 53, 87, 98, 126, 134, 141, 150, 167, 190, 206, 235, 264, 282) | 9.83 |
| | CCID.$L_\infty$ | 23 | (2, 24, 53, 68, 71, 73, 98, 116, 138, 141, 144, 147, 151, 173, 188, 206, 225, 227, 259, 261, 264, 273, 278) | 20.29 |
| | CCID.$L_2IC$ | 7 | (126, 150, 186, 203, 214, 234, 264) | 36.68 |
| | CCID.$L_\infty IC$ | 4 | (42, 196, 237, 278) | 84.21 |
| | δCCID.$L_2$ | 4 | (53, 98, 190, 235) | 0.02 |
| | δCCID.$L_\infty$ | 3 | (2, 53, 188) | 0.01 |
| | δCCID.$L_2IC$ | 2 | (126, 234) | 0.05 |
| | δCCID.$L_\infty IC$ | 4 | (42, 196, 237, 278) | 0.02 |
| 9 | SBS | 1 | 212 | 110.36 |
|  | Factor com | 1 | 236 | 139.03 |
|  | Factor id | 0 | - | 139.03 |
|  | BO | 10 | (34, 59, 81, 124, 147, 168, 200, 215, 244, 257) | 365.12 |
| | CCID.$L_2$ | 8 | (68, 79, 145, 189, 201, 235, 246, 270) | 11.36 |
| | CCID.$L_\infty$ | 19 | (8, 11, 35, 57, 60, 87, 113, 134, 137, 142, 172, 188, 217, 234, 246, 269, 271, 273, 277) | 15.39 |
| | CCID.$L_2IC$ | 9 | (29, 42, 68, 87, 127, 171, 189, 215, 264) | 33.94 |
| | CCID.$L_\infty IC$ | 5 | (51, 86, 177, 225, 269) | 57.17 |
| | δCCID.$L_2$ | 3 | (79, 145, 189) | 0.02 |
| | δCCID.$L_\infty$ | 4 | (11, 87, 142, 246) | 0.01 |
| | δCCID.$L_2IC$ | 6 | (29, 87, 127, 171, 215, 264) | 0.05 |
| | δCCID.$L_\infty IC$ | 4 | (86, 177, 225, 269) | 0.02 |
| 10 | SBS | 1 | 202 | 109.35 |
|  | Factor com | 0 | - | 178.42 |
|  | Factor id | 0 | - | 178.42 |
|  | BO | 11 | (33, 69, 97, 112, 129, 149, 161, 193, 211, 237, 249) | 363.08 |
| | CCID.$L_2$ | 12 | (2, 42, 65, 97, 108, 149, 166, 200, 215, 235, 251, 282) | 11.47 |
| | CCID.$L_\infty$ | 28 | (17, 43, 45, 67, 74, 77, 97, 117, 119, 148, 159, 162, 164, 167, 170, 172, 186, 188, 198, 200, 202, 205, 207, 233, 235, 237, 256, 282) | 16.06 |
| | CCID.$L_2IC$ | 10 | (17, 41, 66, 82, 117, 144, 166, 234, 251, 276) | 42.29 |
| | CCID.$L_\infty IC$ | 12 | (17, 38, 58, 65, 107, 119, 138, 157, 188, 220, 256, 282) | 101.19 |
| | δCCID.$L_2$ | 5 | (2, 42, 108, 215, 282) | 0.02 |
| | δCCID.$L_\infty$ | 5 | (17, 67, 117, 186, 282) | 0.01 |
| | δCCID.$L_2IC$ | 5 | (17, 82, 144, 234, 276) | 0.05 |
| | δCCID.$L_\infty IC$ | 5 | (17, 58, 119, 188, 282) | 0.02 |

**Table 13:** The distribution of 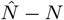 and the scaled Hausdorff distance over 100 simulated data sequences from Simulations 1, 3, and 5, with the noise following the *t*_5_ distribution, for the Cross-covariance isolate detect with the *L*_2_ threshold (CCID.*L*_2_), the *L*_∞_ threshold (CCID.*L*_∞_), the *L*_2_ information criterion (CCID.*L*_2_*IC*) and the *L*_∞_ information criterion (CCID.*L*_∞_*IC*). The computational times are also provided.

| Simulation data | Method | $\hat{N} - N$ | | | | | | | $d_H$ | Time (s) |
| --- | --- | --- | --- | --- | --- | --- | --- | --- | --- | --- |
| | | $\leq -3$ | -2 | -1 | 0 | 1 | 2 | $\geq 3$ | | |
| 1 | SBS | - | - | - | 100 | 0 | 0 | 0 | - | 2.82 |
|  | Factor com | - | - | - | 100 | 0 | 0 | 0 | - | 16.62 |
|  | Factor id | - | - | - | 95 | 5 | 0 | 0 | - | 16.62 |
|  | BO | - | - | - | 85 | 15 | 0 | 0 | - | 7.66 |
| | CCID.$L_2$ | - | - | - | 39 | 21 | 25 | 15 | - | 0.28 |
| | CCID.$L_\infty$ | - | - | - | 28 | 23 | 21 | 28 | - | 0.36 |
| | CCID.$L_2IC$ | - | - | - | 97 | 1 | 2 | 0 | - | 0.42 |
| | CCID.$L_\infty IC$ | - | - | - | 98 | 1 | 1 | 0 | - | 0.39 |
| 3 | SBS | 5 | 5 | 16 | 74 | 0 | 0 | 0 | 0.50 | 3.90 |
|  | Factor com | 100 | 0 | 0 | 0 | 0 | 0 | 0 | 4 | 18.22 |
|  | Factor id | 73 | 11 | 10 | 6 | 0 | 0 | 0 | 2.96 | 18.22 |
|  | BO | 3 | 2 | 13 | 57 | 22 | 3 | 0 | 0.55 | 39.87 |
| | CCID.$L_2$ | 0 | 0 | 12 | 59 | 21 | 6 | 2 | 0.31 | 0.17 |
| | CCID.$L_\infty$ | 0 | 0 | 4 | 44 | 30 | 16 | 6 | 0.35 | 0.18 |
| | CCID.$L_2IC$ | 1 | 1 | 12 | 74 | 7 | 5 | 0 | 0.31 | 0.56 |
| | CCID.$L_\infty IC$ | 1 | 5 | 11 | 76 | 3 | 4 | 0 | 0.37 | 0.32 |
| 5 | SBS | 69 | 26 | 1 | 4 | 0 | 0 | 0 | 3.03 | 3.95 |
|  | Factor com | 100 | 0 | 0 | 0 | 0 | 0 | 0 | 4 | 19.32 |
|  | Factor id | 16 | 11 | 2 | 71 | 0 | 0 | 0 | 0.89 | 19.32 |
|  | BO | 60 | 21 | 8 | 6 | 4 | 1 | 0 | 2.89 | 24.26 |
| | CCID.$L_2$ | 0 | 0 | 0 | 57 | 24 | 12 | 7 | 0.21 | 0.18 |
| | CCID.$L_\infty$ | 0 | 2 | 6 | 37 | 21 | 13 | 21 | 0.45 | 0.21 |
| | CCID.$L_2IC$ | 15 | 10 | 2 | 53 | 7 | 10 | 3 | 0.94 | 0.65 |
| | CCID.$L_\infty IC$ | 24 | 19 | 1 | 44 | 5 | 6 | 1 | 1.38 | 0.46 |

## B Figures showing the results of SBS, Factor and BO for the fMRI data sets

**Figure 14:**
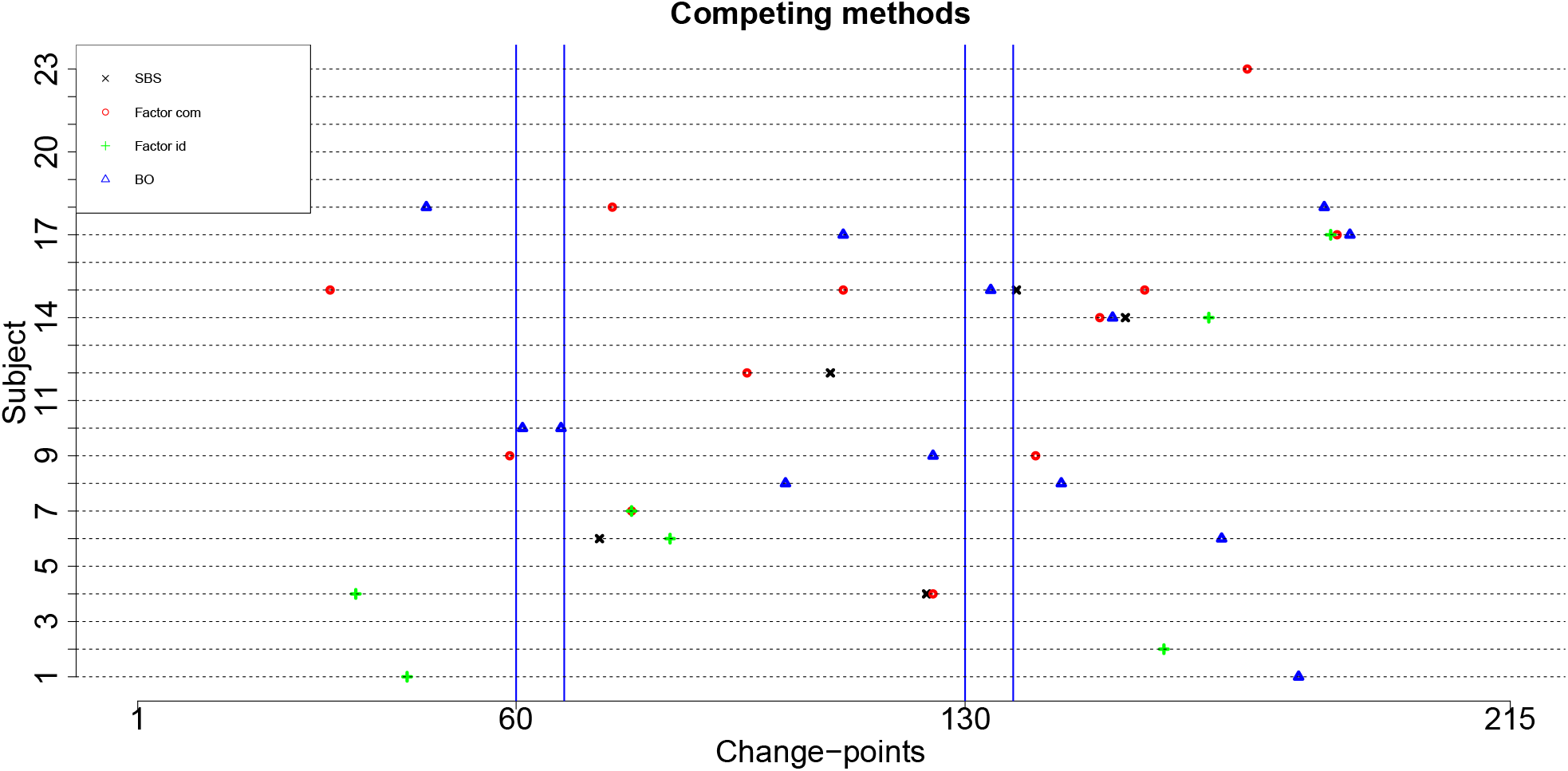
The detected change-points for the 4 ROI and heart rate data set using SBS, Factor, and BO. The y-axis depicts the subject number while the x-axis shows the change-point location times. The blue vertical lines indicate the times of the showing and of the removal of the visual cues.

**Figure 15:**
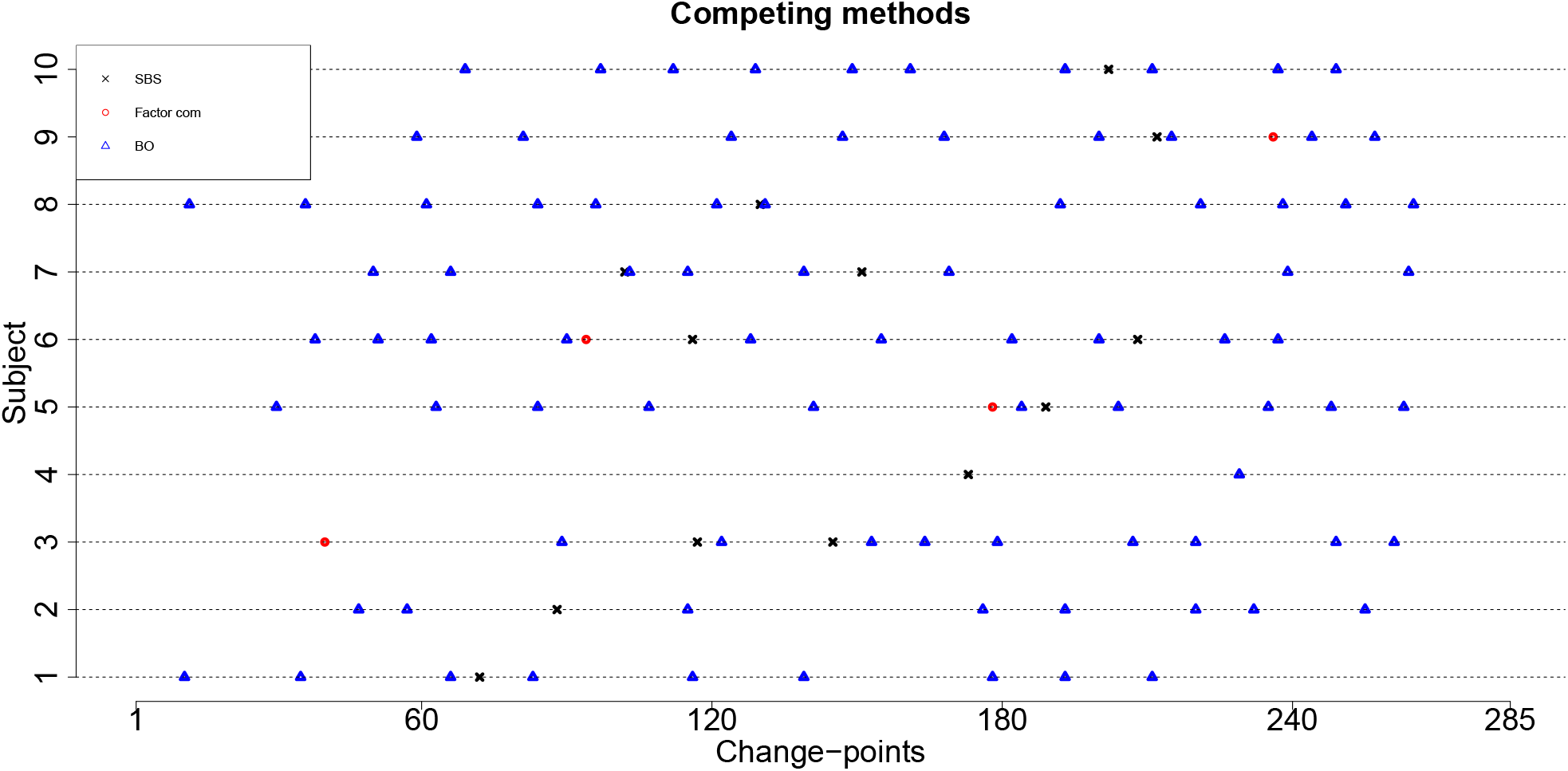
The detected change-points for the resting state fMRI data set using SBS, Factor com, and BO. The results for Factor idiosyncratic are not presented because this method does not detect any change-points for all subjects. The y-axis depicts the subject number while the x-axis shows the change-point location times.

